# Chronic Circadian Disturbance Reshapes Hippocampal Connectivity and Cognitive Function

**DOI:** 10.64898/2025.12.27.696650

**Authors:** Inês Marques-Morgado, Marcelo Dias, Joana E. Coelho, Carolina Peixoto, Mariana Temido-Ferreira, Joana Gomes-Ribeiro, Miguel Remondes, Neil Dawson, Luísa V. Lopes

**Author notes:** Correspondence to: Luísa V. Lopes Gulbenkian Institute for Molecular Medicine Av Prof Egas Moniz 1649-028 Lisbon.

## Abstract

Circadian rhythms regulate a wide range of physiological and cognitive functions, yet the neural mechanisms linking circadian timing to memory remain poorly defined. Here, we identify a direct anatomical projection from the suprachiasmatic nucleus (SCN) to the dorsal hippocampus using complementary viral anterograde, retrograde, and monosynaptic tracing approaches, alongside established indirect pathways via septal and neuromodulatory regions. These findings revise prevailing models that restrict SCN influence on hippocampal function to polysynaptic routes.

Functionally, chronic circadian disruption (CCD) induced region-specific metabolic alterations and widespread reorganization of brain-wide functional connectivity (FC). Hypometabolism emerged in the SCN, dentate gyrus, and perirhinal and entorhinal cortices, accompanied by loss of SCN– hippocampal connectivity and compensatory engagement of cortical and subcortical networks. In contrast, the serotonergic dorsal raphe nucleus showed selective hypermetabolism and increased connectivity with the SCN, prefrontal cortex, and hippocampus, suggesting altered neuromodulatory control under circadian misalignment.

Behaviorally, CCD produced a selective impairment in object recognition memory, while spatial memory, working memory, cognitive flexibility, and hippocampal synaptic plasticity at the Schaffer collateral–CA1 synapse remained intact. Disrupted functional coupling within perirhinal–entorhinal– hippocampal and prefrontal–hippocampal circuits, together with altered SCN–medial septum– hippocampus connectivity, indicates that cognitive dysfunction arises from desynchronization of distributed brain networks rather than local synaptic failure.

Together, these findings provide a new mechanistic framework linking circadian disruption to selective memory impairment and highlight network-level synchrony as a potential target for chronotherapeutic intervention.

## Introduction

In mammals, a ∼24-hour periodic circadian rhythm orchestrates nearly every physiological and behavioral process, including sleep-wake cycles, hormonal secretion, body temperature, metabolism, and cognitive function (Foster & Kreitzman, 2017; Kumar, 2017). These rhythms are driven by a master pacemaker located in the suprachiasmatic nucleus (SCN) of the anterior hypothalamus. The SCN synchronizes peripheral clocks throughout the body via neural and hormonal signals, ensuring systemic temporal coherence (Dibner et al., 2010).

A key adaptive advantage of circadian organization is the temporal separation of incompatible biological processes. Such temporal structuring is critical for maintaining internal homeostasis while optimizing responses to external cues and stresses (Golombek & Rosenstein, 2010). Consequently, circadian disruption — whether due to genetic mutation, lifestyle (e.g., shift work) or artificial light exposure — has been implicated in a wide array of disorders, including sleep disturbances, metabolic syndrome, mood disorders and cognitive decline (Karatsoreos, 2012).

There is growing evidence that circadian rhythms play an essential role in modulating neural activity in brain regions involved in learning and memory, particularly the hippocampus (Smarr et al., 2014a). Fluctuations in hippocampal excitability, plasticity, and gene expression have been shown to follow circadian patterns, suggesting that optimal memory formation and retrieval occur at specific times of day (Ruby et al., 2008). Indeed, electrophysiological studies reveal circadian modulation of long-term potentiation (LTP) and long-term depression (LTD), which are core mechanisms of synaptic plasticity and memory encoding (Chaudhury et al., 2005).

Despite the above evidence, the mechanisms and brain structures underlying the circadian regulation of memory functions, particularly those relating SCN fluctuations with neural activity in hippocampal circuits, remain poorly understood. While some studies suggest indirect, polysynaptic pathways involving hypothalamic and septal relays (Swanson and Cowen, 1977), other studies suggest the existence of direct structural connectivity between the SCN and hippocampus (Vrang et al, 1995). Direct SCN-hippocampal structural connectivity remains controversial, partly due to technical limitations in earlier tracing methods.

Chronic circadian disturbance (CCD), as experienced by shift workers or individuals in irregular light environments, has been associated with memory impairments and increased risk for neurodegenerative diseases (Logan & McClung, 2019). In rodent models, CCD results in hippocampal dendritic spine loss, reduced neurogenesis, and impaired cognition (Karatsoreos et al., 2011; Craig & McDonald, 2008). Interestingly, while some studies highlight broad cognitive deficits under CCD, others suggest a more selective impact, affecting specific memory domains or temporal windows (Smarr et al, 2014a.b). Moreover, the disturbed neurocircuitry contributing to these deficits is not yet defined. Thus, there is a need for high-resolution anatomical, physiological and behavioral investigations to clarify how circadian networks interact with memory systems, and how these are perturbed in CCD.

To address the gaps in our understanding of how circadian timing systems influence memory-related brain networks, we employed an integrative, multi-scale approach combining viral tracing, metabolic functional brain imaging, FC modeling, and behavioral assays. We first characterized the anatomical link between the suprachiasmatic nucleus (SCN) and the hippocampus using transsynaptic and monosynaptic tracers, revealing direct structural projections from the SCN to multiple hippocampal subfields. We then investigated how CCD impacts on brain-wide glucose metabolism, as a measure of regional neuronal activity, and inter-regional FC, uncovering region-specific alterations and the widespread reorganization of inter-regional coupling, particularly between circadian and cognitive hubs. Our behavioral assessments revealed a selective CCD-induced impairment in object recognition memory, with spatial and working memory function unaffected. Finally, *in vitro* electrophysiology showed that synaptic plasticity at the Schaffer collateral–CA1 synapse was preserved, suggesting that the observed memory deficits are not due to synaptic failures in this hippocampal circuit but may arise from disrupted brain network connectivity, including SCN-hippocampal dysconnectivity, as indicated by altered FC between neural systems.

Collectively, these findings shed light on a previously underappreciated SCN–hippocampus circuit and demonstrate how its functional integrity is compromised under chronic circadian misalignment, and its relationship with selective cognitive impairment and brain-wide circuit changes.

## Results

### Characterization of the Anatomical Connectivity between the SCN and Hippocampus

The suprachiasmatic nucleus (SCN) is the central pacemaker of circadian rhythms, yet its structural connectivity to memory-related regions, such as the hippocampus, remains poorly defined. To address this, we employed viral transsynaptic tracing and monosynaptic labeling tools to systematically define the anatomical connectivity between the SCN and the hippocampal formation.

#### SCN–Hippocampal Connectivity via Anterograde Transsynaptic Tracing

To delineate SCN efferents, we injected the anterograde transsynaptic tracer rVSV-(VSV-G)-Venus into the SCN of Sprague Dawley rats. At six days post-injection, robust labeling was observed in canonical first-order SCN targets, and associated regions of this circuitry, including the Medial Septum (MS), Lateral Septum (LS), Paraventricular Hypothalamic Nucleus (PVH), and Subparaventricular Zone (SPZ) (**Figure 1**). Consistent with prior work, destructive lesions were also apparent at the SCN and its direct targets, as a consequence of high viral replication (Beier et al., 2011). Surprisingly, we observed substantial anterograde viral spread to the dorsal hippocampus. Initial labeling was strongest in the CA2 region, with labeled axons projecting to CA3 and the Dentate Gyrus (DG) also observed in an anteroposterior gradient (**Figure 2**). This CA2-centered spread was visible from three days post-infection, with no labelling seen in the CA3 and DG at this timepoint, suggesting a direct SCN-to-hippocampus projection in the CA2 subfield .

**Figure 1.**
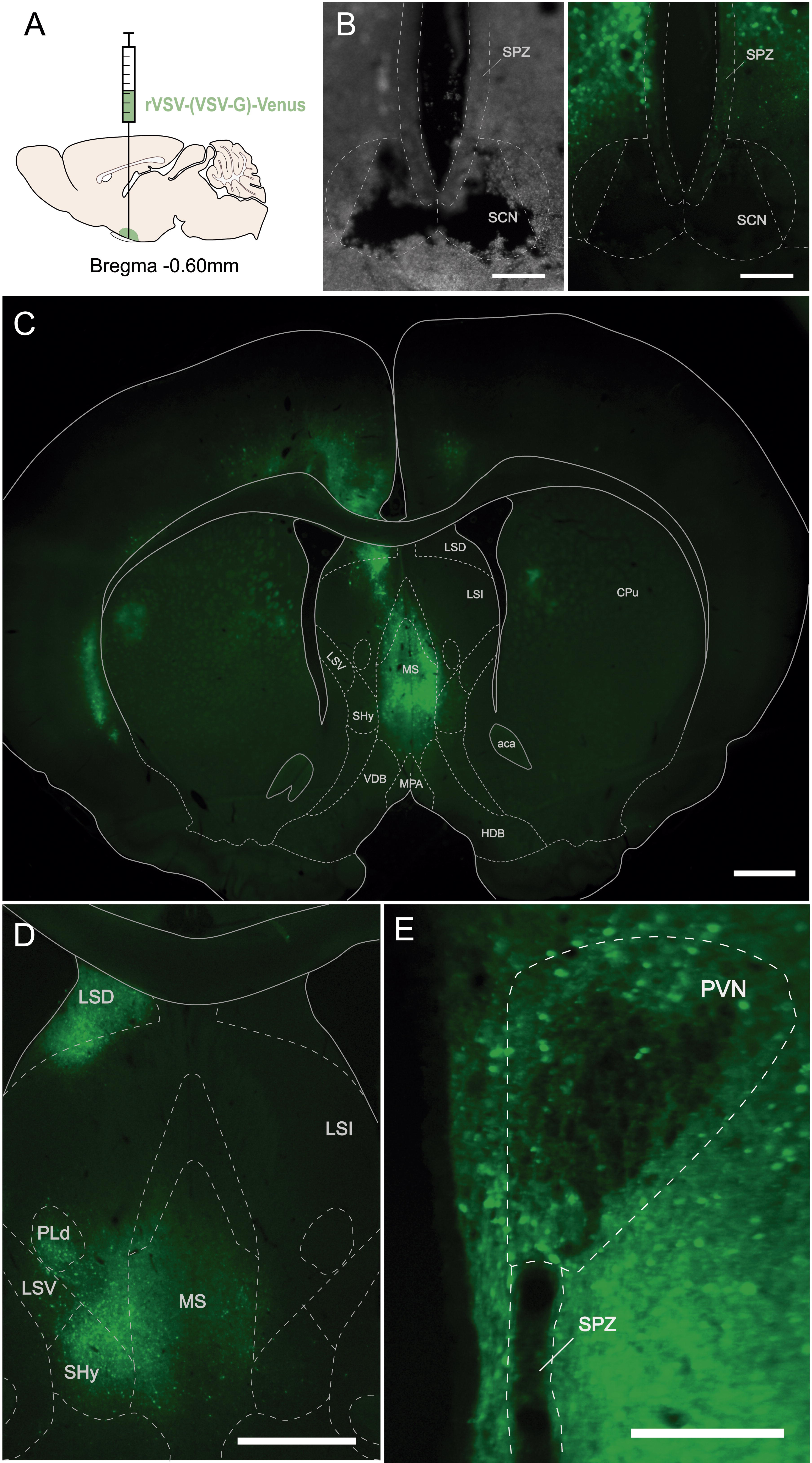
rVSV-(VSV-G)-Venus labeling of efferent targets after injection into the SCN. Regional localization of rVSV-(VSV-G)-Venus labelling six days after injection into the SCN. (A) Schematic representation of viral injection into the SCN. (B) Brightfield and fluorescence microscopic images of the injection spot, showing focal lesions from viral replication. Scale Bar: 250 μm (C) Coronal brain section showing rVSV-(VSV-G)-Venus expression (green) in the Medial Septum (MS) and Lateral Septum (LS), expected targets for anterograde spread from the SCN. Scale Bar: 1000 μm (D) Magnification showing Venus-labeled neurons in Dorsolateral Septal Nucleus (LSD), Ventrolateral Septal Nucleus (LSV) , MS and adjacent brain regions. Scale Bar: 1000 μm (E) Focal lesions on the Paraventricular Hypothalamic Nucleus (PVH) and Subparaventricular Zone (SPZ), indicating high viral replication in these regions. Scale Bar: 500 μm. Abbreviations: *Caudate–Putamen (CPu); Horizontal Diagonal Band of Broca (HDB); Lateral Septum, Dorsal (LSD); Lateral Septum, Intermediate (LSI); Lateral Septum, Ventral (LSV); Medial Septum (MS); Nucleus Accumbens (Acb/NAc); Paraventricular Nucleus of the Hypothalamus (PVN); Septohypothalamic Nucleus (SHy); Subparaventricular Zone*

**Figure 2.**
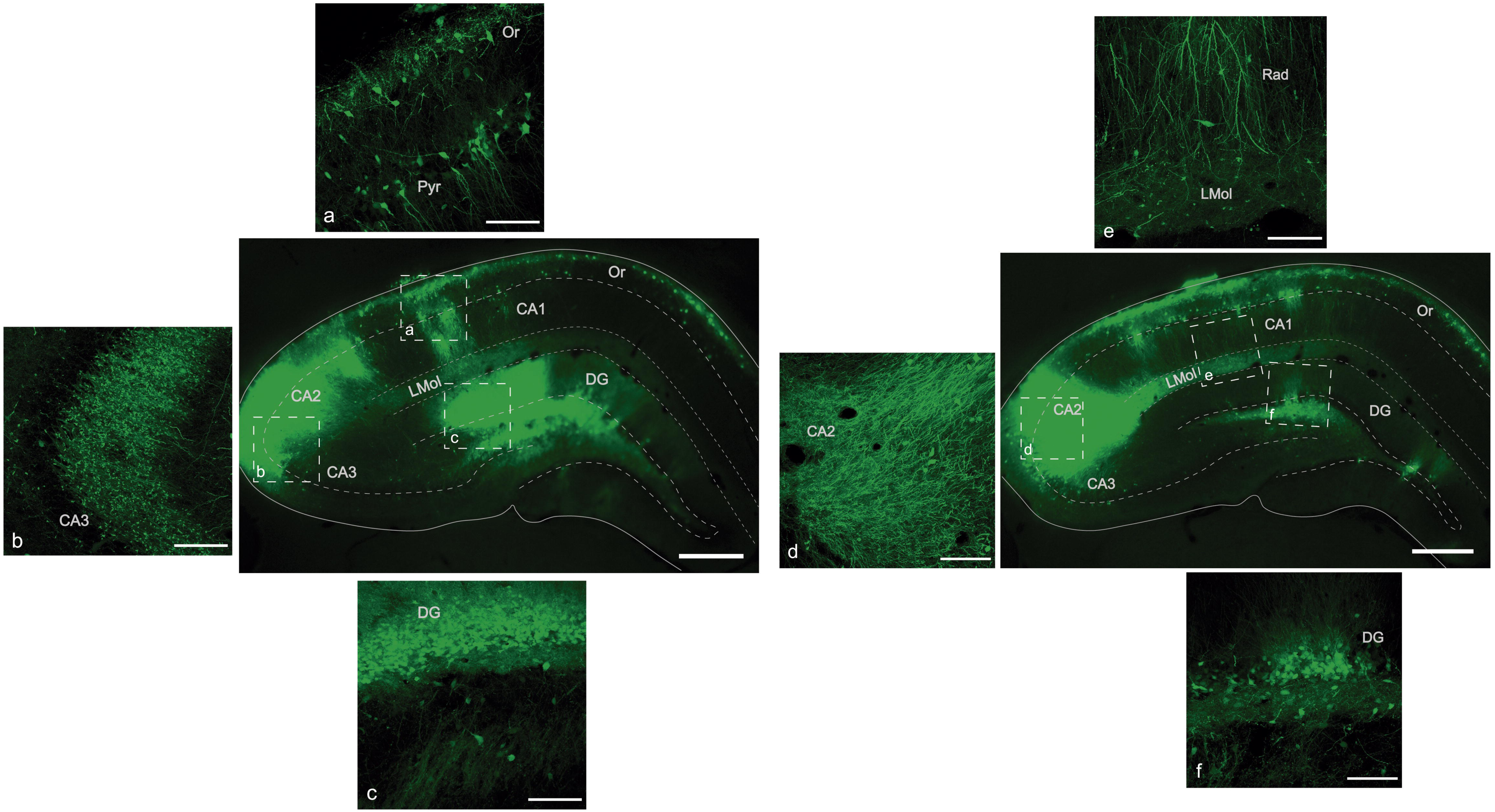
A comprehensive overview of rVSV-(VSV-G)-Venus labeling via transsynaptic anterograde spread from the SCN in distinct hippocampal subfields Intrahippocampal distribution of rVSV-(VSV-G)-Venus labeling six days after injection into the SCN. Scale bars: 500 μm. (A) Higher magnification revealing a dense population of Venus-labeled neurons in *strata oriens* and *pyramidale* of the dorsal hippocampal CA1 region. Scale bar: 100 μm. (B) Higher magnification revealing Venus-labeled fibers in the *stratum lucidum* of dorsal hippocampal CA3 region. Scale bar: 100 μm. (C) Higher magnification revealing a dense population of Venus-labeled neurons in the granular and molecular layers of the suprapyramidal blade of the Dentate Gyrus. Venus-labeled neurons and fibers in hilus of the Dentate Gyrus are also visible. Scale bar: 100 μm. (D) Higher magnification revealing Venus-labeled fibers in the dorsal hippocampal CA2 region. Scale bar: 100 μm. (E) Higher magnification revealing Venus-labeled axons in *stratum radiatum* and Venus-labeled cell bodies and fibers in *stratum lacunosum moleculare* of the dorsal hippocampal CA1 region. Scale bar: 100 μm. (F) Higher magnification revealing Venus-labeled neurons in the granular layer of the suprapyramidal blade of the Dentate Gyrus as well as Venus-labeled neurons and fibers in the subgranular zone. Scale bar: 100 μm. Abbreviations: *CA1 field of the hippocampus (CA1); CA2 field of the hippocampus (CA2); CA3 field of the hippocampus (CA3); Dentate Gyrus (DG); Lacunosum–Moleculare layer (LMol); Oriens layer (Or); Pyramidal cell layer (Pyr); Radiatum layer (Rad)*.

**Figure 3.**
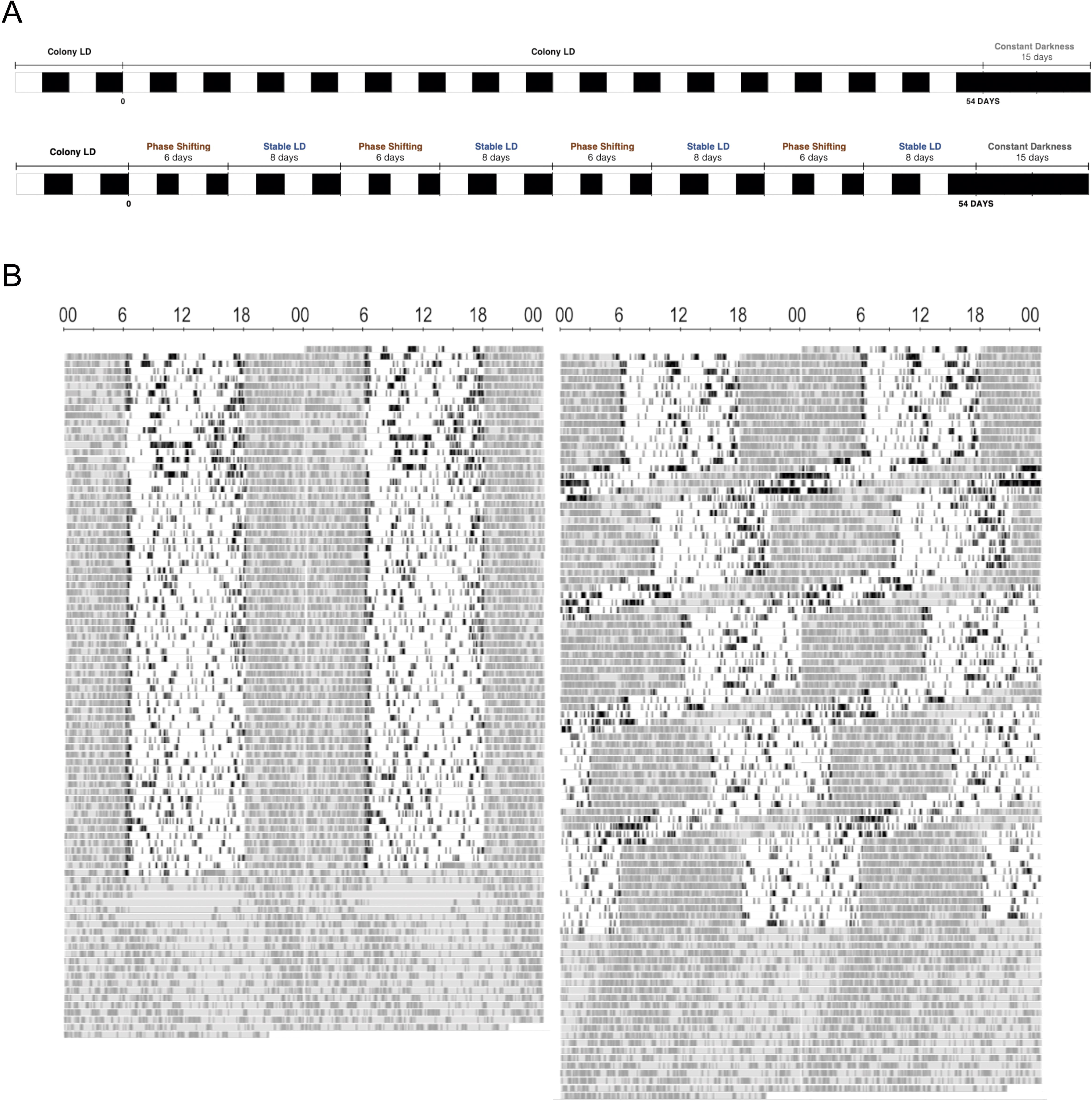
Chronic manipulation of the light period successfully alters the circadian rhythm of CCD/Shifted animals (A) Schematic representations of the light-dark cycle imposed on control animals (top) and on shifted animals (bottom). While control animals were kept under a normal 12:12h light/dark (LD) cycle, the shifted animals were subjected to 4 cycles of phase shifts followed by stable 12:12h LD cycle, to continually challenge the entrainment properties of the circadian system. After 54 days, control and shifted animals were kept under constant darkness to analyze free-running circadian rhythmicity and the after-effects of the chronic circadian manipulation. (B) Representative actograms from a control (left) and a shifted animal (right). Activity data is double-plotted for clarity. Each horizontal line corresponds to a 2-day interval, with the vertical bands denoting activity. The shaded regions indicate the periods when the lights were turned off.

**Figure 4.**
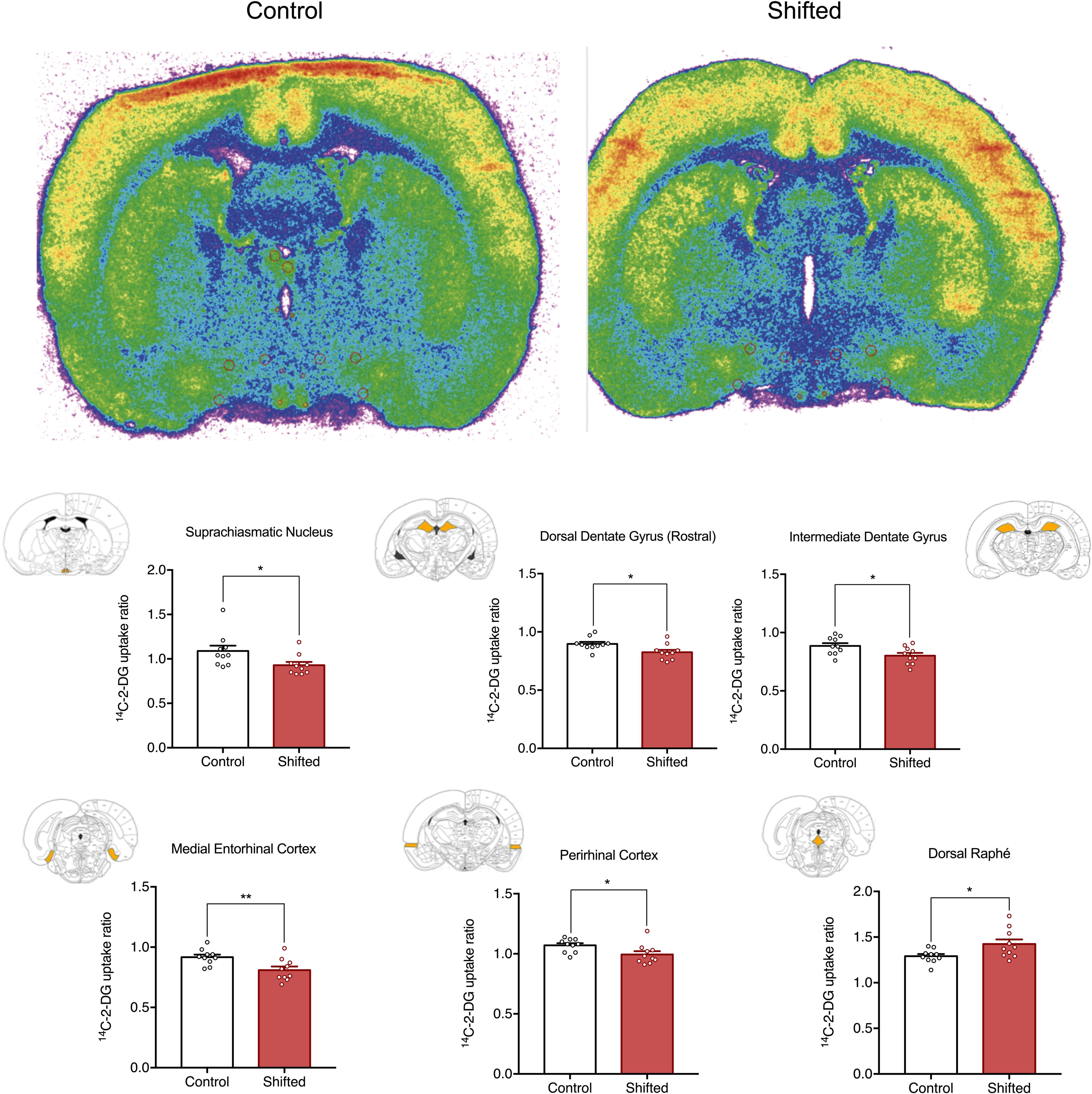
Chronic manipulation of circadian rhythms induces significant alterations in regional brain metabolism (A) Representative pseudo-color autoradiograms showing metabolic alterations induced by the chronic manipulation of circadian rhythms. Autoradiograms are of coronal sections at the level of the rostral Suprachiasmatic Nucleus (rSCN), where hypometabolism is seen. Higher rates of metabolism are indicated by warm colors (red/ yellow) and lower rates of metabolism are indicated by colder colors (green/blue). (B) CCD significantly decreased LCGU (^14^C-2-DG uptake ratio) in the Rostral Suprachiasmatic Nucleus (Mean difference: -0.162, 95% CI: -0.317 to -0.051), Rostrodorsal Dentate Gyrus (Mean difference: -0.073, 95% CI: -0.121 to -0.018), Intermediate Dentate Gyrus (Mean difference: -0.085, 95% CI: -0.146 to -0.020), Medial Entorhinal Cortex (Mean difference: -0.104, 95% CI: 0.164 to -0.035) and Perirhinal Cortex (Mean difference: -0.076, 95% CI: -0.124 to -0.004) while increasing LCGU in Dorsal Raphé (Mean difference: 0.134, 95% CI: 0.044 to 0.253). Schematic diagrams depict the anatomical location of each region in the rat brain. Data shown as the mean ± SEM (n=10, * p<0.05, ** p<0.01, unpaired Student’s t-test).

Comprehensive mapping at higher resolution confirmed widespread SCN-derived input across multiple hippocampal subfields (**Supplementary Figure 1**). Venus-labeled terminals were found in *stratum oriens*, *pyramidale, radiatum*, and *lacunosum moleculare* of CA1 and CA2. In the CA3 region, labeled fibers were detected mostly in the stratum lucidum. In the DG, we observed dense terminal labeling in the granular and molecular layers, as well as in the hilus and subgranular zone.

To corroborate the anterograde labelling results, we performed a complementary experiment: injecting the retrograde transsynaptic tracer rVSV-(RABV-G)-eGFP into the dorsal hippocampus (CA2 subfield). Five days post-injection, retrogradely labeled neurons were found in the SCN, confirming the direct SCN-hippocampus projection (**Supplementary Figure 2**). Additional labeling in MS, LS, and PVH was also identified, suggesting these regions may act as potential intermediaries in the SCN–Hippocampal pathway, given the established structural connectivity of these regions to the hippocampus (Watts et al., 1987a).

#### Neurochemical Identity of Labeled Hippocampal Neurons

To identify the hippocampal neurons targeted by SCN axons we next characterized the neurotransmitter phenotype of anterogradely Venus-labeled neurons using immunohistochemistry. In CA1 stratum oriens, Venus-positive neurons were surrounded by ChAT+ cholinergic terminals and showed appositions with GAD65/67+ GABAergic synapses (**Supplementary Figure 3**). Similar appositions of Venus-positive neurons with cholinergic and GABAergic terminals was also seen in CA2 and CA3 regions (**Supplementary Figures 4 and 5**), with some Venus-labeled neurons co- localizing with CamKII, a glutamatergic marker, allowing us to classify these targets of SCN axons as pyramidal (excitatory) glutamatergic neurons

**Figure 5.**
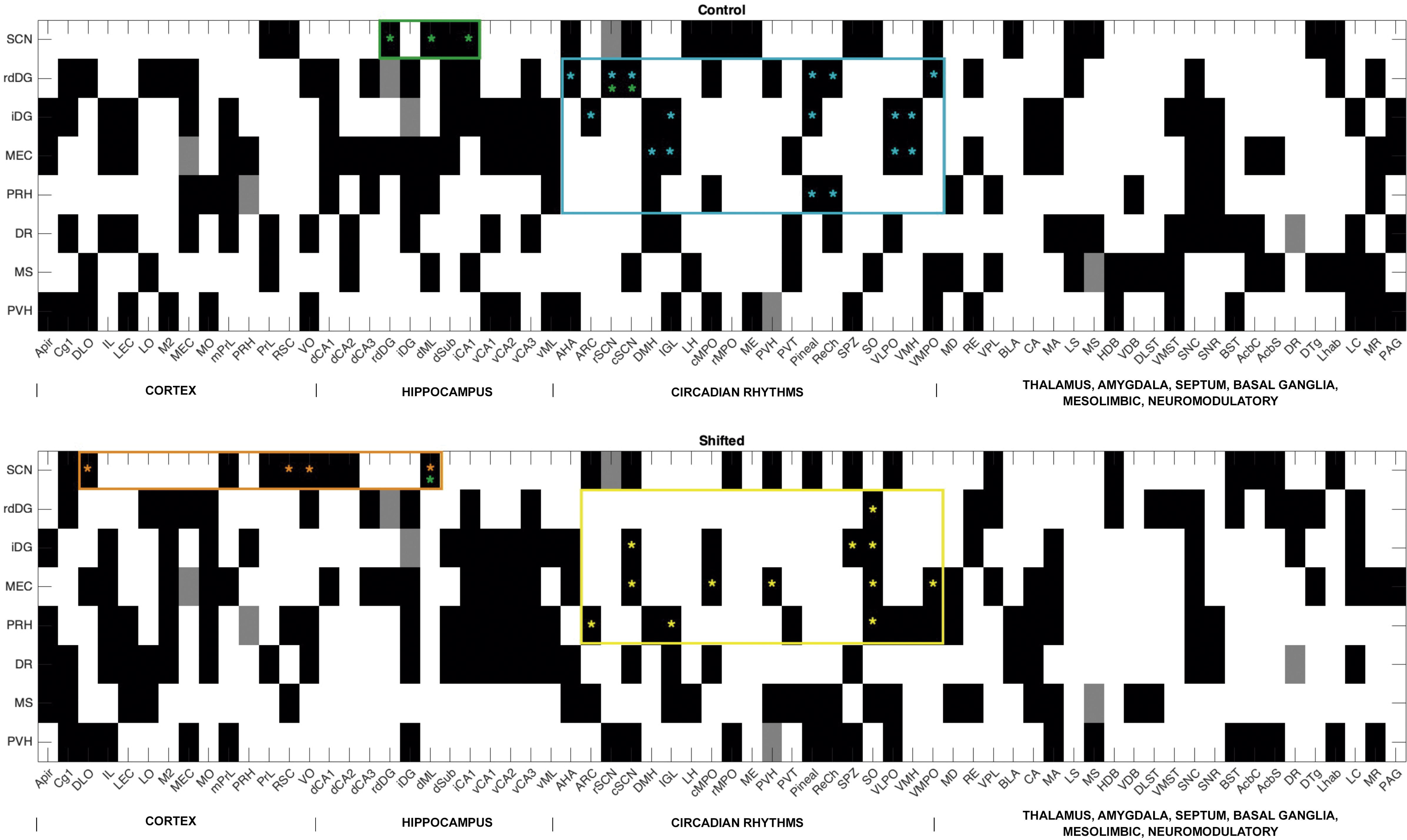
Visual representation of the connectivity pattern of control and shifted (CCD) groups 8 x 68 matrices displaying the connectivity patterns between each seed region and all brain regions of interest (RoIs) in control and shifted groups. Black denotes functionally connected regions, as per 95% Confidence Interval (95% CI) of the variable importance to the projection (VIP) statistic > 1.0 in the experimental group. The green bordering highlights connections between SCN, or its efferents, and memory-related regions that were identified by our anatomical tracing experiments; Orange bordering highlights connections between SCN and memory-related regions that are only present in the group that was subjected to chronic manipulation of circadian rhythms; Blue bordering highlights connections between memory-related regions and SCN efferents that are present in control animals. Yellow bordering highlights connections between memory-related regions and SCN efferents that are only present in the group that was subjected to chronic manipulation of circadian rhythms. The rat connectogram database ((Schmitt & Eipert, 2012)) was used as a tool to identify novel functional connections. **Cortex:** Amygdalo- Piriform Cortex (APir), Cingulate (Cg), Dorsolateral Orbital (DLO), Infralimbic (IL), Lateral Entorhinal (LEC), Lateral Orbital (LO), Supplementary Motor Area (M2), Medial Entorhinal (MEC), Medial Orbital (MO), Medial Prefrontal (mPFC), Prelimbic (PrL), Retrosplenial (RSC), Ventral Orbital (VO). **Hippocampus:** Dorsal CA1 (dCA1), Dorsal CA2 (dCA2), Dorsal CA3 (dCA3), Dorsal Dentate Gyrus (dDG), Intermediate Dentate Gyrus (iDG), Dorsal Hippocampus Molecular Layer (dML), Dorsal Subiculum (dSub), Intermediate CA1 (iCA1), Intermediate CA3 (iCA3), Ventral CA1 (vCA1), Ventral CA2 (vCA2), Ventral CA3 (vCA3), Ventral Dentate Gyrus (vDG), Ventral Hippocampus Molecular Layer (vML). **Circadian Rhythms**: Anterior Hypothalamus (AHA), Arcuate Nucleus (ARC), Suprachiasmatic Nucleus – Rostral (rSCN), Suprachiasmatic Nucleus – Caudal (cSCN), Dorsomedial Hypothalamus (DMH), Intergeniculate Leaflet (IGL), Lateral Hypothalamus (LH), Medial Preoptic Nucleus – Caudal (cMPO), Medial Preoptic Nucleus – Rostral (rMPO), Median Eminence (ME), Paraventricular Hypothalamic Nucleus – Parvicellular (PVp), Paraventricular Thalamic Nucleus (PVT), Preoptic Area (POA), Retrochiasmatic Area (RCH), Supraoptic Nucleus (SON), Suprachiasmatic Nucleus (SCN), Ventromedial Hypothalamus (VMH), Ventral Thalamic Nucleus (VL). **Thalamus**: Mediodorsal (MD), Reuniens (RE). **Amygdala**: Basolateral (BLA), Central (CeA), Medial (MeA). **Septum**: Lateral Septum (LS), Medial Septum (MS). Diagonal Band of Broca: Horizontal Band (HDB), Vertical Band (VDB). **Basal Ganglia**: Dorsolateral Striatum (DLS), Ventromedial Striatum (VMS), Substantia Nigra pars compacta (SNc), Substantia Nigra pars reticulata (SNr). **Mesolimbic**: Bed Nucleus of the Stria Terminalis (BST), Nucleus Accumbens core (AcbC), Nucleus Accumbens shell (AcbSh). **Neuromodulatory**: Dorsal Raphe (DR), Dorsal Tegmental Nucleus (DTg), Lateral Habenula (LHb), Locus Coeruleus (LC), Median Raphe (MR), Periaqueductal Gray (PAG).

In the DG, Venus-labeled neurons were immunopositive for GAD65/67, ChAT appositions, and CamKII (**Supplementary Figures 6 and 7**), confirming that, contrary to the other regions, both inhibitory and excitatory neurons are targeted by SCN axons in the DG. Notably, the majority of Venus-labeled neurons across DG subfields were GABAergic, while a minority in the hilus co- localized with CamKII, suggesting targeting of a subset of glutamatergic mossy cells.

**Figure 6.**
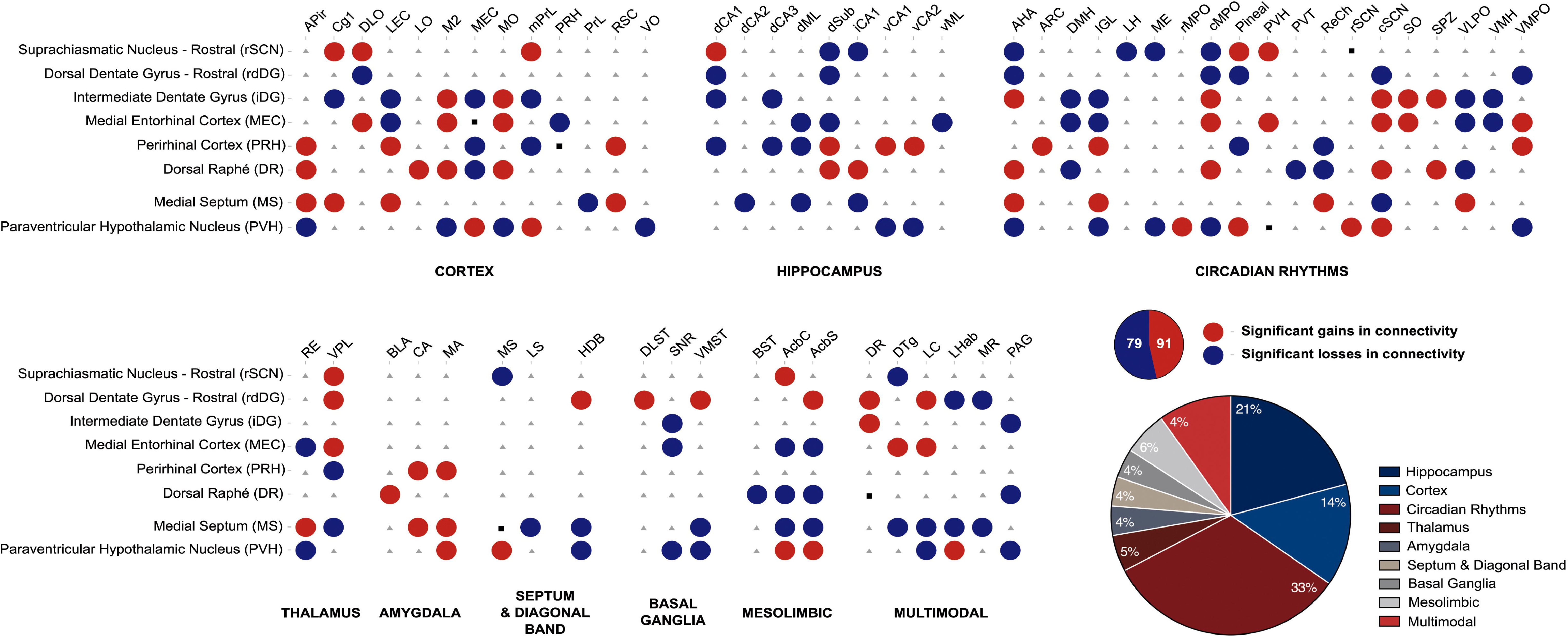
Significant gains and losses in brain region functional connectivity (FC) induced by chronic circadian disruption (CCD). (A) Summary of significant changes in regional FC induced by chronic circadian manipulation. Red denotes significantly gained connectivity, whereas blue represents significantly lost connectivity, in CCD animals in comparison to controls. Inter-regional connectivity changes were determined by statistical comparison of the VIP statistic obtained from PSLR models of the LCGU data, compared across groups using unpaired Student’s t-test with Bonferroni *post hoc* correction for multiple comparisons. Significance was set at p < 0.05. Full data for each seed region are shown in the Supplementary Tables 1 – 10. (B) Pie charts showing the proportion of gained or lost connectivity, and the percentage of FC changed, by CCD within each functional brain region group. **Cortex:** Amygdalo-Piriform Cortex (APir), Cingulate (Cg), Dorsolateral Orbital (DLO), Infralimbic (IL), Lateral Entorhinal (LEC), Lateral Orbital (LO), Supplementary Motor Area (M2), Medial Entorhinal (MEC), Medial Orbital (MO), Medial Prefrontal (mPFC), Prelimbic (PrL), Retrosplenial (RSC), Ventral Orbital (VO). **Hippocampus:** Dorsal CA1 (dCA1), Dorsal CA2 (dCA2), Dorsal CA3 (dCA3), Dorsal Dentate Gyrus (dDG), Intermediate Dentate Gyrus (iDG), Dorsal Hippocampus Molecular Layer (dML), Dorsal Subiculum (dSub), Intermediate CA1 (iCA1), Intermediate CA3 (iCA3), Ventral CA1 (vCA1), Ventral CA2 (vCA2), Ventral CA3 (vCA3), Ventral Dentate Gyrus (vDG), Ventral Hippocampus Molecular Layer (vML). **Circadian Rhythms**: Anterior Hypothalamus (AHA), Arcuate Nucleus (ARC), Suprachiasmatic Nucleus – Rostral (rSCN), Suprachiasmatic Nucleus – Caudal (cSCN), Dorsomedial Hypothalamus (DMH), Intergeniculate Leaflet (IGL), Lateral Hypothalamus (LH), Medial Preoptic Nucleus – Caudal (cMPO), Medial Preoptic Nucleus – Rostral (rMPO), Median Eminence (ME), Paraventricular Hypothalamic Nucleus – Parvicellular (PVp), Paraventricular Thalamic Nucleus (PVT), Preoptic Area (POA), Retrochiasmatic Area (RCH), Supraoptic Nucleus (SON), Suprachiasmatic Nucleus (SCN), Ventromedial Hypothalamus (VMH), Ventral Thalamic Nucleus (VL). **Thalamus**: Mediodorsal (MD), Reuniens (RE). **Amygdala**: Basolateral (BLA), Central (CeA), Medial (MeA). **Septum**: Lateral Septum (LS), Medial Septum (MS). Diagonal Band of Broca: Horizontal Band (HDB), Vertical Band (VDB). **Basal Ganglia**: Dorsolateral Striatum (DLS), Ventromedial Striatum (VMS), Substantia Nigra pars compacta (SNc), Substantia Nigra pars reticulata (SNr). **Mesolimbic**: Bed Nucleus of the Stria Terminalis (BST), Nucleus Accumbens core (AcbC), Nucleus Accumbens shell (AcbSh). **Neuromodulatory**: Dorsal Raphe (DR), Dorsal Tegmental Nucleus (DTg), Lateral Habenula (LHb), Locus Coeruleus (LC), Median Raphe (MR), Periaqueductal Gray (PAG).

**Figure 7.**
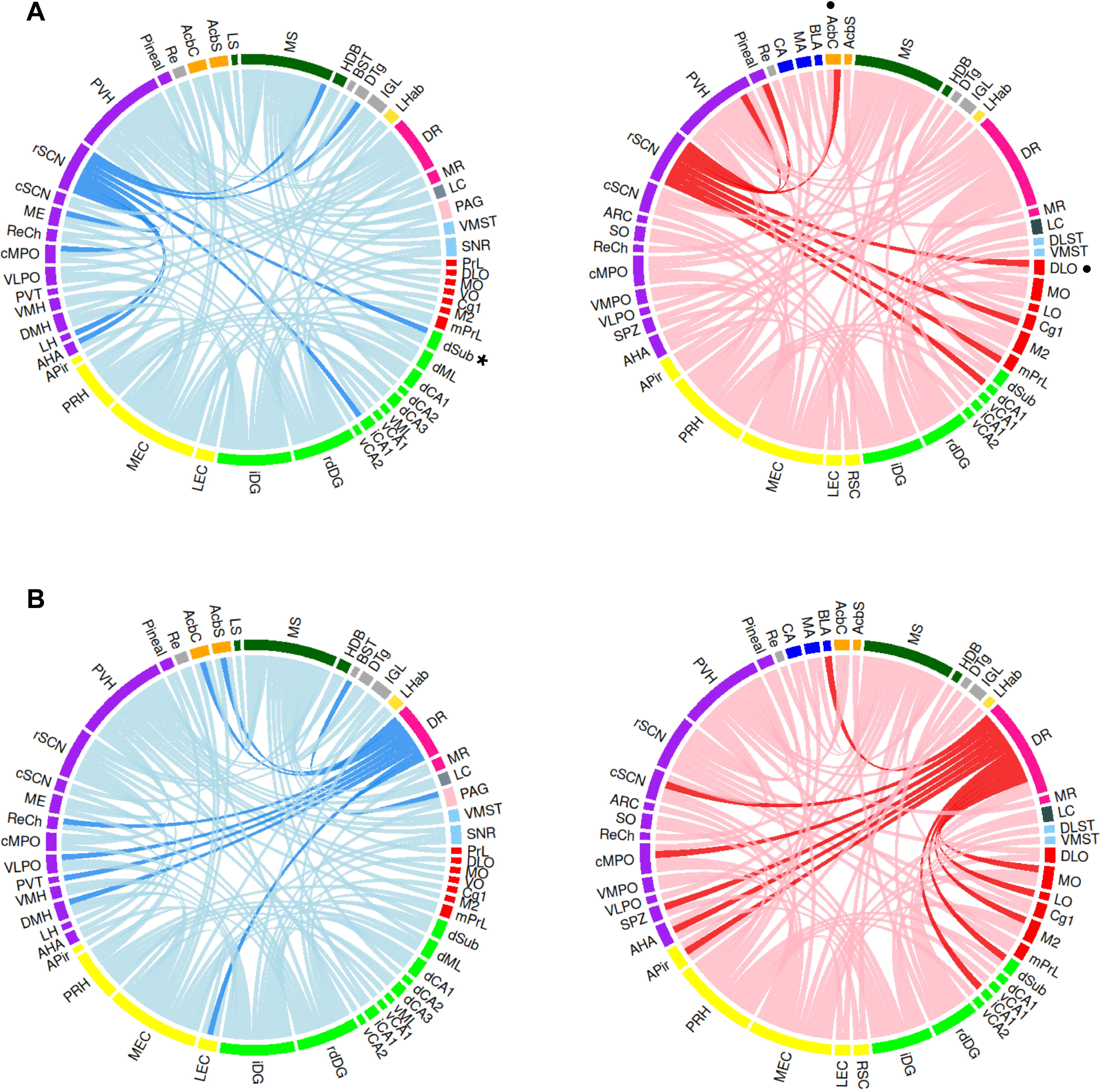
Chronic circadian disruption (CCD) alters rostral suprachiasmatic nucleus (SCN) and dorsal raphe (DR) functional connectivity (FC). Chord diagrams summarizing significant alterations in the FC of the A) rostral SCN and B) Dorsal raphe (DR). Significant losses in FC in CCD animals are highlighted in dark blue. *denotes unreported functional connections with memory-related regions that were physiologically present and were lost upon chronic circadian manipulation. Significant gains in FC in CCD animals are highlighted in dark red. **•** denotes unreported functional connections with memory-related regions that were gained upon chronic circadian manipulation. The significance of FC alterations was determined from the VIP statistic using unpaired Student’s t-test with Bonferroni *post hoc* correction. Significance was set a p<0.05. Inter-regional FC exists in an experimental group if the 95% CI of the VIP statistic > 1.0. Faded colors represent total significant lost/gained connections across all seed regions (see supplemental material for other brain regions). Full regional FC data are shown in the Supplementary material (Tables 1-10). **Cortex:** Amygdalo-Piriform Cortex (APir), Cingulate (Cg), Dorsolateral Orbital (DLO), Infralimbic (IL), Lateral Entorhinal (LEC), Lateral Orbital (LO), Supplementary Motor Area (M2), Medial Entorhinal (MEC), Medial Orbital (MO), Medial Prefrontal (mPFC), Prelimbic (PrL), Retrosplenial (RSC), Ventral Orbital (VO). **Hippocampus:** Dorsal CA1 (dCA1), Dorsal CA2 (dCA2), Dorsal CA3 (dCA3), Dorsal Dentate Gyrus (dDG), Intermediate Dentate Gyrus (iDG), Dorsal Hippocampus Molecular Layer (dML), Dorsal Subiculum (dSub), Intermediate CA1 (iCA1), Intermediate CA3 (iCA3), Ventral CA1 (vCA1), Ventral CA2 (vCA2), Ventral CA3 (vCA3), Ventral Dentate Gyrus (vDG), Ventral Hippocampus Molecular Layer (vML). **Circadian Rhythms**: Anterior Hypothalamus (AHA), Arcuate Nucleus (ARC), Suprachiasmatic Nucleus – Rostral (rSCN), Suprachiasmatic Nucleus – Caudal (cSCN), Dorsomedial Hypothalamus (DMH), Intergeniculate Leaflet (IGL), Lateral Hypothalamus (LH), Medial Preoptic Nucleus – Caudal (cMPO), Medial Preoptic Nucleus – Rostral (rMPO), Median Eminence (ME), Paraventricular Hypothalamic Nucleus – Parvicellular (PVp), Paraventricular Thalamic Nucleus (PVT), Preoptic Area (POA), Retrochiasmatic Area (RCH), Supraoptic Nucleus (SON), Suprachiasmatic Nucleus (SCN), Ventromedial Hypothalamus (VMH), Ventral Thalamic Nucleus (VL). **Thalamus**: Mediodorsal (MD), Reuniens (RE). **Amygdala**: Basolateral (BLA), Central (CeA), Medial (MeA). **Septum**: Lateral Septum (LS), Medial Septum (MS). Diagonal Band of Broca: Horizontal Band (HDB), Vertical Band (VDB). **Basal Ganglia**: Dorsolateral Striatum (DLS), Ventromedial Striatum (VMS), Substantia Nigra pars compacta (SNc), Substantia Nigra pars reticulata (SNr). **Mesolimbic**: Bed Nucleus of the Stria Terminalis (BST), Nucleus Accumbens core (AcbC), Nucleus Accumbens shell (AcbSh). **Neuromodulatory**: Dorsal Raphe (DR), Dorsal Tegmental Nucleus (DTg), Lateral Habenula (LHb), Locus Coeruleus (LC), Median Raphe (MR), Periaqueductal Gray (PAG).

**Table 1.**
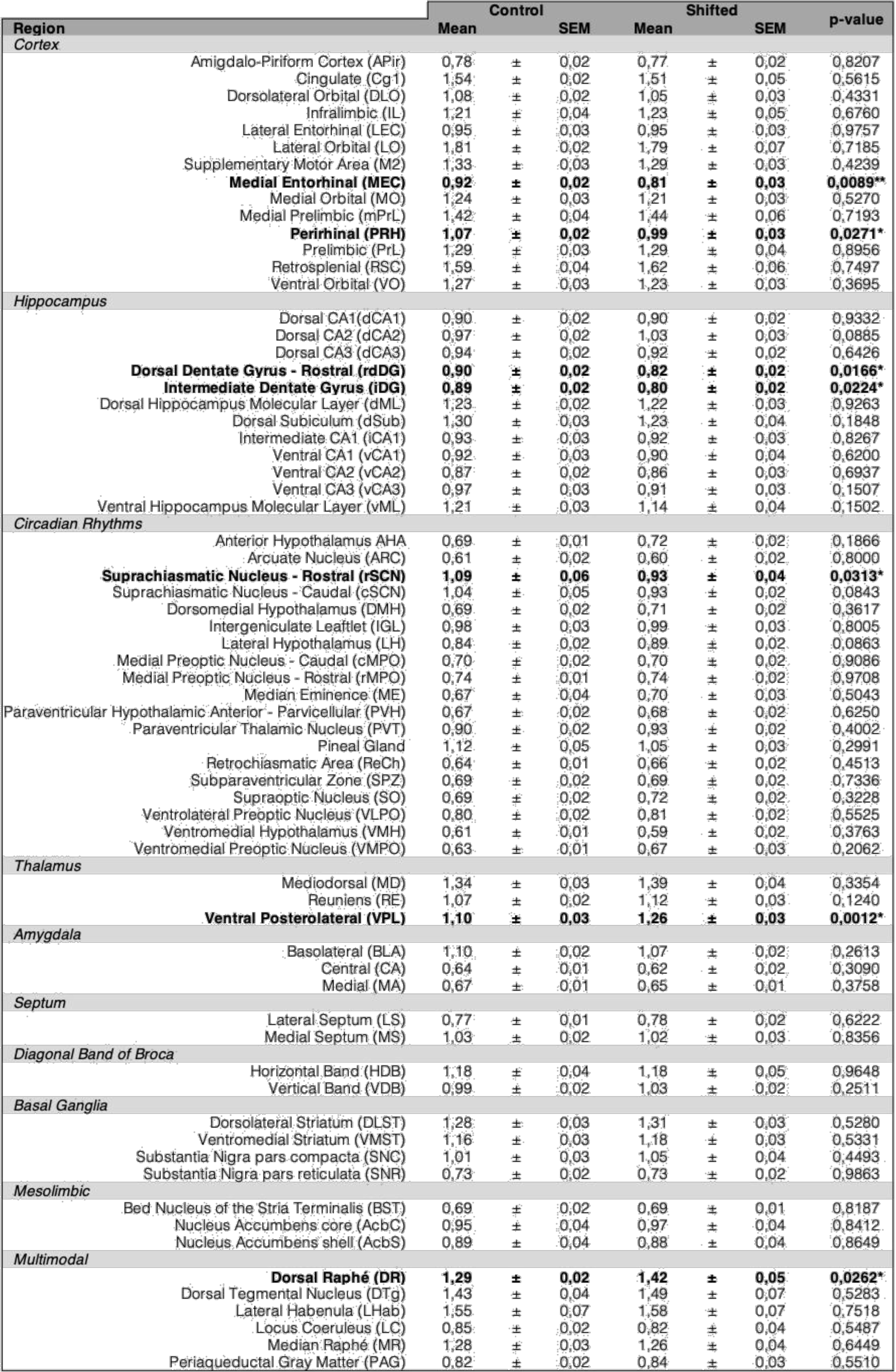
Impact of chronic circadian disturbance (CCD) on regional local cerebral glucose utilization (LCGU) LCGU was evaluated across 68 regions of interest (RoIs) across a diverse range of neural systems. LCGU was calculated as the ^14^C-2-Deoxyglucose uptake ratio, the ratio of ^14^C-2-DG present in each RoI relative to the average ^14^C-2-DG concentration in the whole brain of the same animal. Bold denotes significant differences between groups. * p<0.05, ** p<0.01, unpaired Student’s t-test with Bonferroni correction.

#### Monosynaptic Tracing Confirms Direct SCN–Hippocampal Projecting Neurons

To identify direct SCN to hippocampus projections free from the possible viral expression artifacts, we injected an inert monosynaptic tracer (BDA-Texas Red) into the SCN. Terminal labeling was observed in expected first-order targets including the BNST, PVH, LS, and MS, validating our viral tracing observations (**Supplementary Figure 8**). Notably, terminal labeling was also present in the hippocampus, including the hilus of the DG, CA2 and CA3 (**Supplementary Figure 9**). High- magnification imaging revealed neurons with clustered terminals in these regions (**Supplementary Figure 11,a-h**), indicative of direct synaptic contact, thus confirming the viral tracing results in the hippocampus. No BDA signal was observed in control tissue processed without primary antibody, confirming the specificity of the monosynaptic labeling (**Supplementary Figure 11i**).

Collectively, these results provide converging transsynaptic and monosynaptic evidence of a direct synaptic connection between the SCN and the dorsal hippocampus. This projection strongly and directly targets CA2 making it a potential entry point into the hippocampus. As such, these projections are strategically located to provide circadian modulation of hippocampal function, and thus may have a profound influence over the temporal regulation of memory consolidation and synaptic plasticity. To understand such implications, we then investigated the functional consequences of chronic circadian disturbance (CCD), on this previously underappreciated connection and associated memory- related circuits.

### Functional Consequences of CCD on the Brain and Brain Network Connectivity

#### CCD Induces Persistent Behavioral and Molecular Alterations

To investigate how circadian disruption affects brain function, we subjected rats to a protocol of **chronic photoperiod manipulation (eg shifted animals)**, involving repeated 3-hour phase-advances for six consecutive days followed by 10 days of stable entrainment, repeated four times. Control rats remained on a 12:12 h light/dark (LD) schedule throughout (**Figure 3**).

Activity recordings confirmed that rats under chronic phase-shifts developed fragmented rhythms with increased light-phase activity during shifts. Despite re-entrainment, these animals exhibited altered free-running periods in constant darkness (**Figure 3B**), consistent with a persistent effect of circadian manipulation on their internal clock.

To investigate mechanistic bases of such persistent disruption, we sought to identify biochemical correlates of circadian disturbance by studying the transcriptional profiles of target cells. We thus quantified hepatic mRNA levels of *Cd36* and *Cyp2c7*, established biomarkers of circadian disruption. Shifted animals exhibited increased *Cd36* and decreased *Cyp2c7* expression, consistent with known transcriptomic signatures of circadian stress (**Supplementary Figure 10**; Adamovitch et al. 2014).

Importantly, despite this physiological disruption, we found no differences in anxiety-like behaviour, as detected in the Elevated Plus Maze or the Open Field Test. Likewise, diurnal corticosterone profiles remained rhythmic and comparable across the experimental groups, indicating that altered mood and hypothalamic-pituitary-adrenal (HPA) axis activation were altered by CCD and did not confound our other observations (**Supplementary Figure 11**).

#### Chronic Circadian Manipulation Alters Regional Brain Metabolism

Using ^14^C-2-deoxyglucose autoradiography, we examined how circadian disturbance impacts **brain- wide metabolic activity,** as a reflection of regional activity. While whole-brain average glucose uptake in experimental animals was not different from controls (**Supplementary Figure 12A**), we detected significant, region-specific changes in local cerebral glucose utilization (LCGU) in 7 out of 68 analyzed brain regions of interest (ROIs) (**Figure 4**). Full regional metabolic data are shown in **Table 1 and Suppl Figure 12B-H**. In this way, CCD animals exhibited **hypometabolism** in the rostral SCN, DG (rostrodorsal DG: rdDG and intermediate DG: iDG), Medial Entorhinal Cortex (MEC) and Perirhinal Cortex (PRH) (**Figure 4**). Conversely, **hypermetabolism** was observed in the serotonergic **Dorsal Raphé** (DR) and **Ventral Posterolateral Thalamus**. Pseudocolor representative autoradiograms showed reduced SCN activity in CCD animals, particularly in the shell region, highlighting the SCNs metabolic vulnerability to phase shifts (**Figure 4**).

#### Circadian Manipulation Disrupts Functional Brain Connectivity

We used **Partial Least Squares Regression (PLSR)** modeling to elucidate how chronic circadian disturbance affects interregional functional coupling. Given the significant changes in metabolic activity within the rSCN, rdDG, iDG, MEC, PRH and DR of the CCD/shifted animals, these regions were selected as "seed regions" for the functional regional connectivity analysis. Moreover, the Medial Septum (MS) and the Paraventricular Hypothalamic Nucleus (PVH) were also added as seed regions for this analysis, based on our structural connectivity results from tracing injections that suggest these brain regions as putative relays of circadian information from the SCN into the hippocampus (**Supplementary Figure 12**).

The rat connectome database (Schmitt & Eipert, 2012) was used as a reference to analyze the connectivity patterns resolved by our PLSR models. The accuracy of our PLSR models can be demonstrated as the control group matrix includes connections within cortico-hippocampal regions or the circadian system that are well-established and supported by numerous studies in the literature **(Figure 5 top)**.

Furthermore, our PLSR models uncovered functional connections between the SCN or its outputs and memory-related regions that are physiologically present in control animals but had not been reported in previous studies **(Figure 5, top, highlighted by green and blue *)**. Noteworthy, some of these connections between the SCN and memory-related regions, corroborate our structural connectivity results from tracing injections in the SCN **(Figure 5, highlighted by green *)**.

PLSR-based connectivity analysis revealed a profound functional reorganization of neural circuits (**Figure 5**). In CCD (shifted) animals, we found altered FC between SCN and memory-related structures whose anatomical interconnectivity we have described above using anatomical tracing. Notably, we found that some of these connections were present in controls, and were significantly lost in CCD animals, while other functional connections were not present in controls and were only found in CCD animals, suggesting an adaptive reorganization of SCN-HIPP functional circuitry Importantly, our PLSR model also revealed functional connectivity patterns between the SCN or its downstream projections and memory-associated brain regions that were exclusively present in the animals exposed to chronic circadian manipulation and had not been documented in prior research (Figure 5, bottom, highlighted by orange and yellow *).

Visualization of unfiltered variable importance for projections (VIP) statistics allows us to observe the contribution of all quantified connections for the full data variance, thus highlighting the ones that change most between experimental conditions (Supplementary **Figure 13**. This plot shows reduced coupling within dorsal hippocampal subfields, MS, and related cortices, increased coupling of SCN with cortico-hippocampal regions, and enhanced FC of ventral hippocampus, potentially indicating circuit reorganization.

Statistical comparison of inter-regional variable importance to the project (VIP) scores revealed **170 significantly altered functional connections** in CCD animals, with 39% of these evident in memory- related regions (hippocampus, cortex, amygdala), and 33% evident in the extended circadian circuitry (**Figure 6**). These changes underscore widespread functional decoupling and reorganization due to circadian disturbance.

Chord diagrams for each PLSR seed region highlight specific patterns of gain and loss of functional connectivity for these regions (**Figure 7** and **Supplemental Figures 14-15**). We summarise these inter-regional connectivity changes below.

- **SCN**: Lost connectivity with core outputs (Medial Septum, Anterior Hypothalamus, Lateral Hypothalamus, Median Eminence, and Medial Preoptic Nucleus), but gained abnormal connectivity to dorsal hippocampus CA1, prefrontal cortex, and pineal gland (**Figure 7A**).
- **DR**: Note increased DR connectivity to the SCN, hippocampus and PFC in CCD animals, supporting altered serotonergic modulation of these systems (Glass et al., 2000; **Figure 7B**)
- **Rostrodorsal DG (rdDG)**: Lost connectivity with SCN, as well as several of its afferents, and other hippocampal regions, including dorsal CA1, but gained coupling to Nucleus Accumbens, DR, and Locus Coeruleus—regions providing neuromodulatory input into the hippocampus (Lisman et al., 2011) (**Supplementary Figure 14A**).
- **Intermediate Dg (iDG)**: Lost canonical links with entorhinal cortex and prefrontal cortex, but gained FC with SCN, SPZ, and Supraoptic Nucleus, suggesting remapping of theta-rhythmic inputs (Buzsáki, 2002) (**Supplementary Figure 14B**).
- **MEC and PRH**: Showed loss of hippocampal and cortical connectivity, replaced by gain of novel cortico-cortical and SCN connections (**Supplementary Figure 14C and 15A**).
- **PVH**: Became disconnected from several hypothalamic and cortical regions but gained novel functional connectivity from the hippocampus, entorhinal cortex, and SCN, suggesting potential compensation or altered function in hormonal regulation pathways (**Supplementary Figure 15B**).
- **MS**: This neuromodulatory hub lost substantial connectivity with both circadian and memory networks **(Supplementary Figure 15C).**

### Chronic Manipulation of Circadian Rhythms Selectively Impairs Object Recognition Memory in the Absence of Hippocampal Synaptic Plasticity Changes

To assess the impact of CCD on behaviour and cognitive functions, a comprehensive battery of behavioral tests was performed following the final phase-shifting and re-entrainment cycle.

We found evidence for a deficit in learning and memory as detected using the **Novel Object Recognition (NOR) test**, which assesses recognition memory based on rodents’ innate preference for novel over familiar objects. During the encoding phase of NOR, both experimental groups explored all objects equally, indicating similar levels of attention and motivation. In the retrieval phase, control animals showed a strong preference for the novel object, while shifted animals did not discriminate between familiar and novel stimuli (**Figure 8**). This resulted in a **significant reduction in the Discrimination Index** in the shifted group (**Figure 8D**), indicating an impairment in object recognition memory. Given that NOR performance is particularly sensitive to perirhinal cortex, entorhinal cortex, ventral hippocampus and PFC-hippocampus circuit integrity (Wang et al. 2021) , this result may reflect altered function in these circuits— which is consistent with the hypometabolism and disrupted connectivity in this circuitry in CCD animals (see **Figure 4 and Supplementary Figure 14**).

**Figure 8.**
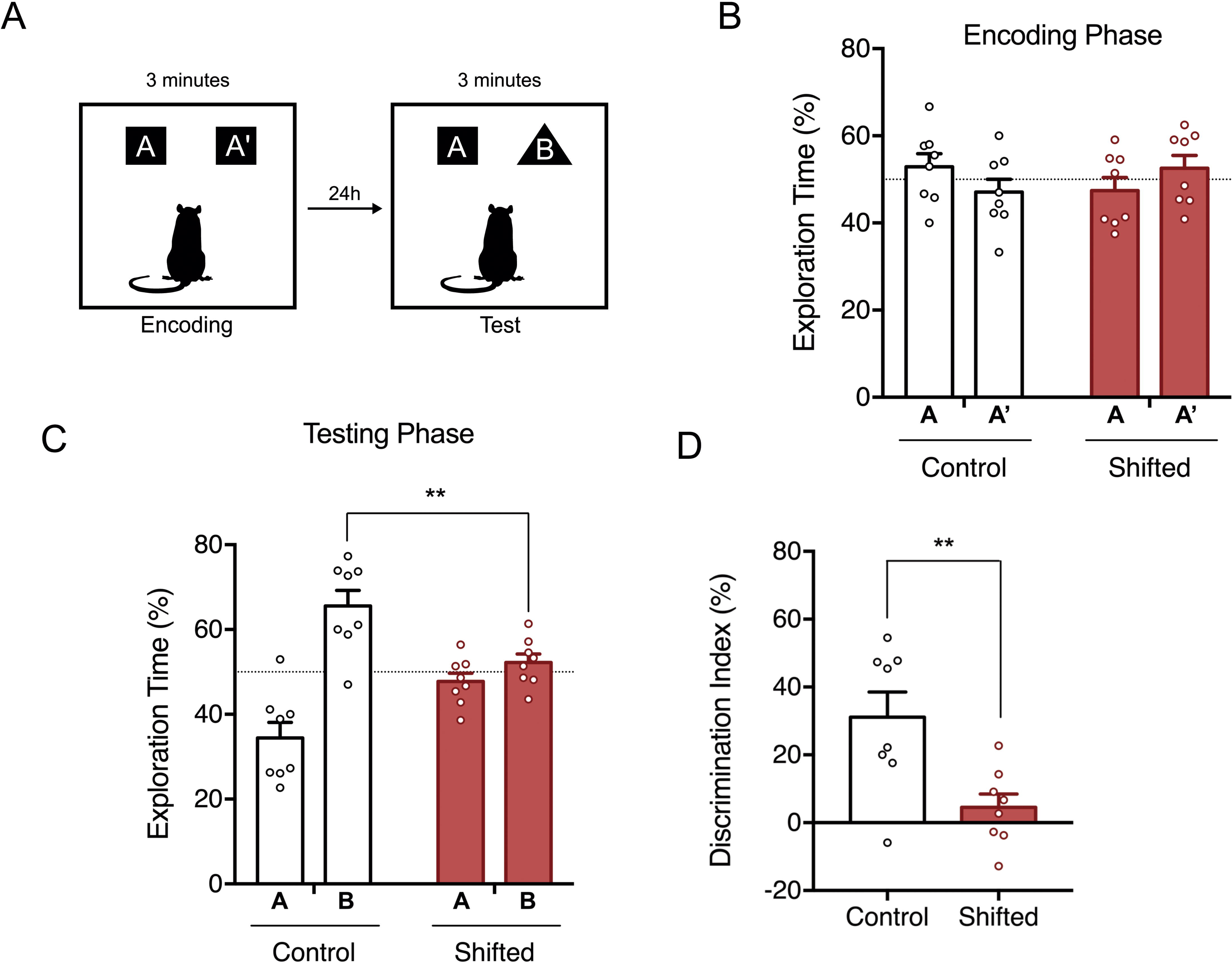
Chronic circadian disruption selectively impairs object recognition memory (A) Schematic representation of the Novel Object Recognition test (NORT), showing the encoding and testing phases with a 24 hour inter-trial interval. (B) Performance during the encoding phase was not significantly different between both groups. (C) CCD/shifted animals displayed a loss of preference for the novel object in the test phase of the NORT, and a significant difference in performance when compared to control animals. (n=8, ** p<0.01, unpaired Student’s t-test). (D) The Discrimination Index (DI) was significantly different between CCD and control animals (n=8, ** p<0.01, unpaired Student’s t-test), with CCD animals showing no novel object preference. All values are mean ± SEM.

Contrasting with the generally held assumption that circadian misalignment may globally impair cognition, we found that CCD animals had cognitive function spared in most domains. We found no significant differences between control and CCD animals in terms of spontaneous alternation, short- term memory, working memory, long-term memory, reversal learning or pattern descrimination, as assessed through T-maze, Y-maze, Morris Water Maze and Pattern Separation paradigms, respectively (**Supplementary Figure 16**).

To determine whether these behavioral impairments stemmed from altered synaptic plasticity in the hippocampus, we assessed **Long-Term Potentiation (LTP)** and **Long-Term Depression (LTD)** at the Schaffer collateral–CA1 synapse. There were no significant differences in LTP or LTD magnitude between control and CCD animals (**Supplementary Figure 17**). These findings suggest that the core mechanisms of synaptic plasticity within the dorsal hippocampus, at least at the Schaffer collateral- CA1 synapses, remained intact under circadian disturbance protocol employed, and that deficits in NORT performance were not related to dysfunction in these cellular plasticity processes in this projection.

Taken together, our results demonstrate that CCD induces a selective impairment in recognition memory, while sparing spatial and working memory. These cognitive deficits emerge independently of changes in hippocampal synaptic plasticity, thus they may be driven by the altered FC we have identified in SCN-memory circuits or neuromodulatory tone, rather than synaptic plasticity dysfunction. These results highlight the need for further research to elucidate the domain-specific vulnerability in circadian-related cognitive dysfunction and the systems-levels mechanisms involved.

## DISCUSSION

This study demonstrates that CCD leads to a reorganization of neural circuits linking the SCN with hippocampal and cortical regions, as reflected by regional metabolism and FC changes, and results in the selective impairment of recognition memory. The specificity of the behavioral deficits—combined with preserved hippocampal synaptic plasticity—suggests that disrupted large-scale brain network dysfunction, rather than synaptic failure, is a key driver of cognitive dysfunction in CCD.

### Direct and Indirect Pathways Linking the SCN to the Hippocampus

Early anatomical studies established the SCN as a hub with largely hypothalamus-confined outputs (Stephan, 1981; Berk & Finkelstein, 1982). Ogata et al. (1982), however, reported lesion-induced degeneration of SCN efferents in extrahypothalamic regions. Although provocative, these findings were not replicated by later comprehensive anterograde and retrograde tracing (Watts & Swanson, 1987a,b; Kriegsfeld et al., 2004), which further reinforced the consensus that SCN projections were largely restricted to hypothalamic and thalamic nuclei, with polysynaptic influence on limbic structures via intermediaries such as the lateral septum (Ruby, 2021). In this model of SNC anatomical connectivity, SCN outputs reach the hippocampus indirectly, through septal circuits that regulate hippocampal theta rhythms. However, methodological ambiguities in older studies may have overlooked sparse direct inputs. For instance, some retrograde tracer injections included both lateral and medial septal regions, complicating interpretation (Kriegsfeld et al, 2004). Sparse SCN-derived fibers have also been visualized in medial septum (Watts & Swanson, 1987a). More recently, pseudorabies virus tracing has revealed multi-neuronal links between SCN targets and limbic regions (Sylvester et al., 2002), and François et al. (2023) reported a sparse monosynaptic projection from SCN to the central amygdala. These data suggest that although SCN efferents are predominantly local, sparse long-range projections exist that can now be uncovered by using new, more sensitive, methods.

Using transsynaptic viral tracing, our study provides new anatomical evidence for a direct pathway linking the SCN to the dorsal hippocampus. Anterograde labeling with rVSV-(VSV-G)-Venus revealed robust and specific terminal projections in CA2, CA3, and the DG. CA2 emerged as a primary entry point, consistent with its unique role in temporal coding and its established anatomical connectivity with septal and hypothalamic regions (Mankin et al., 2015). Hippocampal labeling was laminar-specific, with dense fibers seen in stratum oriens—a layer enriched with long-range projecting interneurons and septal fibers regulating hippocampal excitability (Freund and Buzsaki, 1996). Additional terminals were observed in stratum pyramidale, stratum lacunosum-moleculare, and dentate subregions. Immunohistochemistry revealed that most Venus-positive neurons were GABAergic (GAD65/67+), with a small subset of neurons being glutamatergic (CamKII+) in CA fields and the dentate hilus. Venus-labeled SCN axons were frequently opposed by ChAT-positive cholinergic boutons, particularly in stratum oriens, supporting the convergence of circadian and neuromodulatory influences on hippocampal excitability, and consistent with the observation that circadian disruption interacts with the cholinergic system to influence hippocampal function (Smarr et al., 2014b).

Retrograde viral tracing confirmed this connectivity by labeling SCN neurons after hippocampal injections, ruling out nonspecific spread from adjacent hypothalamic regions. Complementary monosynaptic BDA tracing also supported these direct SCN-hippocampal projections, revealing terminals in hippocampal subfields and sparse labeling in the medial septum. Together, these approaches provide convergent evidence for a sparse but reproducible direct SCN–hippocampus projection (**Figure 9**).

**Figure 9.**
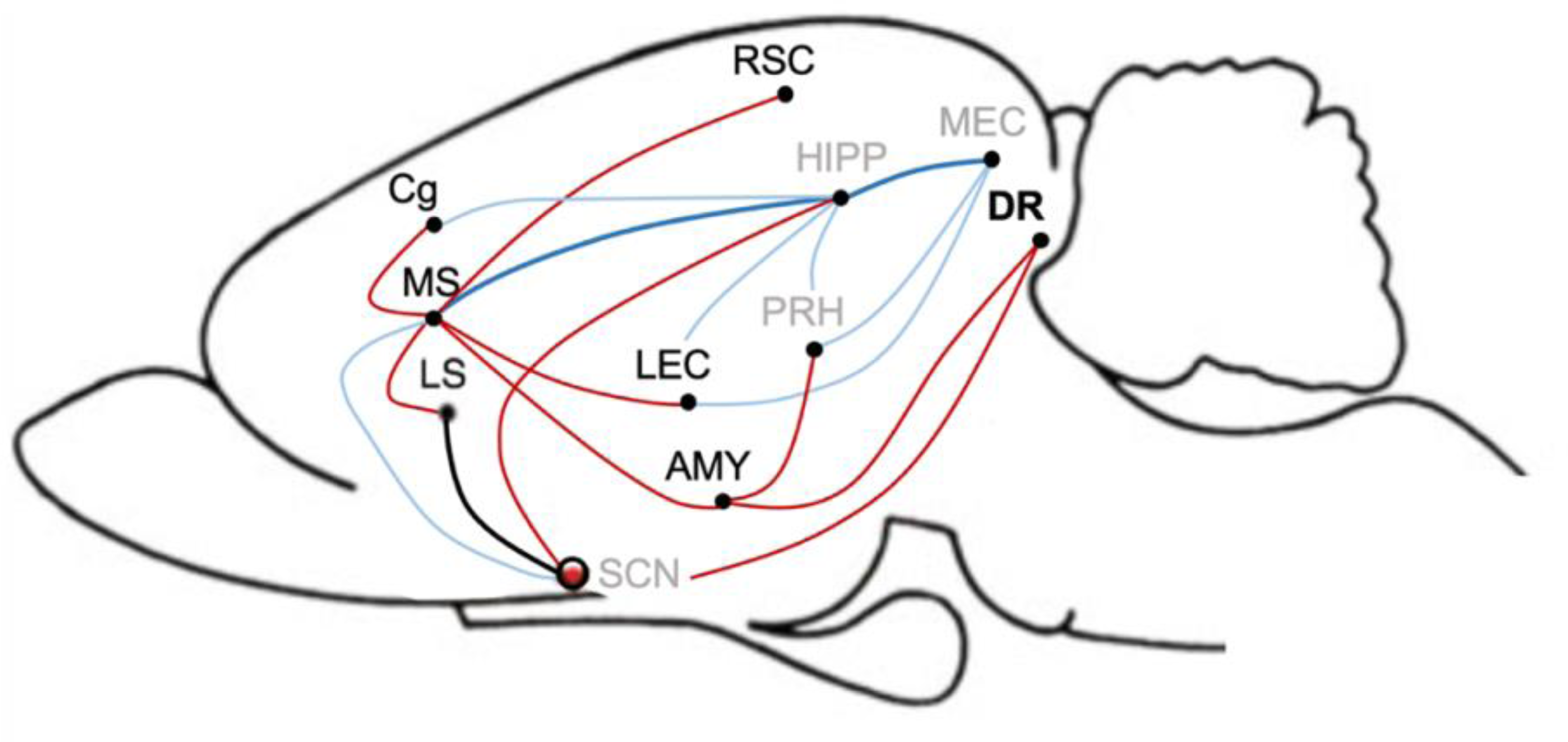
Summary model of SCN-centered circuitry linking circadian and cognitive networks. Schematic representation of the anatomical and functional circuitry linking the suprachiasmatic nucleus (SCN) with hippocampal and associated cognitive networks. Red pathways depict SCN efferent projections identified in this study, including direct and indirect connections to limbic and neuromodulatory regions such as the medial septum (MS), lateral septum (LS), amygdala (AMY), dorsal raphe (DR), and hippocampus (HIPP). Blue pathways indicate established afferent and modulatory inputs to the hippocampus from septal, cortical, and neuromodulatory regions, including MS and DR, based on prior literature. Cortical and parahippocampal regions involved in spatial and mnemonic processing are shown in grey, including retrosplenial cortex (RSC), cingulate cortex (Cg), lateral entorhinal cortex (LEC), medial entorhinal cortex (MEC), and perirhinal cortex (PRH). Together, the model illustrates how circadian signals originating in the SCN may influence hippocampal function both directly and via polysynaptic pathways, providing a framework for understanding how chronic circadian disruption can alter hippocampal network dynamics and cognitive performance.

Our findings may highlight why earlier studies failed to detect these projections. As Morin (2014) noted, even small methodological differences can produce false negatives in sparse connections. Moreover, the SCN is neurochemically heterogeneous; only specific subpopulations may project to the hippocampus, requiring cell-type–specific tools for detection. Modern viral and genetic tracing tools now provide the resolution and sensitivity to reveal such projections.

### Disrupted Functional Connectivity Underlies Selective Cognitive Deficits in CCD

Chronic manipulation of the light-dark cycle in our study caused significant alterations in both regional brain metabolism and inter-regional FC. Although whole-brain glucose uptake remained stable, hypometabolism emerged in the SCN and DG, as well as in the medial entorhinal and perirhinal cortices. FC analyses revealed loss of SCN–hippocampus connections under circadian disturbance, replaced by new, compensatory FC involving cortical and subcortical regions. Over 170 functional connections overall were found to be altered, with ∼40% involving hippocampal or cortical regions. In contrast to the more widespread regional hypometabolism seen, a selective hypermetabolism was seen in the serotonergic DR of CCD animals. This was accompanied by increased FC of the DR to the SCN, regions of the PFC and hippocampus. The suggests abnormal serotonergic modulation of this neurocircuitry in CCD animals, consistent with the known impact of circadian disturbance on serotonin system function, the role of serotonin in the regulation of circadian rhythms and the influence of the DR on SCN function (Glass et al. 2000). Thus, overall, our data suggest that circadian disturbance has a profound influence on brain function and network connectivity.

Behaviorally, deficits were specific, with object recognition memory being impaired while spatial memory, working memory and reversal learning remained intact. This coincided with disrupted connectivity and hypometabolism in perirhinal and entorhinal cortices—regions central to object recognition (Wang et al., 2021). In addition, PFC-hippocampal disturbed FC was also evident in CCD animals, which is of interest given the established role of this circuitry in object recognition (Wang et al., 2021). Impaired object recognition memory in rodents has been reported following a range of other circadian manipulations, and the task is sensitive to circadian variation, suggesting that this memory domain may be particularly sensitive to circadian dysfunction (Pugliane et al., 2025). Pattern separation also trended toward impairment, consistent with dentate gyrus hypometabolism, which further support that circadian misalignment selectively impacts recognition and pattern separation processes [see Gerstner & Yin (2010)]

Interestingly, a key role for serotonergic and DR function is supported in object recognition memory (Bouet et al., 2018), thus the observed DR dysfunction in CCD animals may contribute to this deficit. Moreover, serotonergic transmission in the PFC and hippocampus directly impacts on object recognition memory performance (Bekinschtein et al. 2013), supporting the suggestion that the altered FC between the DR and these systems may contribute to the object recognition memory deficit seen in CCD animals. Further experimental work is needed to fully elucidate the role of abnormal DR dysfunction to the cognitive deficits seen following circadian disruption.

Interestingly, LTP and LTD at the Schaffer Collateral-CA1 synapse were preserved in CCD animals, suggesting that cognitive impairments arise not from synaptic plasticity failure, but from disrupted FC across the distributed brain networks that contribute to object recognition performance. The SCN– MS–hippocampus axis may be particularly relevant here, as hippocampal theta oscillations crucial for memory are modulated by MS inputs (Buzsáki, 2002). Disrupted SCN connectivity to both MS and hippocampus, as seen in CCD animals, could desynchronize hippocampal theta rhythms, impairing hippocampal–cortical communication despite intact synaptic machinery. This hypothesis aligns with reports that theta rhythm co-ordination is influenced by circadian timing during object exploration (Lowke et al. 2020). Therefore, the potential contribution of impaired hippocampal theta rhythm activity to the object recognition deficits seen following the CCD protocol employed in our study, and the contribution of dysfunctional SCN-MS-hippocampus connectivity to these processes, certainly warrants further systematic investigation.

Overall our results extend current models of how circadian and cognitive neural systems integrate to influence cognition and behaviour. Our data show that the SCN influences hippocampal function via both direct and indirect pathways. Indirect routes, through the medial septum or paraventricular nucleus, and via impacts on DR function, provide modulatory and integrative control. In parallel, a sparse direct SCN–hippocampus projection, regulated by circadian disturbances, could deliver high- precision circadian timing cues to specific subfields such as CA2 and DG to regulate cognition. These findings have implications for understanding how shift work, jet lag, or artificial light exposure may impact cognition in modern society. Moreover, they highlight the need for interventional strategies that target network-level synchrony, such as timed light exposure, chronopharmacology or neuromodulatory interventions, to mitigate the cognitive effects of circadian disturbance.

By revealing a direct structural SCN–hippocampal projection and showing how its disturbance leads to memory-specific deficits, this study contributes to a new mechanistic framework for how circadian systems interface with higher-order cognition. Future work may further explore how these circuits develop, adapt, or deteriorate across lifespan and pathology, and how their synchronization can be restored to protect cognitive function.

## Author contributions

Inês Marques-Morgado has designed the experiments, written the draft, performed all the experiments and data analysis at Gulbenkian Institute for Molecular Medicine (GIMM), except where otherwise stated. Marcelo Dias and Joana Ribeiro assisted with the tracing injections. Marcelo Dias also assisted with the set-up of the activity monitoring recording system and wrote the scripts for activity data processing. Carolina Peixoto performed the plotting of the activity data into actograms. Joana Coelho assisted with the ^14^C-2-Deoxyglucose injections and animal dissection. The brain tissue labeled with ^14^C-2-Deoxyglucose was processed and autoradiographed by Inês Marques-Morgado at Lancaster University, Lancaster, United Kingdom, in a Short-Term scientific mission, under the supervision of Neil Dawson. Neil Dawson (Lancaster University) performed the PLSR analysis with the ^14^C-2- Deoxyglucose metabolic data. Mariana Temido-Ferreira performed the field potential electrophysiology recordings. Miguel Remondes and Luísa V Lopes coordinated the project. All authors have revised the manuscript, discussed the experimental findings, and approved the manuscript.

## Funding and acknowledgements

Marques -Morgado is an FCT Fellow (NeurULisboa PhD program), this work was funded by the Grant BIA 135/18 from Bial Foundation, by the EU Horizon 2020 Twinning SynaNet and by Fundação para a Ciência e a Tecnologia. We are indebted to Cátia Reis, Luisa Pilz and Samuel Deslauriers-Gauthier for helpful discussions and inputs. We would also like to acknowledge the Rodent, Histopathology and Bioimaging Facilities of GIMM for their technical support.

## METHODS

### Animals and housing

Animal procedures were performed in accordance with the European Community guidelines (Directive 2010/63/EU), Portuguese law on animal care (DL 113/2013). All animal research projects were reviewed by the Animal Welfare Bodies (ORBEA) of the Gulbenkian Institute of Molecular Medicine (GIMM) and approved by Portuguese Animal Ethics Committee (Direção-Geral de Veterinária), to ensure animal use was in accordance with the applicable legislation and following the 3R’s principle. Male Sprague-Dawley rats (Charles River Laboratories, France) were used for all experiments. Standard environmental conditions were kept constant: food and water ad libitum, 22- 24°C, 45-65% relative humidity, 12h light/12 dark cycles (lights on 8 am), and housed in groups of five, except where otherwise stated.

### Transection of the Sagittal Sinus

Rats were anaesthetized and securely placed on a stereotaxic frame. After exposing the skull, bregma and lambda were identified and used to level the animal’s head. Using a small drill-bit (Fine Science Tools, catalog #19007-05) a craniotomy window was gradually and carefully opened until a single bone flap is loosened (as described in Dias, Morgado et al., 2021). The flap was lifted on one side with fine tweezers and with the help of a small, blunt instrument (e.g., the small tip of a battery- powered cauterizer; Fine Science Tools, catalog #18010-00) the dura mater was carefully detached from the underside of the bone piece (blunt dissection). After exposure, the dura mater was punctured with a small needle or sharp tweezers and gradually opened from the lateral borders of the craniotomy toward the superior sagital sinus (SSS). The SSS was then lifted by the loose dural flaps, and with the help of a ligation aid (Fine Science Tools, catalog #18062-12) a 1.5 absorbable surgical suture (10 cm long) was threaded under the sinus. The suture loop was cut and each of the two remaining threads was gently pulled toward the anterior and posterior borders of the craniotomy. Here, the SSS was securely ligated and blood flow was interrupted. Finally, the SSS was cauterized between the ligations and retracted to maximally expose the underlying cortex and the longitudinal fissure. This surgical procedure allows the simultaneous bilateral targeting of midline structures. In control animals, without SSS transection, all surgical steps were equal to the experimental group, except the ligation and sectioning of the sinus. In the sham surgery, the suture was threaded under the sinus as described above and removed shortly after, without ligation. Materials and step-by-step surgical procedure are described in Detailed Protocol (https://imm.medicina.ulisboa.pt/intranet/links/view_file/TAS66TLXI/2633

### Microinjection of viruses and tracer

For the transsynaptic tracing experiments, after SSS transection was performed, animals were injected either with a transsynaptic anterograde tracer [rVSV-(VSV-G)-Venus] (Salk Institute for Biological Studies) in the SCN (100nl from 2x10^9^ ffu/ml stock) or a transsynaptic retrograde tracer [rVSV(RABV-G)-eGFP] (Salk Institute for Biological Studies) in the Hippocampus (400nl from 8.6x10^7^ ffu/ml stock), using a microinjection control system attached to the stereotaxic frame and a glass micropipette. The same procedure was used for the monosynaptic dissection of the circuit. For this experiment, animals were injected in the SCN with 100nl of 10% Biotinylated Dextran Amine- Texas Red conjugated 10,000 MW (SP-1140, Vector Laboratories) diluted in PBS. Briefly, the micropipette was filled with mineral oil and placed in the microinjector, after which the virus or tracer were aspirated to fill the tip of the pipette. Using bregma as a reference, the pipette was carefully positioned over the microinjection coordinates, slowly lowered until reaching the targets suprachiasmatic nucleus (SCN): -0.60 mm AP, ±0.3 mm ML, -9.2 mm DV; (dorsal hippocampal CA1): -3.60 mm AP, ±3 mm ML, -2.00 mm and left in place for 5 minutes. The virus/tracer was then delivered at a rate of 100 nl/min. The micropipette was then slowly retracted after waiting 10 minutes, to avoid contamination of overlaying structures. The craniotomy was then covered with a single drop of 1.5% agarose at body temperature (37°C). The skin incision was sutured and Ringer’s lactate was administered for hydration. For postoperative recovery, the animals were placed in a heated cage for 24 h, before being returned to a standard home cage. During this period, ad libitum food, nutritional gel, and water were provided.

### Transsynaptic Anatomical Tracing

Three to six days (anterograde transsynaptic tracer) or five days (retrograde transsynaptic tracer) post injection, animals were euthanized and transcardially perfused with ± 300 ml of PBS followed by ± 500 ml of 10% neutral buffered formalin. Brains were gently dissected from the skull to avoid damage to the optic chiasm and SCN, kept in formalin for 24 h at room temperature, then placed in 15% sucrose in PBS until sinking, followed by submersion in 30% sucrose in PBS. After sinking, the brains were embedded in gelatin and frozen. Frozen brains were sectioned coronally in 50μm slices using a cryostat (LEICA, CM3050 S) and virus expression was assessed using a Zeiss Axio Zoom V16 Observer fluorescence stereo microscope.

### Monossynaptic Anatomical Tracing

Eleven days after Biotinylated Dextran Amine-Texas Red injection in the SCN, animals were euthanized and transcardially perfused with ± 300 ml of PBS followed by ± 500 ml of 10% neutral buffered formalin. Brains were gently dissected from the skull to avoid damage to the optic chiasm and SCN, then kept in formalin for 24 h at RT. After sinking, the brains were embedded in gelatin . Brains were then sectioned coronally in 100μm slices using a vibratome (LEICA, VT1200 S). The tracer expression was then assessed using a Zeiss Axio Zoom V16 Observer fluorescence stereo microscope.

### Immunohistochemistry

Free-floating coronal brain slices were incubated with PBS at 37°C for 10 minutes to remove the gelatin and subsequently incubated with PBS + 0.1M glycine for 20 minutes for quenching. For blocking and permeabilization, slices were incubated in TBS + 0.5% Triton X-100 + 10% FBS at RT on a shaker. Incubation with primary antibodies was performed overnight with rotation at 4°C, with antibodies diluted in TBS + 0.2% Triton X-100, FBS 4% (Table below). Following three washes in TBS + 0.1% Triton X-100, slices were then incubated with secondary antibodies overnight with rotation at 4°C, with antibodies diluted in TBS + 0.2% Triton X-100, FBS 4%. After incubation, samples were washed three times in TBS + 0.1% Triton X-100, slices were incubated with Hoechst 33258 (12 ug/mL in PBS, Life Technologies) for 10 minutes, washed once in PBS, mounted in slides with Fluoromount aqueous mounting medium (Sigma), coverslipped and left to dry in the dark.

### Primary and secondary antibodies and related conditions used in immunohistochemistry experiments

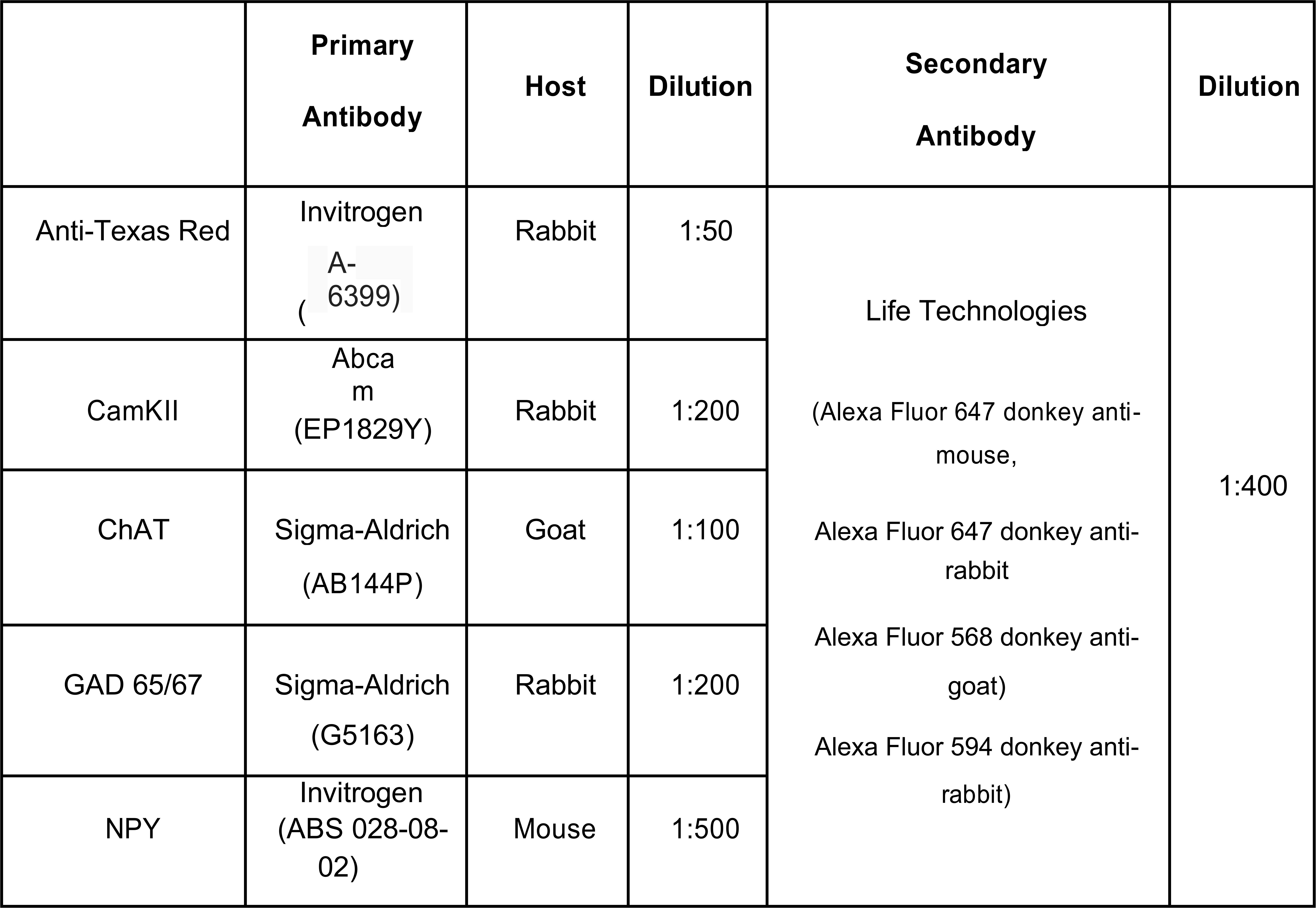

All antibodies were diluted in TBS + 0.2% Triton X-100, FBS 4%. Abbreviations: CamKII - Ca2+/calmodulin-dependent protein kinase; ChAT - Choline acetyltransferase; GAD 65/67 - Glutamic Acid Decarboxylase 65/67; NPY - Neuropeptide Y.

### Microscopy imaging and analysis

Z-stack images were acquired using a Zeiss LSM 880 Point Scanning Confocal Microscope equipped with a GaAsP detector for increased sensitivity. Z-stack images were acquired with 20x (Plan-Apochromat, numerical aperture 0.80, working distance 0.55mm) 40x (C-Apochromat Corr, numerical aperture 1.20, working distance 0.28mm) and 63x objectives (Plan-Apochromat DIC, numerical aperture 1.40, working distance 0.19mm).

### Chronic Circadian Manipulation

Twenty 5 weeks-old Sprague-Dawley rats (Charles River) were used in this experiment and divided into phase-shifted group (n=10) and control group (n=10). Animals were housed in groups of five. Three additional animals from each group were single housed for representative activity monitoring. The chronic circadian manipulation was adapted from (Craig & McDonald, 2008). Briefly, a two- stage phase-shifting manipulation was implemented, consisting of a shifting phase - in which the light/dark cycle was advanced by 3 hour/day for 6 consecutive days, followed by a recovery period in which the light/dark cycle was kept constant (12:12) for 8 consecutive days. This dual strategy was repeated for four times, allowing a continuous challenge of the circadian system throughout the duration of the experiment **(according to the schedule below)**. Control rats remained on a constant 12 h light/dark cycle (light on 8 am) for the experiment. To assess the consequences of active photoperiod shifting, after the fourth recovery period, shifted animals started a new phase shifting. Two days after the beginning of this last shifting phase (day 54), brain metabolism was evaluated.

### Schedule of Chronic Manipulation of Circadian Rhythms

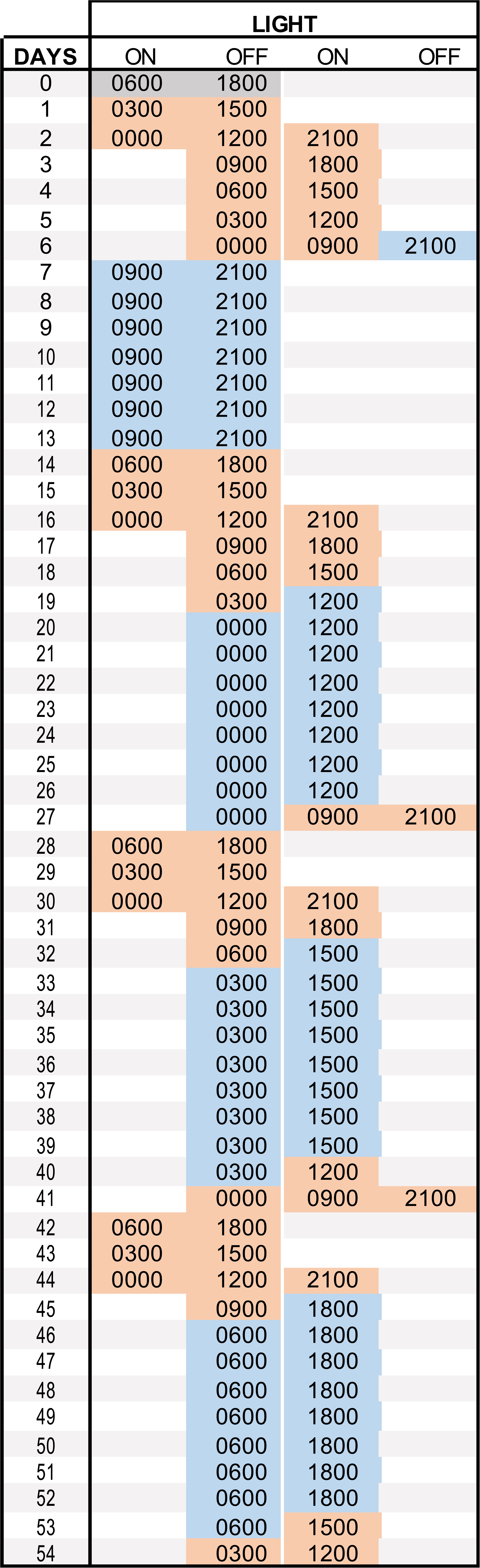

*While control animals were kept under a normal 12:12h light/dark (LD) cycle (Colony LD), the shifted animals were subjected to 4 cycles of phase shifts (orange to blue transitions) followed by stable 12:12h LD cycle to continually challenge the entrainment properties of the circadian system. After the fourth recovery period, a new shifting phase was initiated. Two days after the beginning of this last shifting phase, brain FC was assessed. Three animals from each experimental group were single housed for representative activity monitoring. After 54 days, after the completion of the Chronic Manipulation of Circadian Rhythms protocol, these animals were released into constant darkness for 15 days to analyze their free-running rhythm*.

### Video-Based Animal Activity Tracking

Rats were individually housed in acrylic arenas (40 × 50 × 40 cm). Movements were recorded non- invasively with ELP 1MP USB dome cameras with integrated infrared (IR) LEDs, mounted at 50 cm above the arena center with wide-angle lenses to capture the full arena under visible light or infrared illumination.

Video was acquired at 1 fps directly in Bonsai software (v2.3, https://bonsai-rx.org/), using the *Bonsai.Vision* package. The workflow consisted of two parallel processing branches that ran continuously, each optimized for a different lighting condition and each scheduled to run for 12 h: one for lights ON (day) and one for lights OFF (IR/night). In the lights ON condition, intensity-threshold segmentation was used, whereas as in the lights OFF condition, background subtraction was applied, involving the subtraction of a background image of the cage without the animal present from the recorded video images, using the contrast difference between the rat and the background.

After segmentation, connected-component analysis was performed on the binary image so that the largest detected object was taken as the animal and its centroid (X,Y) was recorded per frame with synchronized timestamps. Ambient light was monitored with an Arduino with a light sensor connected to Bonsai, in which the signal was thresholded to label lights ON/OFF with True/False, accordingly.

For each 24-h cycle the software produced four CSV files (two optimized for lights-on, two for lights- off) with five outputs: absolute clock time measured as .NET ticks timestamp, centroid X position, centroid Y position, human-readable timestamp, and light state.

CSV files exported from Bonsai were then processed in MATLAB software. For each 24-h session, only rows whose lighting state matched the branch were kept: rows from night-optimized files were retained only when Light = False, and rows from day-optimized files only when Light = True. The selected rows were then concatenated into a single CSV file and sorted by the absolute timestamp.

To ensure uniform sampling, timestamps were curated to have a fixed interval of one second between observations using R software. A complete sequence of one-second intervals spanning the first to the last timestamp was generated and merged with the raw data. Missing numeric values (X and Y coordinates) were imputed as the mean of the preceding and following timepoints, while missing light values were replaced with the value from the preceding timepoint. This procedure produced a sequential dataset with one-second interval. Position coordinates reflecting tracking errors outside the boundaries of the arenas were replaced with NA.

To calculate velocity, X and Y coordinates were converted to centimeters using the real size and image maze dimensions:

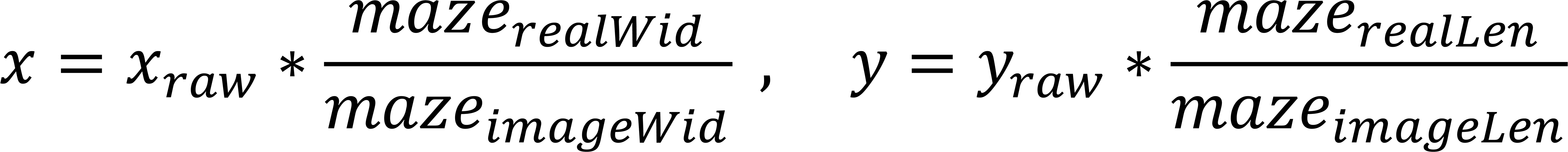

A moving median filter was applied to smooth the position signals. Instantaneous changes in X and Y directions were calculated as the difference between consecutive timepoints (Δ𝑥 = 𝑥_𝑡+1_ − 𝑥_𝑡_), and velocity (cm/s) was derived from these measures (𝑡 = 1):

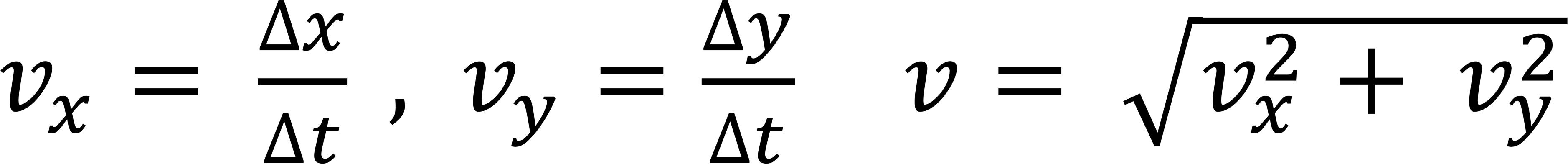

Velocity data was aggregated into one-minute bins by averaging 60 consecutive one-second measurements, based on a similar video-based tracking method (Fisher et al. 2012). Each timepoint was classified as active or inactive based on whether its velocity exceeded the overall mean velocity of the dataset.

### Actogram plotting

Binned activity data were visualized as actograms using the ***ggplot2*** package in R, with light/dark cycles indicated for reference.

### Code availability

All analyses and visualizations were performed in R. The code used for processing and plotting actograms is available upon request.

### Serum Corticosterone Measurement

At the completion of the Chronic Manipulation of Circadian Rhythms protocol, after the shifted animals had finished the fourth recovery period with a stable 12:12h light-dark cycle, blood samples were collected from the tail vein of the previously handled animals at four different timepoints without the use of anesthesia. The plasma was isolated by centrifugation at 2000 g for 15 minutes at 4°C. Serum corticosterone was measured using a Corticosterone ELISA Kit (Cat. No: EU3108, FineTest Biotech), according to the manufacturer’s instructions.

### Quantitative PCR

Liver samples were collected from the animals at the time of sacrifice and cryopreserved at -80°C. Total liver RNA was extracted and purified using the RNeasy Lipid Tissue Mini Kit (QIAGEN, Germany). RNA quality was assessed by NanoDrop 2000 (Thermo Scientific, USA) analysis (A260/A280≈2; 260/235 >1.8). Total RNA was reverse-transcribed using random primers and SuperScript First-Strand Synthesis System for RT-PCR (Invitrogen, USA). Real-time quantitative polymerase chain reaction (RT-qPCR) analysis was performed on a Corbett Rotor-gene 6000 apparatus (QIAGEN, Germany) using Power SYBR Green PCR Master Mix (Applied Biosystems, UK), 0.2 mM of each primer and 0.4 ng/μL of cDNA. The thermal cycler conditions were optimized for the genes under analysis. For the *Cyp2c7* gene, the conditions were: 10 minutes at 95°C, followed by 40 cycles of a two-step PCR consisting of 95°C for 15 seconds and 56°C for 25 seconds, concluding with a final thermal ramp from 72°C to 95°C. For the *CD36* gene, the conditions were: 10 minutes at 95°C, 40 cycles of a two-step PCR with 95°C for 15 seconds and 62°C for 60 seconds, and a final thermal ramp from 72°C to 95°C. The primers presented high amplification efficiency (>80%) with R^2^ values of standard curves around 0.99. Reference genes used for the analysys were *CypA* (cyclophilin A), *Rpl13A* (ribosomal protein L13A) and *Pgk1* (phosphoglycerate kinase 1). All amplifications were carried out in triplicates and according to the MIQE guidelines (Bustin et al., 2009). The relative expression of target genes was determined by the comparative delta-delta CT method (Schmittgen & Livak, 2008).

### Primer design

The RT-qPCR primers were designed using the Primer-Blast tool (http://www.ncbi.nlm.nih.gov/tools/primer-blast/). The design followed standard parameters, including: 1) melting temperature between 57 and 63°C, with a maximum 3°C difference between the primers, 2) primer length between 15 and 25 nucleotides, and 3) GC content between 30 and 80%. Additionally, the "Max Self Complementarity" and "3’ complementary value" were set at 8 and 3, respectively, and the PCR product size was limited to 80-120 base pairs. To avoid amplification of genomic DNA, the "Primer must span an exon-exon junction" option was selected whenever possible. For "Primer Pair Specificity Checking Parameters", the database used was "Ref Seq RNA" from the organism *Mus musculus*, and the options "Enable search for primer pairs specific to the intended PCR template" and "Allow primer to amplify mRNA splice variants" were chosen. The resulting best primer pair sequences were further analyzed for potential genomic DNA amplification and secondary structures.

### 14C-2-Deoxyglucose functional brain imaging

#### 14C-2-Deoxyglucose injection

Two days after starting a new shifting phase at the end of the chronic circadian manipulation procedure, during the light phase of the light/dark cycle (6h after light onset), animals were injected intraperitoneally with 3.7MBq/kg (100μCi/kg) of [^14^C]-2-Deoxyglucose (Cat. No: ARC 0111A, American Radiolabeled Chemicals Inc., USA) at a steady rate before being returned to their home cage. As previously described in (Hugues et al., 2020), forty-five minutes after isotope injection animals were decapitated without anesthesia and a terminal blood sample was collected in heparinized centrifuge tubes. Terminal glucose blood concentration was determined using a glucometer. Brains were gently dissected from the skull to avoid damage to the optic chiasm and SCN, frozen in isopentane (-40°C) and stored at -80°C until sectioning. Blood samples were centrifuged to separate the plasma for further determination of ^14^C concentrations by liquid scintillation analysis.

### Autoradiogram preparation and Densiometric Analysis

Frozen brains were coronally sectioned (20 µm) in a cryostat (-20°C). A series of three consecutive sections were retained from every 200 µm, except between the Preoptic Area and Retrochiasmatic Area, where all serial slices were retained to include all hypothalamic nuclei in study. Slices were thaw mounted onto slide covers and rapidly dried on a hot plate (70°C). Autoradiograms were generated by apposing these sections, together with pre-calibrated ^14^C-standards (40–1069 nCi/g tissue equivalents; ARC in, USA) to X-ray film (Kodak, Biomax MR for 10 days. Autoradiographic images were analyzed by a computer-based image analysis system (MCID/M5+). The local isotope concentration for each brain ROI was derived from the optical density (OD) of autoradiographic images relative to that of the co-exposed ^14^C standards. Twelve replicate measurements were taken from 68 anatomically distinct brain regions, defined with reference to a stereotaxic rat brain atlas (Paxinos & Watson, 2007) **(see Diagram below)**. The rate of metabolism, Local Cerebral Glucose Utilization (LCGU), in each ROI was determined as the ratio of ^14^C present in that region relative to the average ^14^C concentration in the whole brain of the same animal. Whole brain average ^14^C levels were determined from the average ^14^C concentration across all sections in which a ROI was measured. Significant alterations in LCGU were analyzed using Student’s t-test and significance was set at p=0.05. For regional LCGU data, effect sizes (mean and 95% confidence interval: CI) were also estimated, through 5000 permutations of the data (Figure 4).

### Functional Brain Connectivity Analysis

Regional FC was analysed in control and phase-shifted animals as previously described in Mouro et al. 2018. The Partial Least Squares Regression (PLSR) algorithm (pls package in R, https://CRAN.R-project.org/package=pls) was used to characterize statistically significant differences in the FC of defined ‘seed regions’ to all the other ROI analysed. In our analysis, the seed regions were selected based on significant differences in LCGU observed upon chronic manipulation of circadian rhythms, as well as the results from anatomical tracing experiments. The selected seed brain regions were the Suprachiasmatic Nucleus (SCN), the rostrodorsal and intermediate Dentate Gyrus (DG), the Medial Entorhinal Cortex (MEC), the Perirhinal Cortex (Prh), the Dorsal Raphé (DR), Medial Septum (MS), and the Paraventricular Hypothalamic Nucleus (PVN). The application of the PLSR algorithm to functional ^14^C-2-DG imaging data was performed as previously reported (Dawson et al., 2012a; Mouro et al., 2018). PLSR models were generated independently for each experimental group. The FC between each seed region and all other ROI measured was defined by the variable importance to the projection (VIP) statistic obtained from PLSR analysis. A significant functional connection between brain regions was considered to exist if the 95% confidence interval (CI) of the VIP statistic exceeded 1.0, denoting a considerable contribution of the explanatory variable (ROI) to the dependent variable (seed region) in PLSR analysis. The Standard Deviation (SD) and CI of the VIP statistic were estimated by jack-knifing. The significance of the alterations in the VIP statistic was determined by unpaired Student’s t-test with post hoc Bonferroni correction for multiple comparisons. Significant differences between experimental groups were set at p<0.05. Alterations were defined as being significantly lost or newly gained FC (shifted animals significantly different from controls and the 95% CI of the VIP statistic exceeds the 1.0 threshold in only one of the experimental groups).

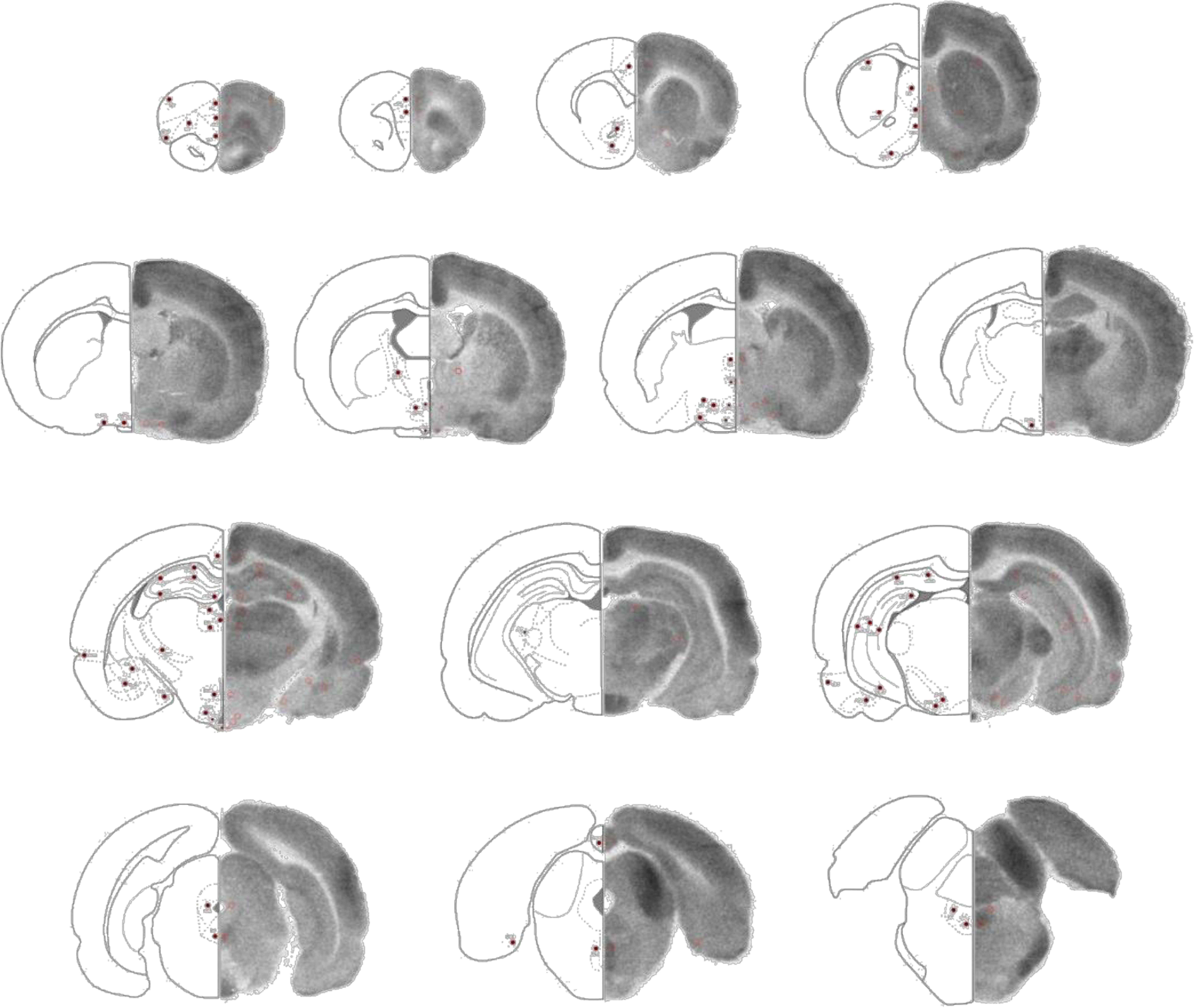

**Anatomical location of the 68 ROI** - Representative diagrams and autoradiograms depicting the anatomical localization of the 68 ROI analyzed Behavior

### Elevated Plus Maze (EPM)

All testing was conducted during the dark phase, when rodents are naturally most active. Anxiety- like behavior was analyzed using Elevated Plus Maze (EPM), as previously described (Coelho et al., 2014). The EPM maze is shaped like a plus sign and consists of two ‘open’ and two ‘closed’ arms, arranged perpendicularly, and elevated 50cm above the floor. During the test, each animal was placed at the center of the maze facing an open arm. Each test lasted 5 minutes from which the total time spent in the open arms and the total number of arm entries were used as measures of anxiety-like behavior and locomotor activity. The maze was cleaned with a 70% ethanol solution between trials.

### Open Field (OF)

Locomotor and exploratory behavior were assessed with Open field test as previously described (Coelho et al., 2014). Rats were placed at the center of a squared arena (66 x 66 cm) and freely explored during 5 minutes. Activity was recorded and analyzed using a video-tracking software (Smart 2.5, PanLab) to assess total distance traveled, average speed, resting time and permanence time in peripheral versus central portions of the arena. The maze was cleaned with a 70% ethanol solution between each animal.

### T-Maze: Spontaneous alternations

Spontaneous alternation behavior was assessed throughout the Chronic Manipulation of Circadian Rhythms, at the end of each recovery period. The test was performed as previously described in (Deacon & Rawlins, 2006). Rats are confined to the start chamber for one minute, and then permitted to access the rest of the maze for seven minutes. An alternation attempt was scored when all four feet of the rat entered one of the lateral arms, re–entered the stem arm, and then entered the lateral arm opposite the one previously chosen. Re-entry into the same arm was considered a non- alternation. Alternation performance was thus defined by the number of alternations observed divided by the number of alternation attempts × 100. To control for odor cues, the maze was cleaned with 70% ethanol between trials.

### Y-Maze: Short term reference memory

Short-term reference memory was assessed in a spontaneous novelty-based spatial preference Y- maze test as previously described (Temido-Ferreira et al., 2020). The test was performed as a two- trial recognition test in a Y-shaped maze with three arms (each with 35x10x20 cm), angled at 120° and with opaque walls. Different visual cues were placed at the end of each arm. Allocation of the arms was counterbalanced across trials. During the habituation phase (learning trial) rats were placed at the end of the “start” arm, allowed to explore two arms of the maze for 10 min and returned to the home cage. Access to the third arm (“novel” arm) was blocked by an opaque door. After 1 hour, the door of the novel arm was removed and animals were placed again in the start arm to freely explore the maze for 5 minutes (test trial). Rat tracings were recorded using a video-tracking software (Smart 2.5, PanLab). Preference for the novel arm is considered a measure of short-term reference memory. The maze was cleaned with a 70% ethanol solution between each animal.

### Morris Water Maze

Long-term spatial memory was evaluated with the Morris Water Maze test, as previously described (Batalha et al., 2013). The test was performed over the course of 5 consecutive days and consisted of a 4-day acquisition phase and a 1-day probe test. The test was performed in a circular pool (1.8 m diameter, 0.6 m height), filled with water opacified with non-toxic black paint and kept at 25 ± 2 °C. A round 8-cm in diameter platform was hidden 1 cm beneath the surface of the water at a fixed position. Four positions around the edge of the tank were used, dividing the tank into four quadrants: target quadrant (T, quadrant where the platform was hidden), left quadrant (L, quadrant on the left of the target quadrant), right quadrant (R, quadrant of the right of the target quadrant) and opposite quadrant (O, quadrant on the opposite side of the target quadrant). During the acquisition phase, each animal was given four swimming trials per day (30-mins inter-trial interval). A trial consisted of placing the animal into the water facing the outer edge of the pool and allowing the animal to explore and reach for the hidden platform. If the animal reached the platform before 60 s, it was allowed to remain there for 10 s. If the animal failed to find the target before 60 s, it was manually guided to the platform, where it was allowed to remain for 20 s. After the end of each trial, animals were removed from the pool and placed back to their home cages beneath heat lamps in order to prevent temperature loss. During the probe test, the platform was removed and animals were allowed to swim freely for 60 s while recording the percentage of time spent on each quadrant. The latency to find the platform during the acquisition phase and the percentage of time in the platform quadrant during the probe test were recorded and analyzed using the Smart 2.5 tracking system (PanLab, Barcelona) to evaluate hippocampal-dependent memory. Swimming speed was also registered, as a measure of possible motor deficits that could interfere with the ability to perform the task.

### Novel Object Recognition test (NOR)

Object recognition memory was assessed as previously reported in (Mathiasen & DiCamillo, 2010) using a square arena (66 x 66 cm). The test consisted of a sample trial in which two identical objects were placed into two specific locations of the arena and animals were allowed to freely explore the environment and the objects for 3 minutes before returning to the home cage. After a retention interval of 24 hours, animals were placed inside the arena for the choice trial, to explore a familiar object from previous trial and a novel object for 3 minutes. The designations of “familiar” and “novel,” and the novel object’s location in the test box were counterbalanced and randomly assigned to avoid any influence of object and place preference. Activity was recorded using a camera positioned on the ceiling. Exploratory behaviour was quantified as the amount of time (seconds) animals spent examining each object (placing the nose within 2 cm of the object and any of the following: obvious movement of the vibrissae, sniffing, licking, or rearing onto the object). Sitting on the object or close contact in which the nose was not directed to the object were not considered as exploration. A Discriminating Index (DI) was calculated to analyse the exploration of each object (DI = [novel (sec) – familiar (sec)] / [total novel (sec) + familiar (sec)]), using the amount of time spent exploring the novel and the familiar object during the choice trial. This index ranges from -1 to 1 (-1 = exclusive exploration of the familiar object; 0 = absence of discrimination between novel and familiar objects, i.e. equal time exploring both objects; and 1 = exploration of the novel object only). The arena and the objects were cleaned with a 70% ethanol solution between each animal.

### Object Pattern Separation Task

Pattern separation was assessed as previously reported in (van Goethem et al., 2018) using a round arena. Animals were habituated to the empty arena, followed by two trials with the two objects present. The In the learning trial (T1), two identical objects were positioned symmetrically on the midline of the arena (“position 1”), ∼25 cm from each side wall. After a 1 h inter-trial interval (ITI), the test trial (T2) was conducted with one object remaining at position 1 and the other displaced along the vertical axis to position 3 - an intermediate separation halfway between the minimal and maximal displacements commonly used in this assay. Both trials lasted 3 min.

Across animals, the displaced side (left/right) and direction (forward/backward) were counterbalanced according to a predefined randomization schedule to avoid side or direction confounds. Activity was recorded using a camera positioned on the ceiling and object exploration was scored. To ensure adequate sampling, animals were required to meet minimal total exploration time of 7 s in T1 and 10 s in T2.

Using a Discriminating Index (DI) (DI = [displaced (sec) – stationary (sec)] / [total displaced (sec) + stationary (sec)]. This index reflects preference for the novel location and ranges from -1 to 1 (-1 = exclusive exploration of the stationary object; 0 = absence of discrimination between stationary and displaced objects, i.e. equal time exploring both objects; and 1 = exploration of the displaced object only). The arena and the objects were cleaned with a 70% ethanol solution between each animal.

### Field Electrophysiological Recordings

Field electrophysiological recordings were performed as previously described (Temido-Ferreira et al., 2020). After decapitation, the brain was rapidly removed and the hippocampi were dissected free in ice-cold Krebs solu- tion, which is composed of (mM): NaCl 124; KCl 3; NaH2PO4 1.25; NaHCO3 26; MgSO4 1; CaCl2 2 and D- glucose 10, continuously gassed with 95% O2 and 5% CO2, pH 7.4. Transverse hippocampal slices with 400μm thickness were obtained with a McIlwain tissue chopper and field excitatory postsynaptic potentials (fEPSPs) were recorded in the *stratum radiatum* of the CA1 area at 32 °C. LTP was induced by a theta-burst stimulation protocol (TBS, ten trains with four pulses each at 100 Hz, separated by 200 ms) applied to the Schaffer collaterals at the test stimulus intensity (50% half-maximal fEPSP). LTD was induced using a low frequency stimulation protocol (LFS, three trains with 10-min interval of 2 Hz, 1,200 pulses) as previously described (Ferreira et al., 2017). Stimulation, data acquisition and analysis were performed using the electrophysiology software WinLTP (Molecular Devices).

### Statistical Analysis

GraphPad Prism 7 software was used for statistical analysis. The values presented are mean ±SEM of n experiments. To test the significance of differences between experimental groups, unpaired Student’s t-test or two-way ANOVA followed by a Bonferroni’s Multiple Comparison post hoc test were used. Values of P<0.05 were considered to be statistically significant.

