## Supplemental Material for "Chronic Circadian Disturbance Reshapes Hippocampal Connectivity and Cognitive Function"

**SUPP FIGURE 1**

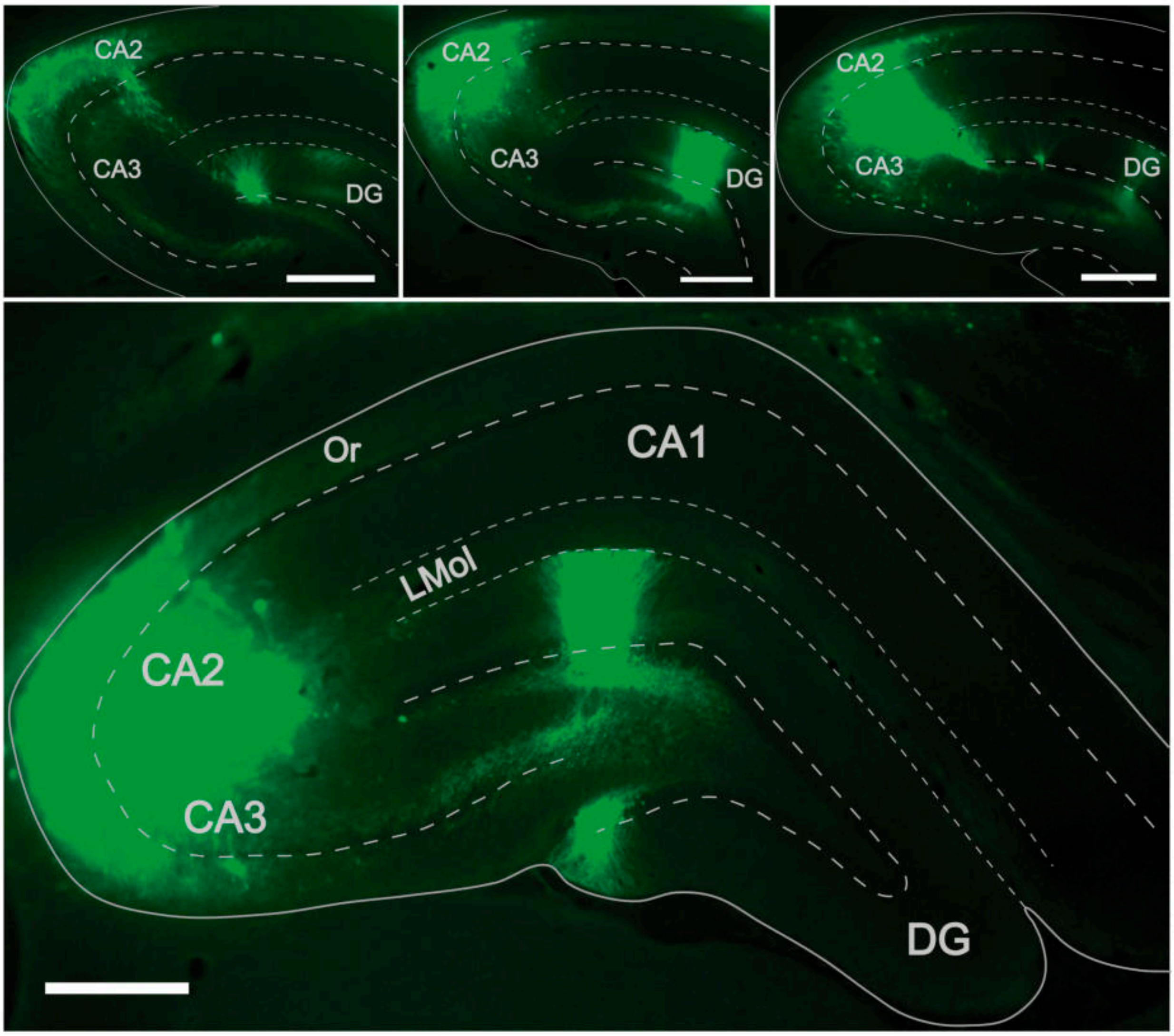

SUPP FIGURE 2

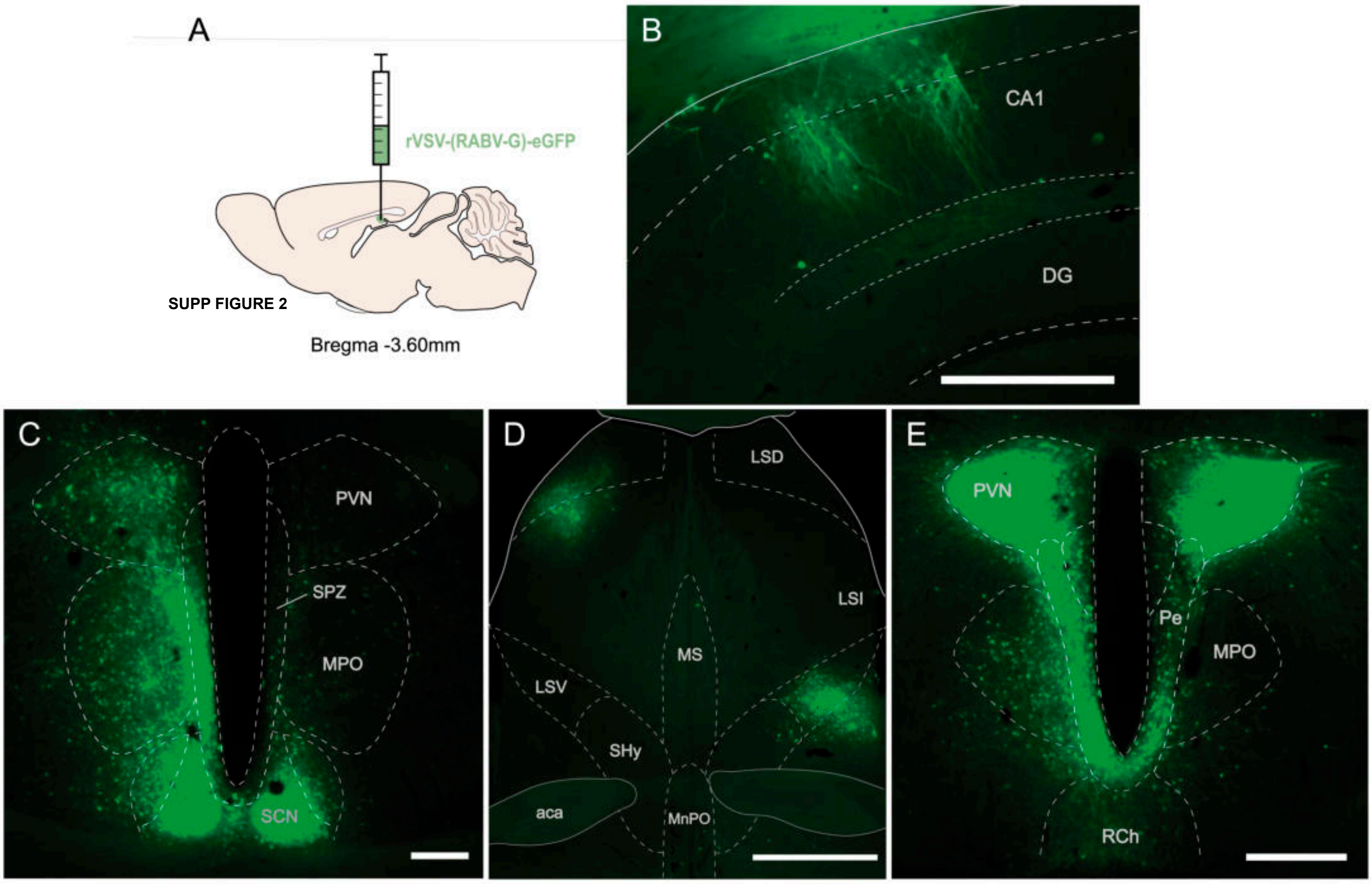


SUPP FIGURE 2

SUPP FIGURE 3

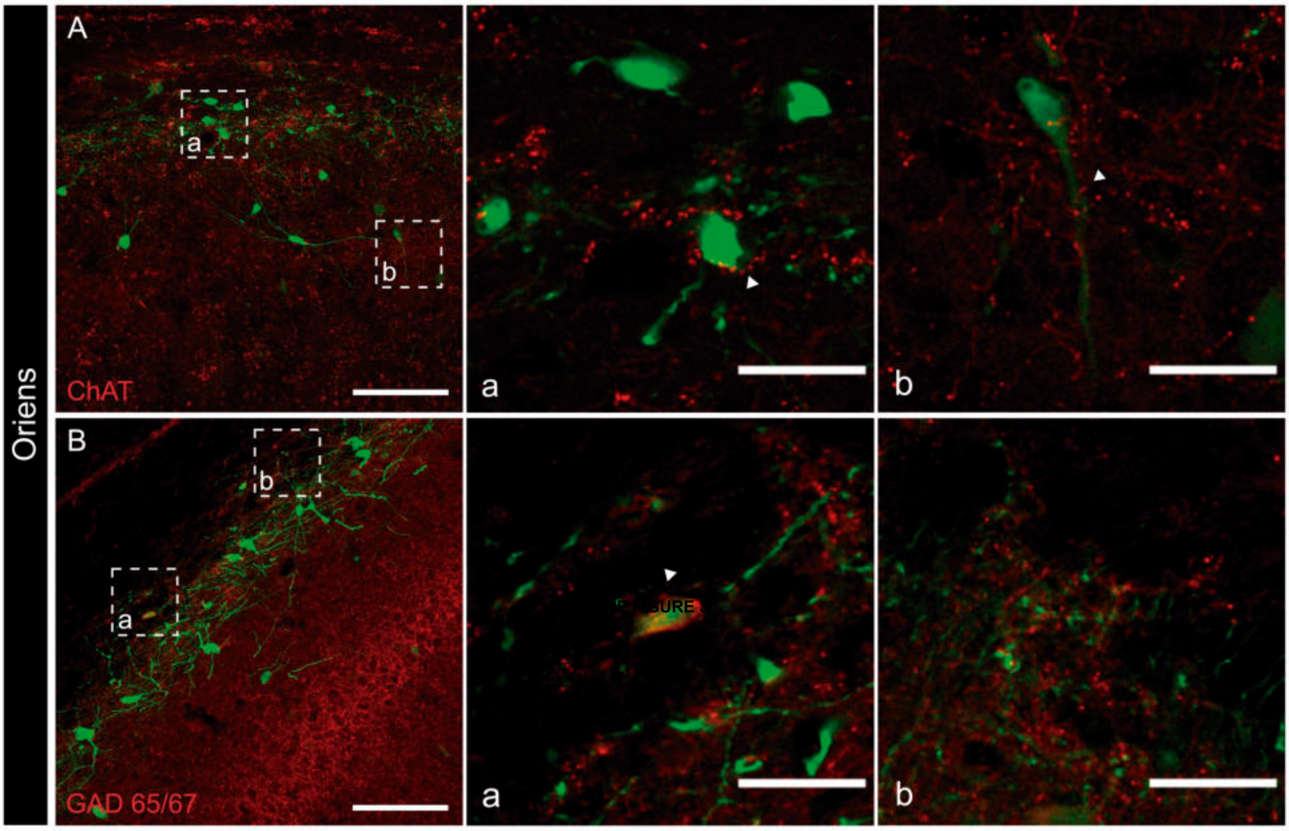

SUPP FIGURE 4

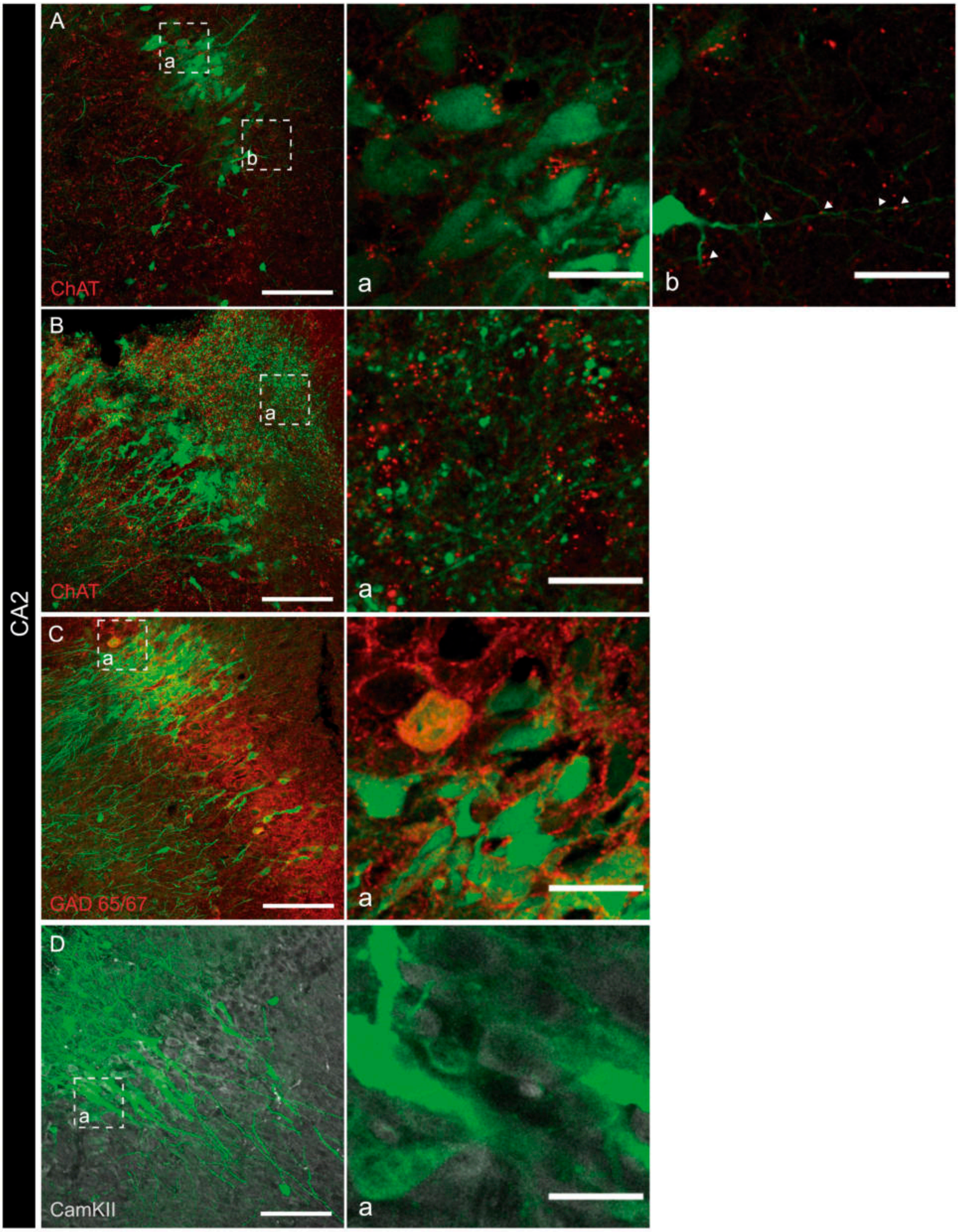

SUPP FIGURE 5

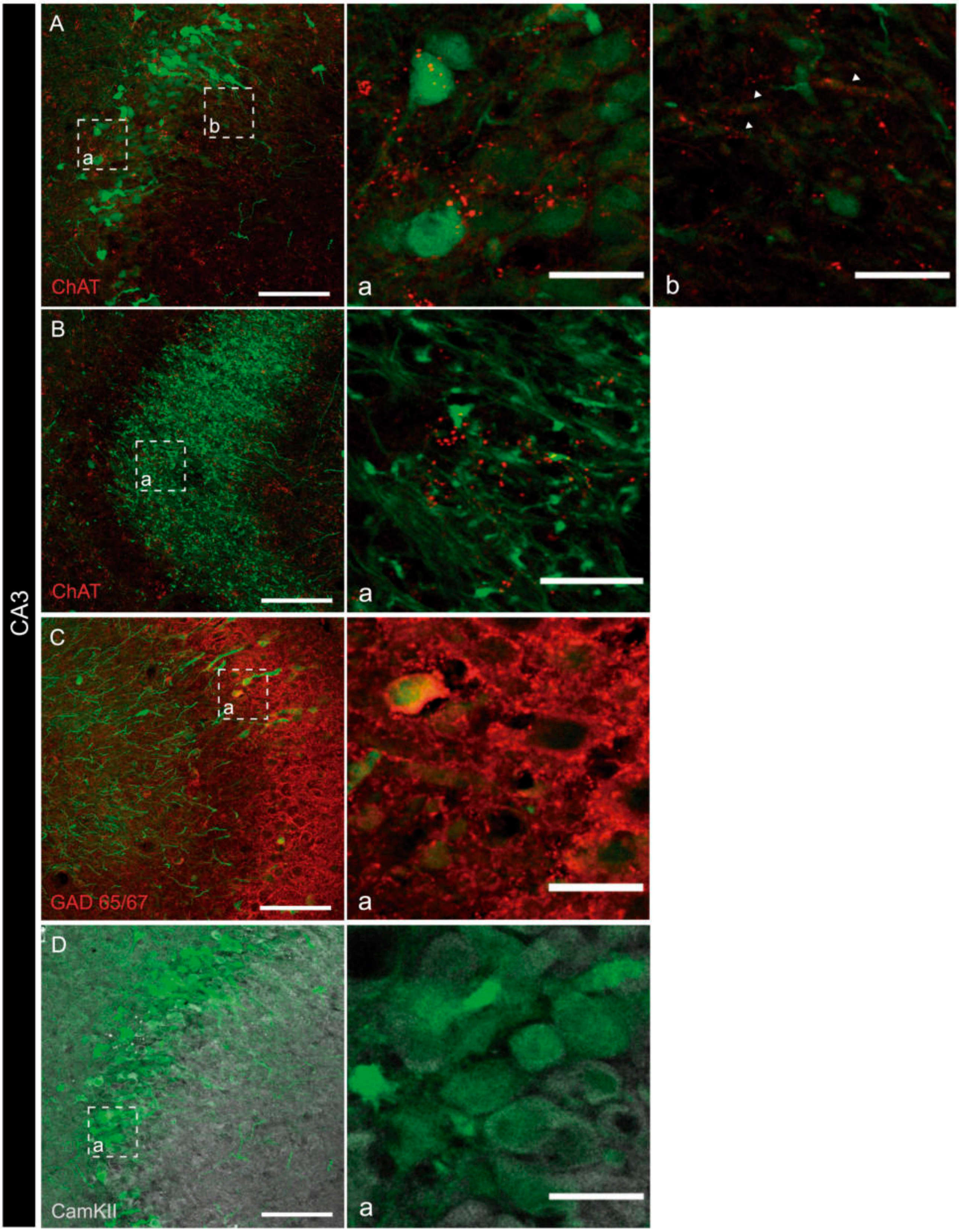

**SUPP FIGURE 6**

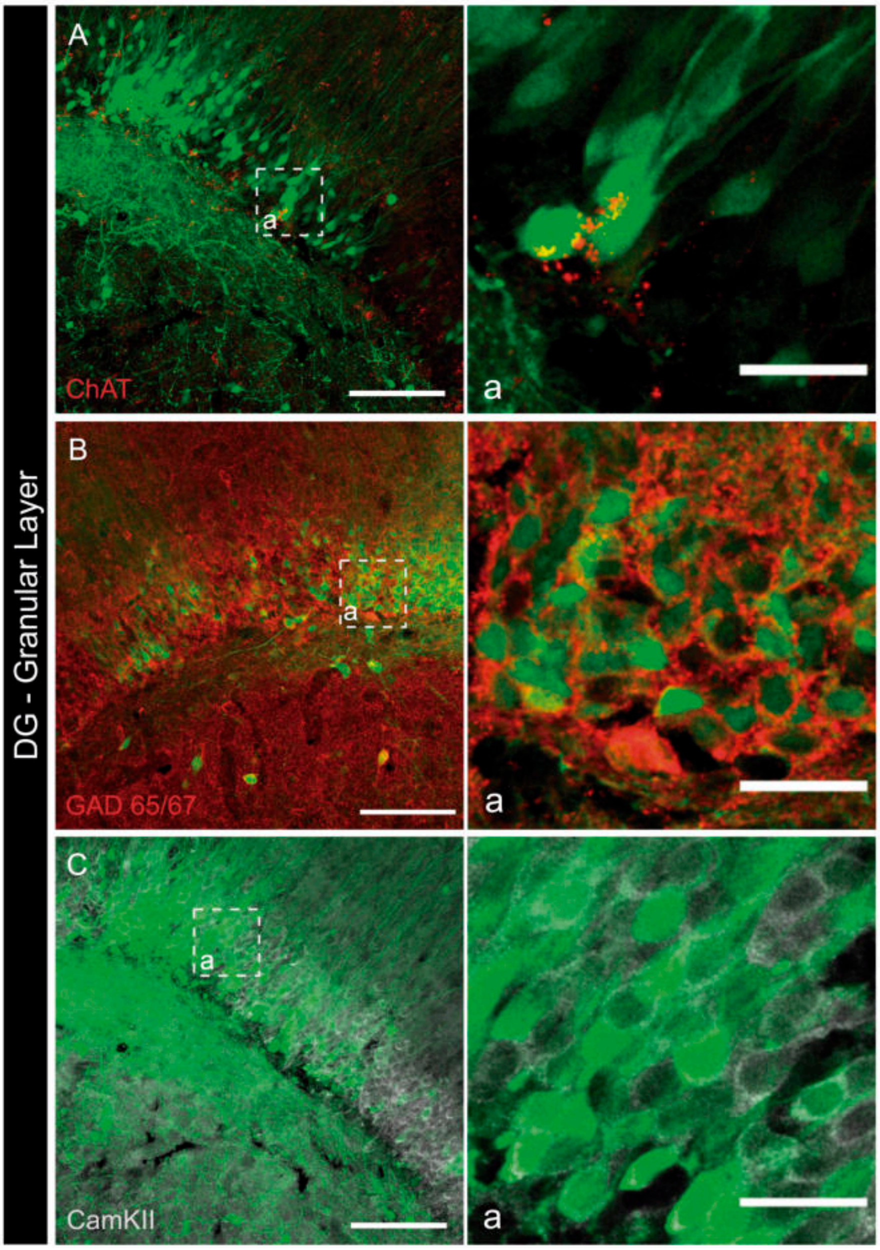

SUPP FIGURE 7

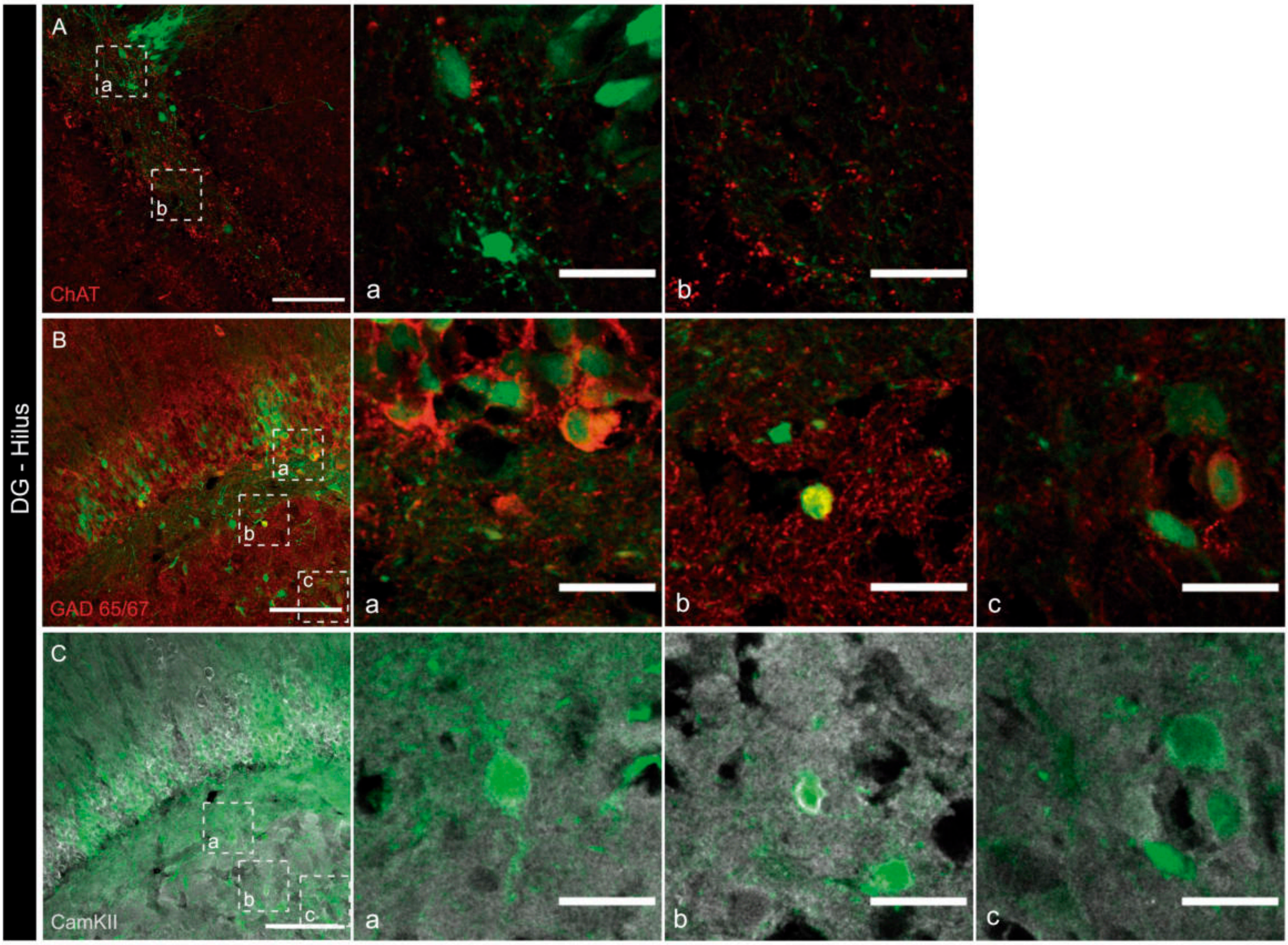

SUPP FIGURE 8

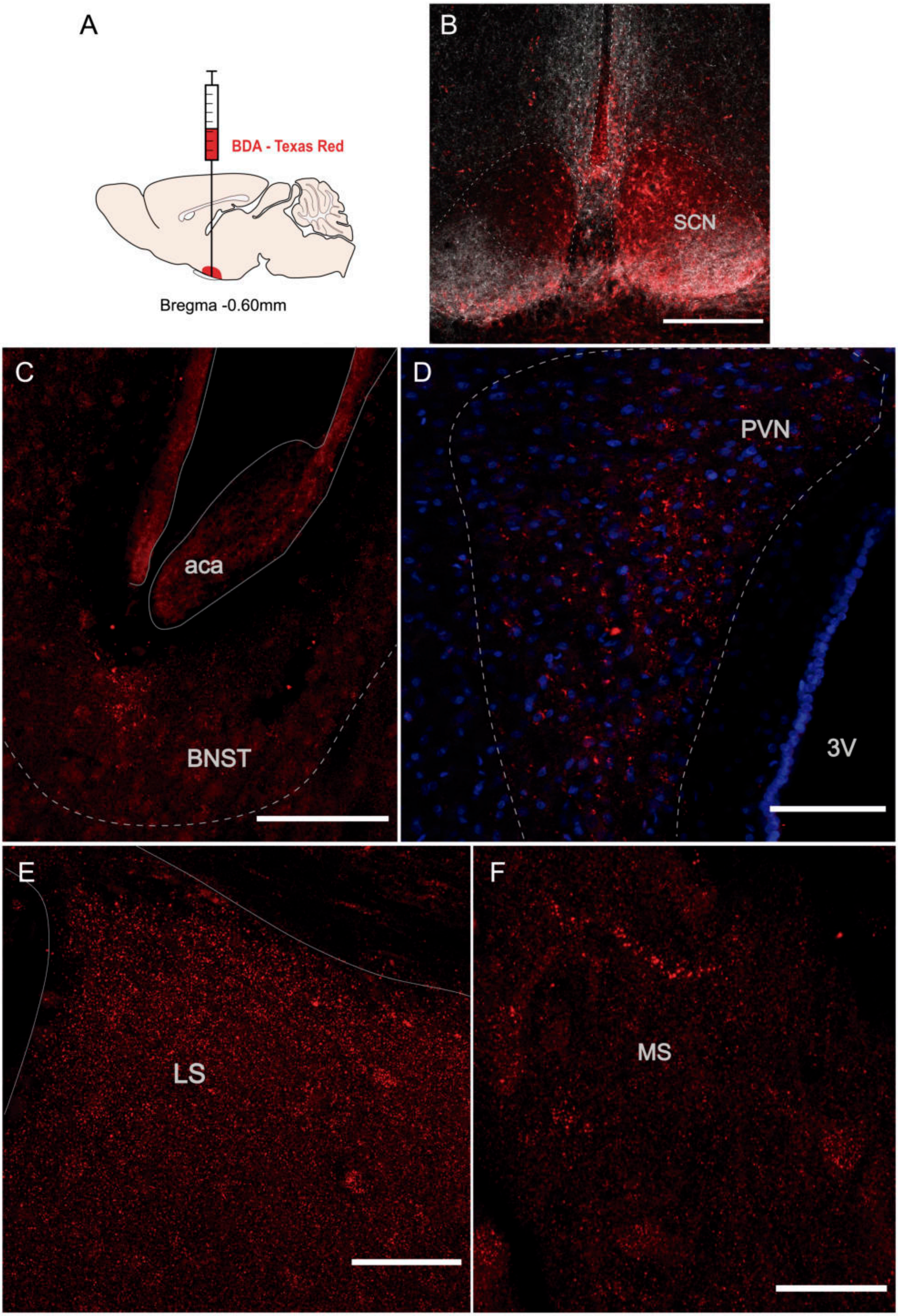

SUPP FIGURE 9

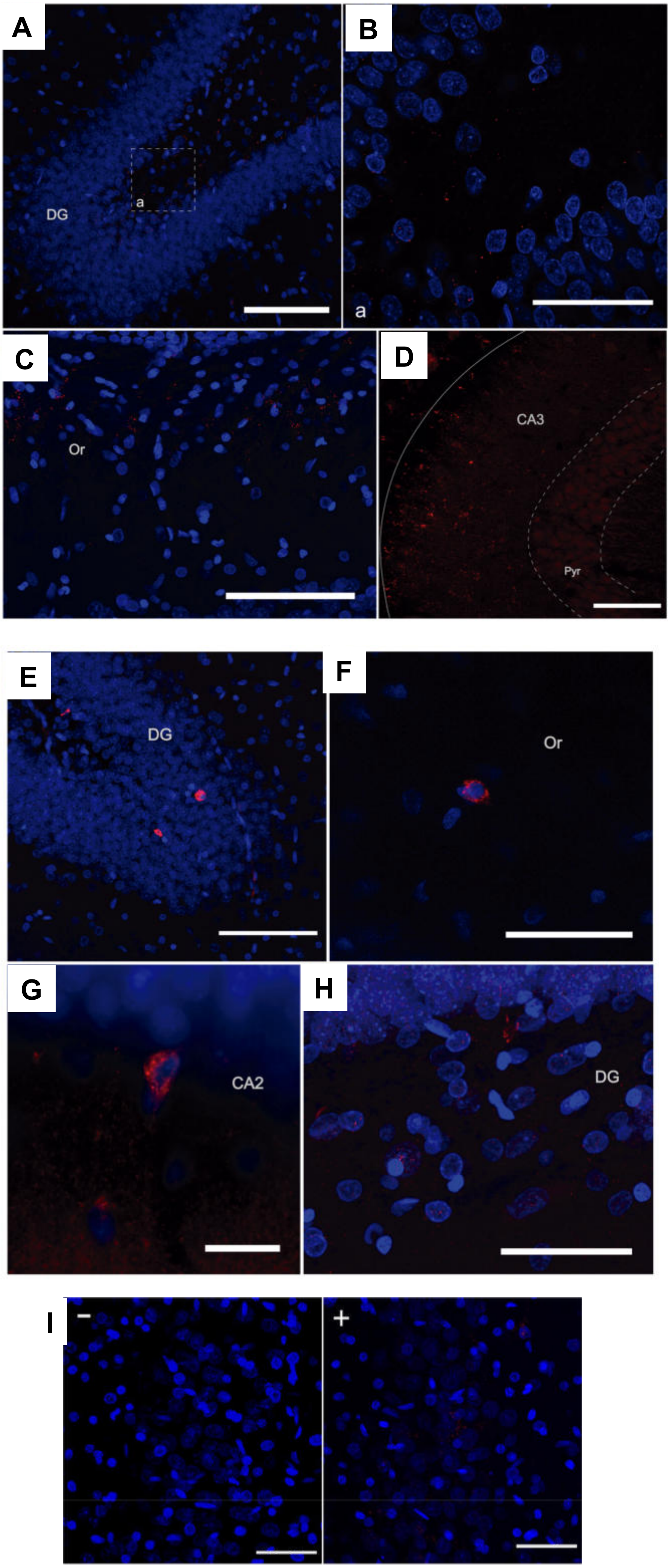

SUPP FIGURE 10

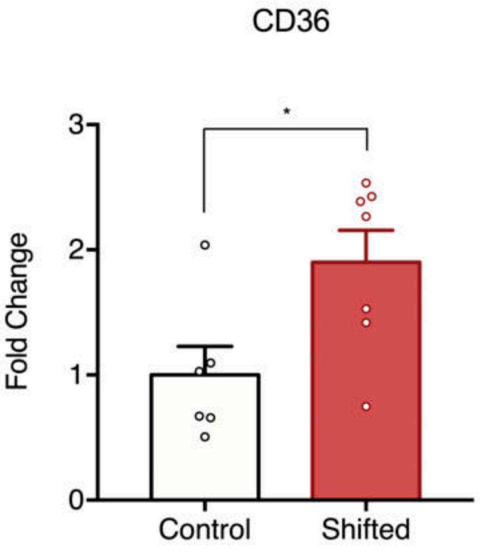


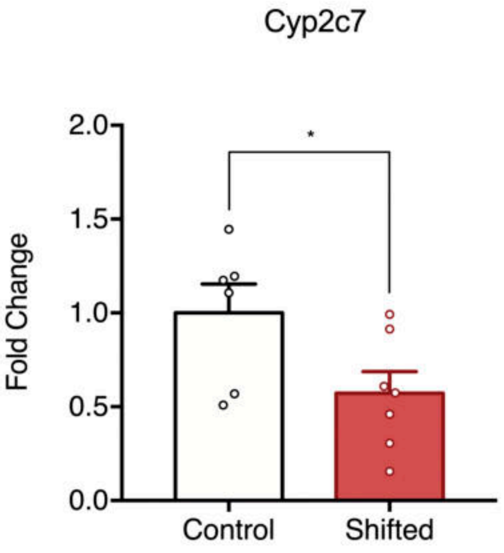

SUPP FIGURE 11

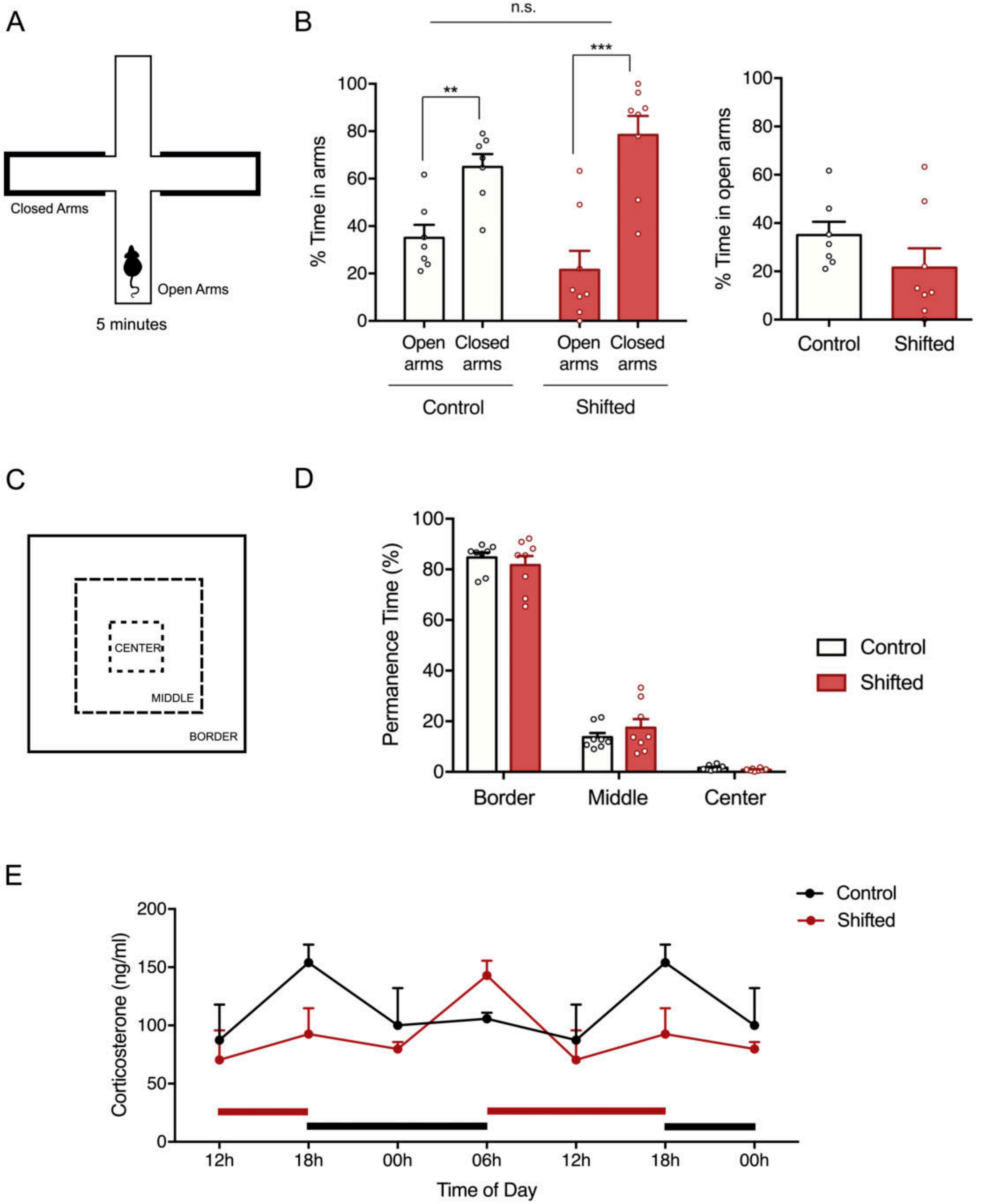

SUPP FIGURE 12

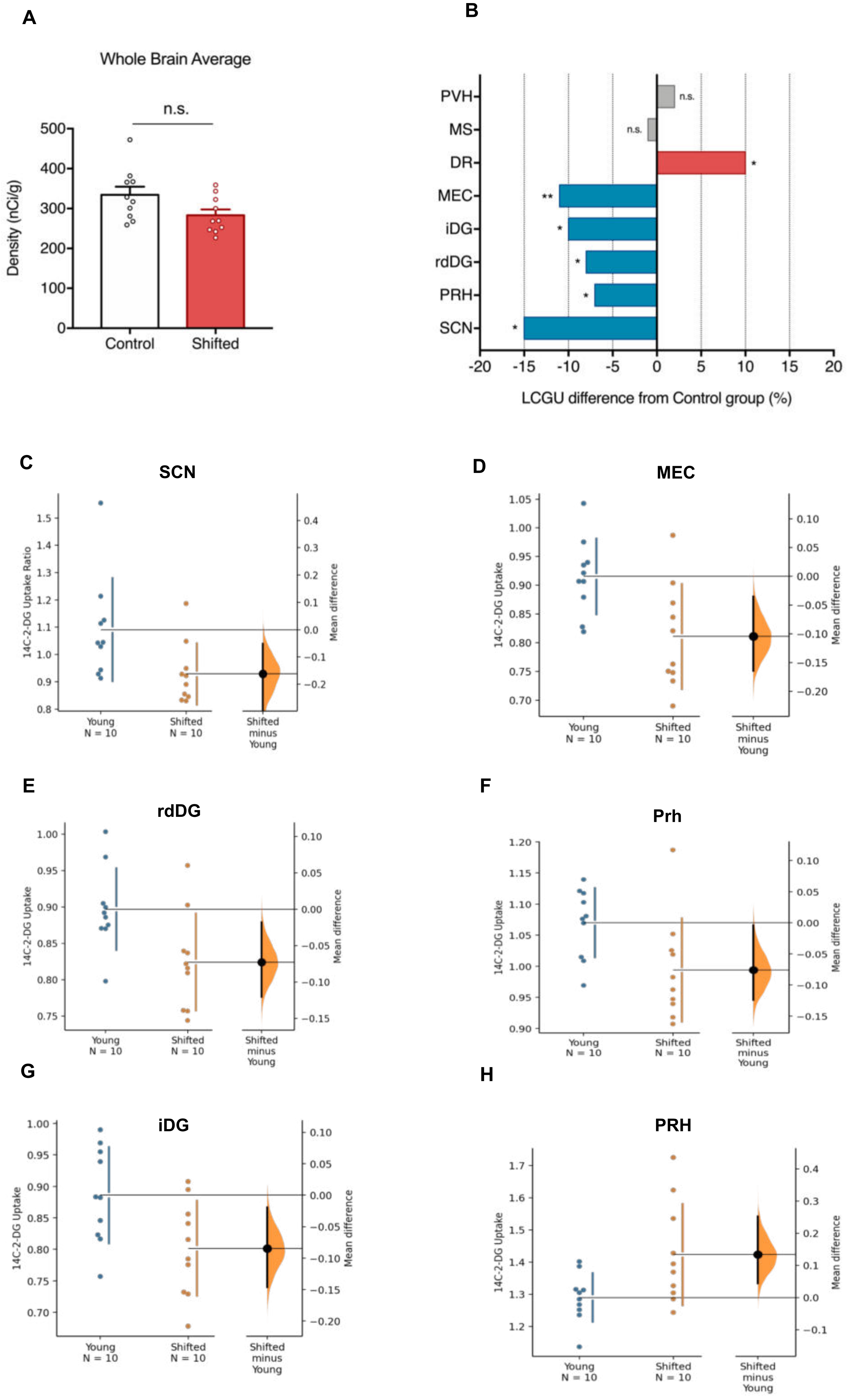

**SUPP FIGURE 13**

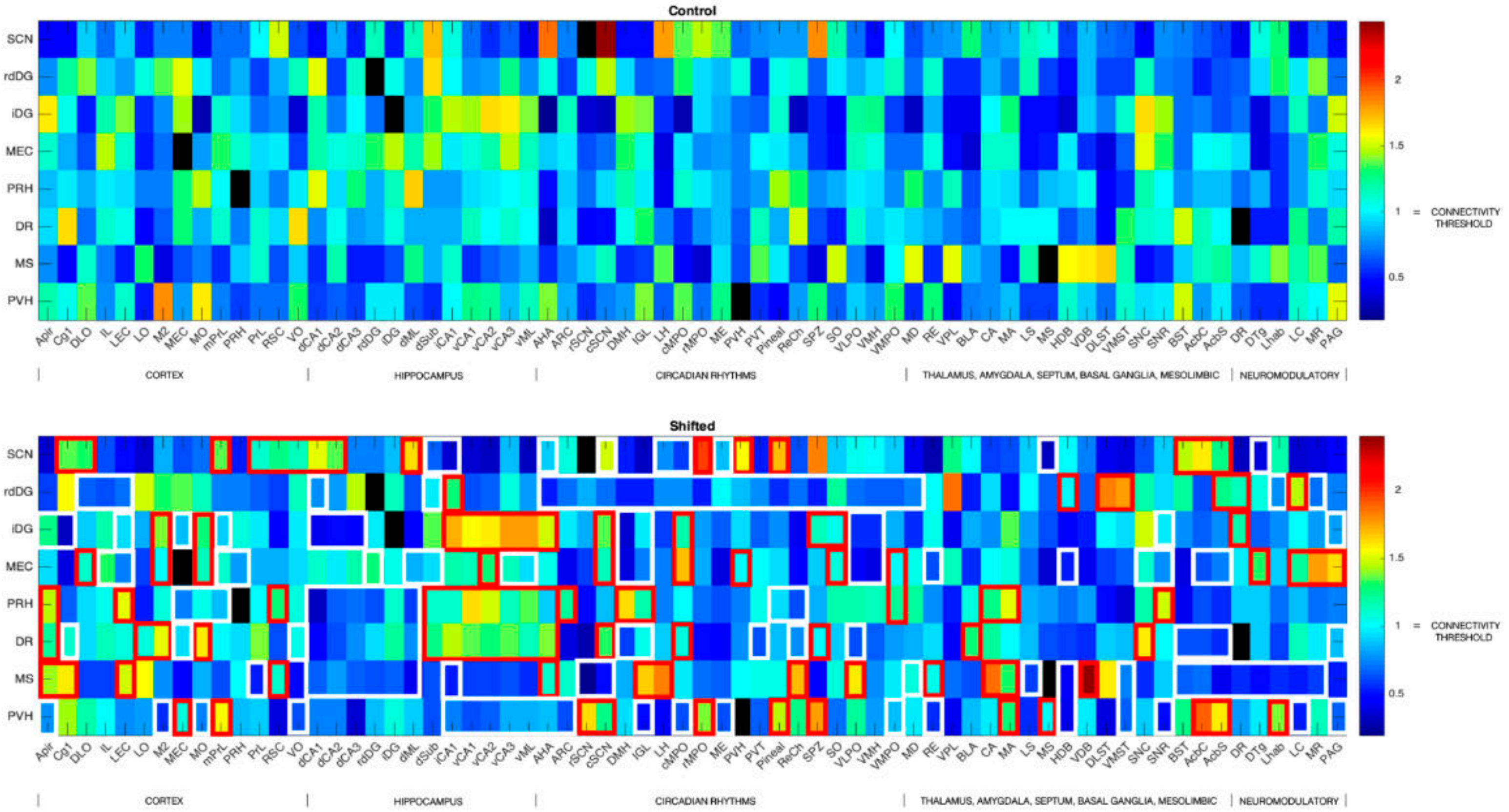

SCN
rdDG
iDG
MEC
PRH
DR
MS
PVH
Apir Cg1 DLO IL LEC LO M2 MEC MO mPrL PRH PrL RSC VO dCA1 dCA2 dCA3 rdDG iDG dML dSub iCA1 vCA1 vCA2 vCA3 vML AHA ARC rSCN cSCN DMH IGL LH cMPO rMPO ME PVH PVT Pineal ReCh SPZ SO VLPO VMH VMPO MD RE VPL BLA CA MA LS MS HDB VDB DLST VMST SNC SNR BST AcbC AcbS DR DTg Lhab LC MR PAG
CORTEX
HIPPOCAMPUS
CIRCADIAN RHYTHMS
THALAMUS, AMYGDALA, SEPTUM, BASAL GANGLIA, MESOLIMBIC
NEUROMODULATORY
2
1.5
1 = CONNECTIVITY THRESHOLD
0.5

SUPP FIGURE 14

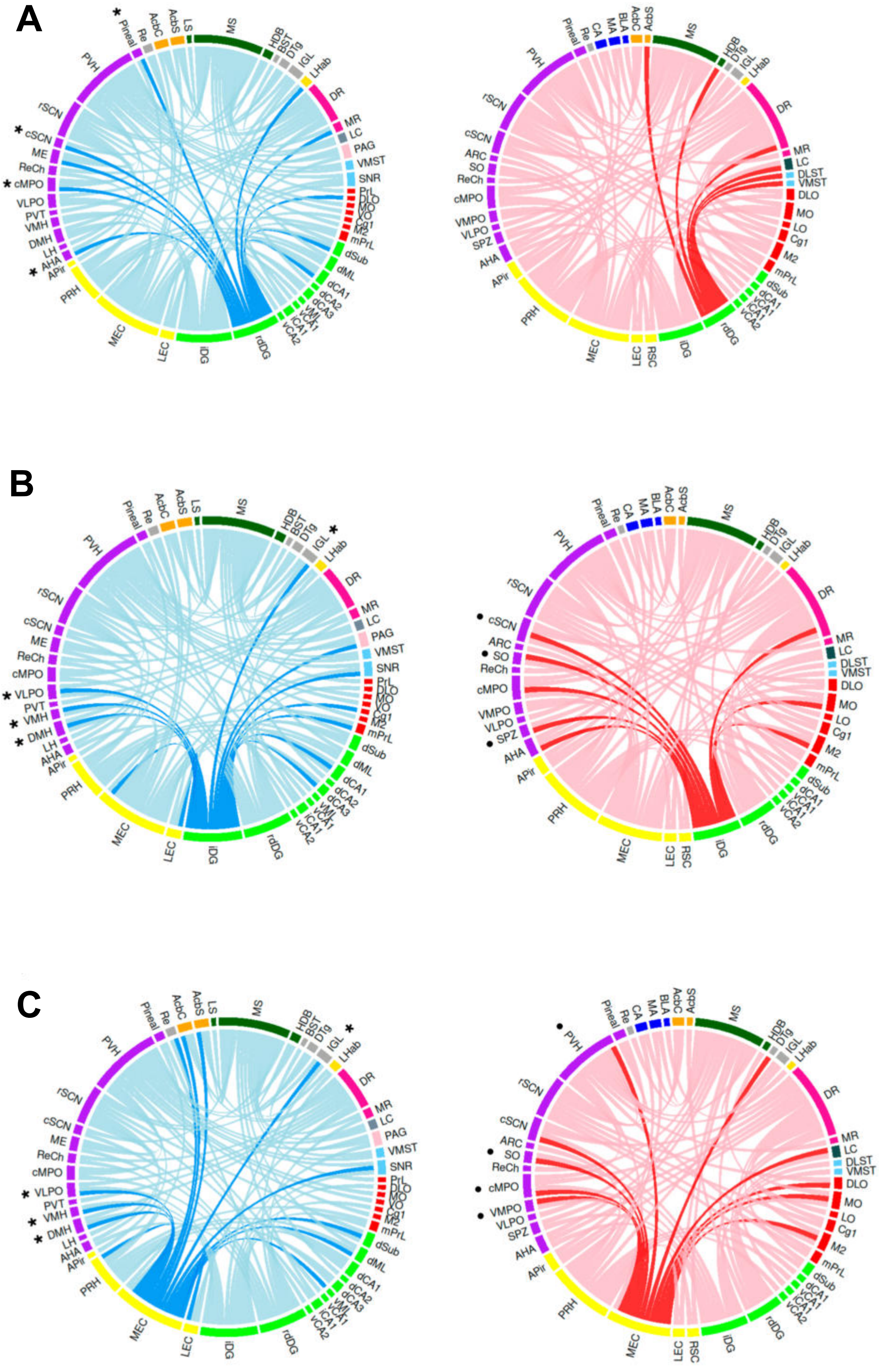

SUPP FIGURE 15

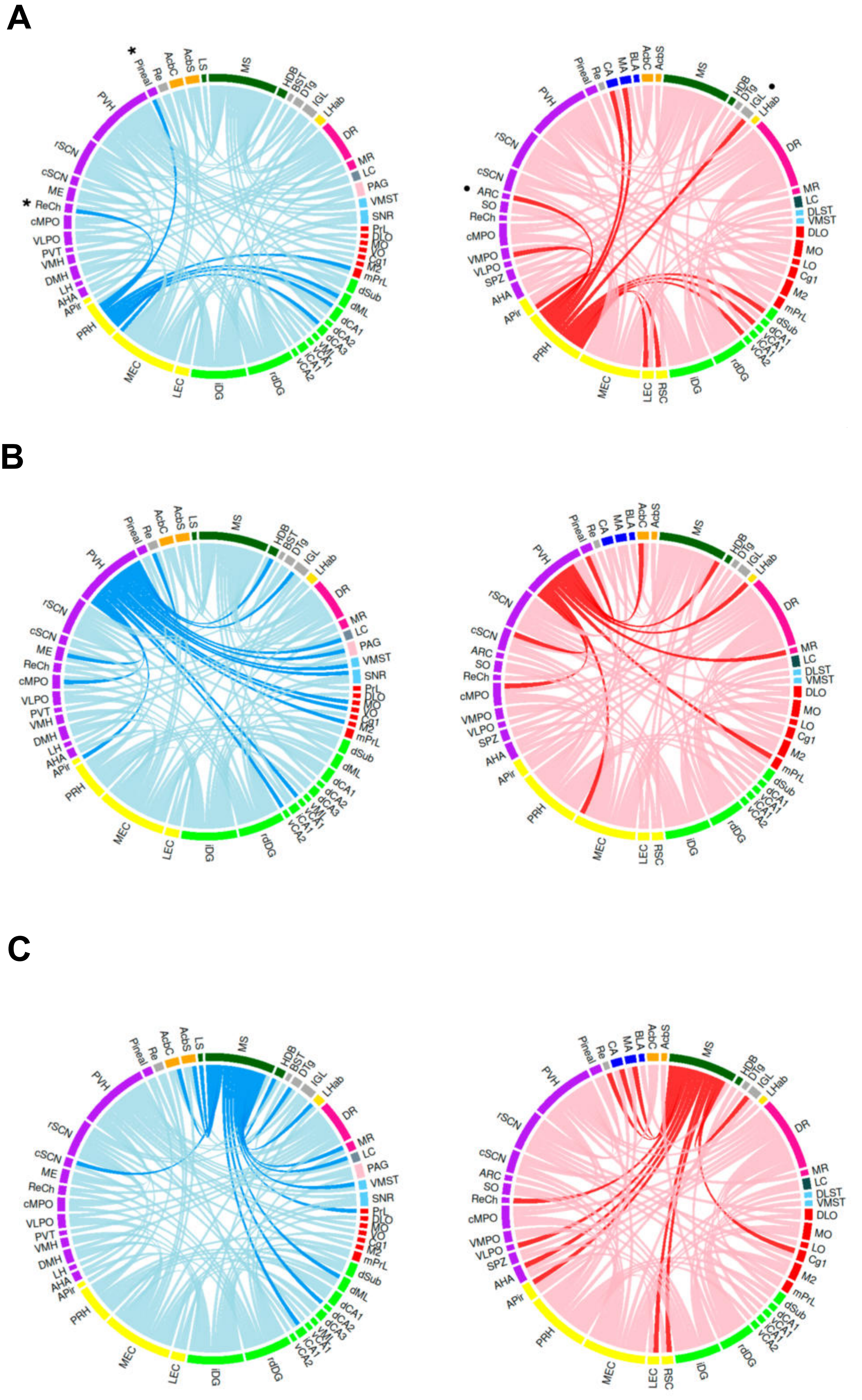

SUPP FIGURE 16

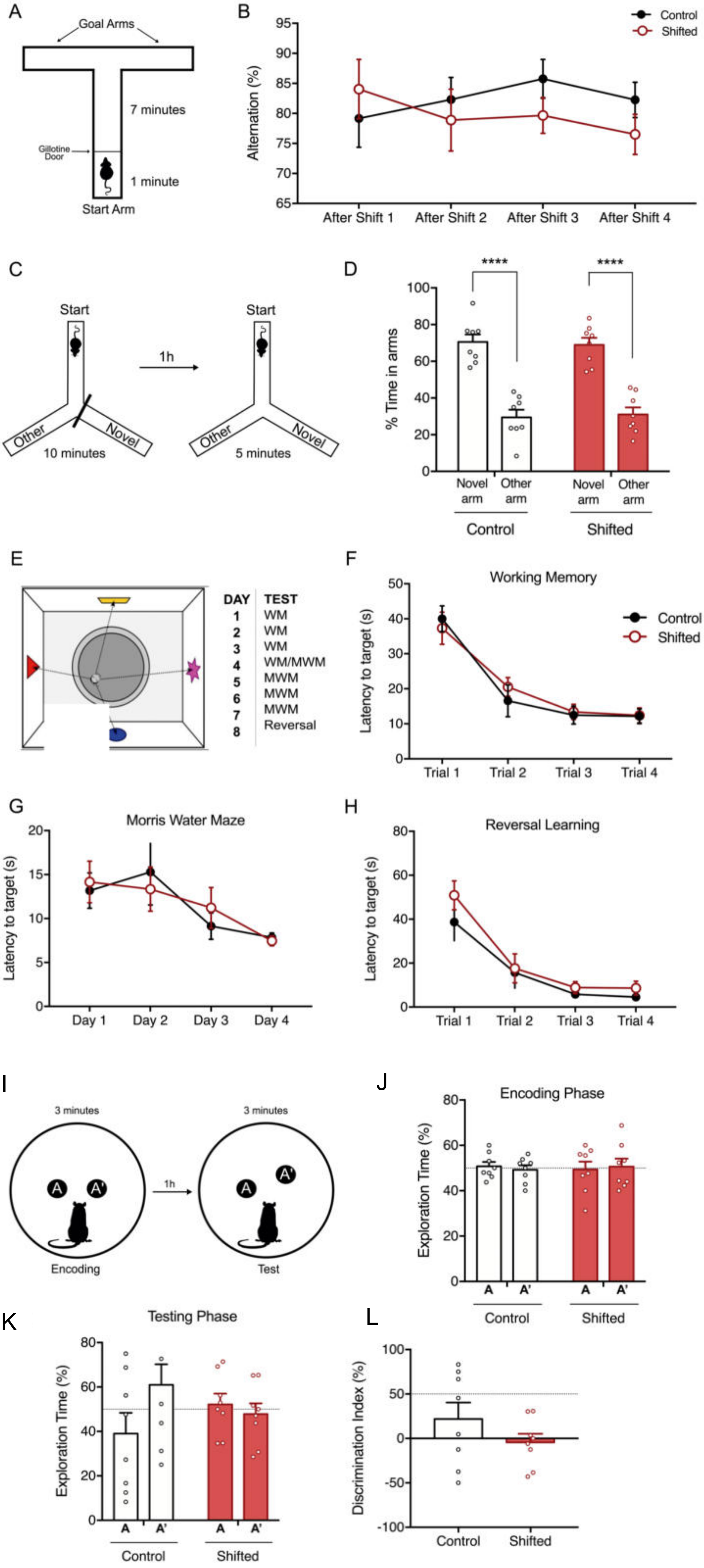

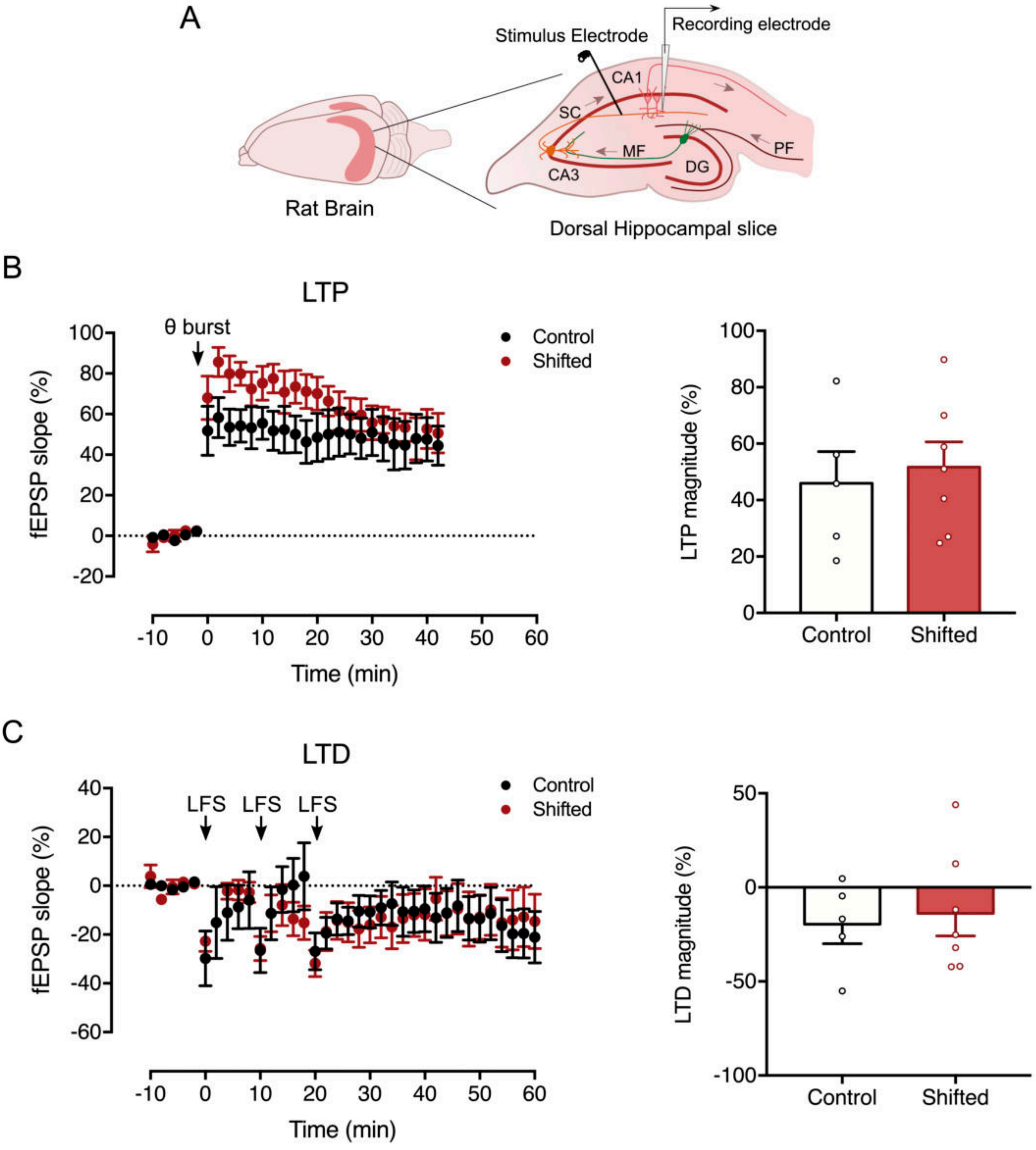
A
Stimulus Electrode
Recording electrode
CA1
SC
MF
PF
CA3
DG
Rat Brain
Dorsal Hippocampal slice
B
LTP
θ burst
Control
Shifted
fEPSP slope (%)
Time (min)
LTP magnitude (%)
Control
Shifted
C
LTD
LFS
LFS
LFS
Control
Shifted
fEPSP slope (%)
Time (min)
LTD magnitude (%)
Control
Shifted

**Supplementary Figure 1 - Substantial transsynaptic anterograde spread from the SCN is found in the dorsal hippocampus.**

The rVSV-(VSV-G)-Venus labeling (green) pattern of the hippocampus formation six days after injection into the SCN. (A) Progressively more posterior sections showing antero-posterior gradient of the spread of viral-labeled terminals from CA2 region to CA3 and Dentate Gyrus. (B) Representative image of the transsynaptic anterograde labeling of the hippocampus. Scale Bars: 500 µm. Abbreviations: *CA1 field of the hippocampus (CA1); CA2 field of the hippocampus (CA2); CA3 field of the hippocampus (CA3); Dentate Gyrus (DG); Lacunosum–Moleculare layer (LMol); Oriens layer (Or); Pyramidal cell layer (Pyr); Radiatum layer (Rad).*

**Supplementary Figure 2 - Substantial transsynaptic retrograde labeling of the SCN, septum and periventricular areas following viral injection into the dorsal hippocampus**

The pattern of spread of rVSV-(RABV-G)-eGFP labeling (green) five days after viral injection into the dorsal hippocampus. (A) Schematic representation of the viral injection into the dorsal hippocampal CA1 region. (B) Representative image of the injection spot. Scale bar: 500 µm. (C) Strong transsynaptic retrograde labeling projection of dorsal hippocampal CA1 to SCN and SPZ. Scale bar: 300 µm. (D) eGFP-labeled neurons in Lateral Septum (LSD, LSV) and Medial Septum (MS). Scale bar: 1000 µm. (E) Strong rVSV-(RABV-G)-eGFP expression in PVN and adjacent periventricular areas (Pe). Scale bar: 500 µm.

**Supplementary Figure 3 - Immunohistochemical analysis of Venus-labeled neurons and terminals in *stratum oriens* of dorsal hippocampal CA1 region.**

(A) Z-stack maximum intensity projection image taken at 20x magnification in the CA1 *stratum oriens*. ChAT positive terminals are labeled red. Scale Bar: 100 µm. (a) Representative high magnification (63x) image showing ChAT+ terminals (arrow) in apposition to Venus-labeled (green) neurons. Scale Bar: 25 µm. (b) Representative high magnification image showing a Venus-labeled axon ensheathed by ChAT+ terminals (arrow). Scale Bar: 25 µm.

(B) Z-stack maximum intensity projection image taken at 20x magnification in the CA1 *stratum oriens*. GAD 65/67 positive terminals are labeled red. Scale Bar: 100 µm. (a) Representative high magnification image showing co-localization of GAD 65/67+ terminals with a Venus-labeled neuronal cell body (arrow). Scale Bar: 25 µm. (b) Representative high magnification image showing the close appositions of GAD 65/67+ terminals with Venus-labeled terminals. Scale Bar: 25 µm.

**Supplementary Figure 4 - Immunohistochemical analysis of Venus-labeled neurons and terminals in dorsal hippocampal CA2 region.**

(A) Z-stack maximum intensity projection image taken at 20x magnification in the CA2 *stratum pyramidale*. ChAT positive terminals are labeled red. Scale Bar: 100 µm. (a) Representative high magnification (63x) image showing ChAT+ terminals in apposition to Venus-labeled (green) neurons. Scale Bar: 25 µm. (b) Representative high magnification image showing a Venus-labeled axon and dendrites wrapped by ChAT+ terminals (arrows). Scale Bar: 25 µm.

(B) Z-stack maximum intensity projection image taken at 20x magnification in CA2 *strata pyramidale and oriens*. ChAT positive terminals are labeled red. Scale Bar: 100 µm. (a) Representative high magnification image of Venus-labeled terminals with ChAT+ terminals. Scale Bar: 25 µm.

(C) Z-stack maximum intensity projection image taken at 20x magnification in the CA2 *stratum pyramidale*. GAD 65/67 positive terminals are labeled red. Scale Bar: 100 µm. (a) Representative high magnification image of GAD 65/67+ with Venus-labeled neuronal cell body. Scale Bar: 25 µm.

(D) Z-stack maximum intensity projection image taken at 20x magnification in the CA2 *stratum pyramidale*. CamKII positive cells are labeled gray. Scale Bar: 100 µm. (a) Representative high magnification image co-localization of CamKII with Venus-labeled neuronal cell bodies. Scale Bar: 25 µm.

**Supplementary Figure 5 - Immunohistochemical analysis of Venus-labeled neurons and terminals in dorsal hippocampal CA3 region.**

(A) Z-stack maximum intensity projection image taken at 20x magnification in the CA3 *stratum pyramidale*. ChAT positive terminals are labeled red. Scale Bar: 100 µm. (a) Representative high magnification (63x) image showing ChAT+ terminals in apposition to Venus-labeled (green) neurons. Scale Bar: 25 µm. (b) Representative high magnification image showing Venus-labeled axons sheathed by ChAT+ terminals (arrows). Scale Bar: 25 µm.

(B) Z-stack maximum intensity projection image taken at 20x magnification in the CA3 *stratum lucidum*. ChAT positive terminals are labeled red. Scale Bar: 100 µm. (a) Representative high magnification image showing co-localization and close proximity of ChAT+ terminals and Venus-labeled terminals. Scale Bar: 25 µm.

(C) Z-stack maximum intensity projection image taken at 20x magnification in the CA3 *strata pyramidale* and *lucidum*. GAD 65/67 positive terminals are labeled red. Scale Bar: 100 µm. (a) Representative high magnification image showing co-localization of GAD 65/67+ staining with Venus-labeled neuronal cell body. Scale Bar: 25 µm.

(D) Z-stack maximum intensity projection image taken at 20x magnification in the CA3 *stratum pyramidale*. CamKII positive cells are labeled gray. Scale Bar: 100 µm. (a) Representative high magnification image showing co-localization of CamKII with Venus-labeled neuronal cell bodies. Scale Bar: 25 µm.

**Supplementary Figure 6 - Immunohistochemical analysis of Venus-labeled neurons in the granular layer of the dorsal Dentate Gyrus (dDG)**

(A) Z-stack maximum intensity projection image taken at 20x magnification in the granular layer and subgranular zone of the dorsal Dentate Gyrus (dDG). ChAT positive terminals are labeled red. Scale Bar: 100 µm. (a) Representative high magnification (63x) image showing co-localization and close apposition of ChAT+ terminals with Venus-labeled (green) neurons. Scale Bar: 25 µm.

(B) Z-stack maximum intensity projection image taken at 20x magnification in the dDG granular layer. GAD 65/67 positive terminals are labeled red. Scale Bar: 100 µm. (a) Representative high magnification image showing co-localization of GAD 65/67 with Venus-labeled neurons. Scale Bar: 25 µm.

(C) Z-stack maximum intensity projection image taken at 20x magnification in the dDG granular layer and subgranular zone. CamKII positive cells are labeled gray. Scale Bar: 100 µm. (a) Representative high magnification image of CamKII with Venus-labeled neuronal cell bodies. Scale Bar: 25 µm. All images were taken in the coronal plane.

**Supplementary Figure 7 - Immunohistochemical analysis of Venus-labeled neurons in the hilus and granular layer of the dorsal Dentate Gyrus (dDG)**

(A) Z-stack maximum intensity projection image taken at 20x magnification in the dDG hilus. ChAT positive terminals are labeled red. Scale Bar: 100 µm. (a) Representative high magnification (63x) image showing apposition of ChAT+ terminals to Venus-labeled (green) neurons. Scale Bar: 25 µm. (b) Representative high magnification image showing co-localization and close proximity of ChAT+ terminals and Venus-labeled terminals. Scale Bar: 25 µm.

(B) Z-stack maximum intensity projection image taken at 20x magnification in the dDG granular layer and hilus. GAD 65/67 positive terminals are labeled red. Scale Bar: 100 µm. (a) Representative high magnification image showing co-localization of GAD 65/67 with Venus-labeled neurons in the subgranular zone. Scale Bar: 25 µm. (b, c) Representative high magnification image showing co-localization of GAD 65/67 with Venus-labeled neurons in the hilus. Scale Bars: 25 µm.

(C) Z-stack maximum intensity projection image taken at 20x magnification in granular layer and hilus. CamKII positive cells are labeled gray. Scale Bar: 100 µm. (a) Representative high magnification of a confocal plane showing co-localization with Venus-labeled neuronal cell body in the subgranular zone. Scale Bar: 25 µm. (b, c) Representative high magnification image showing co-localization of CamKII with Venus-labeled neurons in the hilus. Scale Bars: 25 µm.

**Supplementary Figure 8 – BDA-Texas Red labeling of regional targets after injection into the SCN**

Terminal labeling of first-order efferents eleven days after injection of BDA-Texas Red into the SCN. (A) Schematic representation of the tracer injection into the SCN. (B) Representative fluorescence microscope image of the injection site. The core region is delineated by immunofluorescent labeling of NPY+ terminals (gray). Unilateral injection into the SCN resulted in intense fluorescent labeling of cell bodies across both the core and shell subregions. Labeling was also observed in the contralateral core. Scale Bar: 250 µm. (C) Terminal labeling in the Bed Nucleus of the Stria Terminalis (BNST). Scale Bar: 100 µm. (D) Terminal labeling in the Paraventricular Hypothalamic Nucleus (PVN). Cell nuclei are stained with DAPI in blue. Scale Bar: 100 µm. (E) Terminal labeling in the Lateral Septum (LS). Scale Bar: 50 µm. (F) Terminal labeling in the Medial Septum (MS). Scale Bar: 50 µm.

**Supplementary Figure 9 – BDA-Texas Red labeling of terminals across the Hippocampus after tracer injection in the SCN is consistent with the results from anterograde transsynaptic tracing**

Terminal labeling was found in the hippocampus eleven days after injection of BDA-Texas Red into the SCN. (A) Terminal labeling in the hilus of the dorsal Dentate Gyrus (DG). Scale Bar: 100 µm. (a) Representative high magnification (63x) image showing the labeled terminals in the hilus of the dorsal Dentate Gyrus. Scale Bar: 50 µm. (C) Terminal labeling in the *stratum oriens* (Or). Scale Bar: 100 µm. (D) Terminal labeling in the dorsal CA3 region. Scale Bar: 100 µm. (E) Neurons labeled with clustered terminals in the granular layer and subgranular zone of the dorsal Dentate Gyrus (DG). Scale Bar: 100 µm. (F) Neuron labeled with clustered terminals in the *stratum oriens*. Scale Bar: 50 µm. (G) Neuron labeled with clustered terminals in the dorsal CA2 region. Scale Bar: 25 µm. (H) Magnification (63x) showing clustered terminals in the subgranular zone of the dorsal Dentate Gyrus. Scale Bar: 50 µm.

I) Negative control of the immunohistochemistry for BDA-terminal labeling amplification

(+) Terminal labeling in the CA3 region of the hippocampus amplified with immunohistochemistry using Texas Red Antibody (-) Terminal labeling in the CA3 region of the hippocampus is absent in negative control for the immunohistochemistry. Scale Bars: 50 µm.

**Supplementary Figure 10– Chronic Manipulation of Circadian Rhythms significantly altered biomarkers of circadian rhythm disturbance**

*CD36* and *Cyp2c7* mRNA expression were evaluated in liver samples of control and CCD/shifted animals. Results are presented as the mean ± SEM. (n=6-7, * $p<0.05$, unpaired Student's t-test)

**Supplementary Figure 11– Chronic Manipulation of Circadian Rhythms does not impact on anxiety-like behaviors and corticosterone daily secretion**

Anxiety-like behaviors were assessed using Elevated Plus Maze (EPM) and Open Field Test (OFT). Corticosterone levels were measured across seven timepoints. (A) Schematic representation of the EPM. (B) No significative differences were observed in the time spent in the open or closed arms of the EPM between shifted and control animals. Both shifted and control animals spent significantly more time in the closed arm than the open arm (n=8, p>0.05, unpaired Student's t-test. ** p<0.01, *** p<0.001 unpaired Student's t-test). (C) Schematic representation of the OFT. (D) Permanence in the open-field's subregions was similar between shifted and control animals (n=8, p>0.05, unpaired Student's t-test). (E) Corticosterone levels were rhythmic in shifted animals and were aligned with light/dark cycle. No differences in overall levels of corticosterone were observed between shifted and control animals (n=2-3). Data is double-plotted for clarity on rhythm periodicity. Red bars at the bottom indicate the dark phase of the LD cycle of shifted animals. Black bars indicate the dark phase of the LD cycle of control animals. All values are mean ± SEM.

**Supplementary Figure 12– Differences in Local Cerebral Glucose Utilization (LCGU) between shifted/CCD and control animals in the seed regions used in PLSR model.**

A) The whole brain average $^{14}C$-2-deoxyglucose concentration (nCi/g) was determined as the average $^{14}C$ concentration across all sections in which a RoI was measured. No significant differences were observed between control and shifted animals (n=10; p>0.05, unpaired Student's t-test). Values are mean ± SEM.

B) Quantification of the Chronic Manipulation of Circadian Rhythms-induced metabolic alterations in the seed regions selected for modeling functional regional connectivity. Results are presented as percentage difference between average LCGU between shifted and control animals. (n=10, * p<0.05, ** p<0.01, unpaired)

C-H) The mean difference between Young and CCD/Shifted for the indicated seed region is shown in the above Gardner-Altman estimation plot. Both groups are plotted on the left axes; the mean difference is plotted on a floating axes on the right as a bootstrap sampling distribution. The mean difference is depicted as a dot; the 95% confidence interval is indicated by the ends of the vertical error bar. 5000 bootstrap samples were taken; the confidence interval is bias-corrected and accelerated.Any *P* value reported is the probability of observing the effect size (or greater),assuming the null hypothesis of zero difference is true. For each permutation *P* value, 5000 reshuffles of the control and test labels were performed.

**Supplementary Figure 13– Visual representation of the functional coupling between brain regions**

8x68 matrices heatmaps displaying unfiltered values of VIP statistic, for visual inspection of patterns of functional coupling changes between control and shifted groups. Salient differences between regions are highlighted. Regional connectivity increase in shifted animals is highlighted in red, regional connectivity decrease is highlighted in white. **Cortex:** Amygdalo-Piriform Cortex (APir), Cingulate (Cg), Dorsolateral Orbital (DLO),

Infralimbic (IL), Lateral Entorhinal (LEC), Lateral Orbital (LO), Supplementary Motor Area (M2), Medial Entorhinal (MEC), Medial Orbital (MO), Medial Prefrontal (mPFC), Prelimbic (PrL), Retrosplenial (RSC), Ventral Orbital (VO). **Hippocampus:** Dorsal CA1 (dCA1), Dorsal CA2 (dCA2), Dorsal CA3 (dCA3), Dorsal Dentate Gyrus (dDG), Intermediate Dentate Gyrus (iDG), Dorsal Hippocampus Molecular Layer (dML), Dorsal Subiculum (dSub), Intermediate CA1 (iCA1), Intermediate CA3 (iCA3), Ventral CA1 (vCA1), Ventral CA2 (vCA2), Ventral CA3 (vCA3), Ventral Dentate Gyrus (vDG), Ventral Hippocampus Molecular Layer (vML). **Circadian Rhythms**: Anterior Hypothalamus (AHA), Arcuate Nucleus (ARC), Suprachiasmatic Nucleus – Rostral (rSCN), Suprachiasmatic Nucleus – Caudal (cSCN), Dorsomedial Hypothalamus (DMH), Intergeniculate Leaflet (IGL), Lateral Hypothalamus (LH), Medial Preoptic Nucleus – Caudal (cMPO), Medial Preoptic Nucleus – Rostral (rMPO), Median Eminence (ME), Paraventricular Hypothalamic Nucleus – Parvicellular (PVp), Paraventricular Thalamic Nucleus (PVT), Preoptic Area (POA), Retrochiasmatic Area (RCH), Supraoptic Nucleus (SON), Suprachiasmatic Nucleus (SCN), Ventromedial Hypothalamus (VMH), Ventral Thalamic Nucleus (VL). **Thalamus**: Mediodorsal (MD), Reuniens (RE). **Amygdala**: Basolateral (BLA), Central (CeA), Medial (MeA). **Septum**: Lateral Septum (LS), Medial Septum (MS). Diagonal Band of Broca: Horizontal Band (HDB), Vertical Band (VDB). **Basal Ganglia**: Dorsolateral Striatum (DLS), Ventromedial Striatum (VMS), Substantia Nigra pars compacta (SNc), Substantia Nigra pars reticulata (SNr). **Mesolimbic**: Bed Nucleus of the Stria Terminalis (BST), Nucleus Accumbens core (AcbC), Nucleus Accumbens shell (AcbSh). **Neuromodulatory**: Dorsal Raphe (DR), Dorsal Tegmental Nucleus (DTg), Lateral Habenula (LHb), Locus Coeruleus (LC), Median Raphe (MR), Periaqueductal Gray (PAG).

**Supplementary Figure 14– Chronic Manipulation of Circadian Rhythm induces alterations in rostrodorsal, intermediate Dentate Gyrus (rdDG and iDG) and Medial Entorhinal Cortex (MEC) Functional Connectivity (FC)**

Chord diagrams summarizing significant alterations in the FC of the A) rdDG; B) iDG and C) MEC in CCD animals.

(Left) Significant losses (dark blue) of FC in CCD animals. *denotes functional connections with SCN and its efferents that were anatomically confirmed and were lost upon chronic circadian manipulation.

(Right) Significant gains (dark red) in FC in CCD animals.

The significance of FC alterations was analyzed using unpaired Student's t-test with Bonferroni *post hoc* correction. Significance was set a $p<0.05$. The lighter colors represent all significant lost/gained connections across all seed regions examined. Full data are shown in Supplementary Tables 3-5

**Supplementary Figure 15– Chronic Manipulation of Circadian Rhythm induces alterations in Perirhinal Cortex (PRH), Paraventricular Hypothalamic Nucleus (PVH) and C) Medial Septum (MS) Functional Connectivity (FC)**

Chord diagrams summarizing significant alterations in the FC of the A) Perirhinal Cortex (PRH); B) Paraventricular Hypothalamic Nucleus (PVH) and C) Medial Septum (MS).

Significant losses (dark blue) of FC in CCD animals. *denotes functional connections with SCN and its efferents that were anatomically confirmed and were lost upon chronic circadian manipulation. (Right) Significant gains (dark red) in FC in CCD animals.

The significance of FC alterations was analyzed using unpaired Student's t-test with Bonferroni post hoc correction. Significance was set a $p<0.05$. Faded colors represent total significant lost/gained connections across all seed regions. Strong colors depict PRH's significant lost/gained connections. Full data are shown in Supplementary Tables 6-8.

**Supplementary Figure 16 – Chronic Manipulation of Circadian Rhythms does not affect Short-Term Memory, Long-Term Memory nor Reversal Learning**.

(B)Schematic representation of the T-Maze test, animals spend one minute in the start arm and then freely explore the maze for 7 minutes. (B) Spontaneous alternation, a measure of spatial working memory, was assessed using T-Maze test. No significant differences were found between the performance of shifted and control group across the duration of the Chronic Manipulation of Circadian Rhythms protocol. (C) Schematic representation of the Y-Maze test of short-term memory, assessed through presentation of the novel arm 1 hour after previous maze exposure. (D) No significant differences were found between the performance of shifted and control group across the duration of the Chronic Manipulation of Circadian Rhythms protocol in the Y-Maze short-term memory test. Both groups displayed a preference for the novel arm (n=8, **** $p<0.0001$, novel arm comparing to other arm, two-way ANOVA) (E) Schematic representation of the Morris Water Maze test. Working Memory (WM), Long-Term Memory (MWM) and Reversal Learning were sequentially tested across different days. (F) No significant differences were found in Working Memory between shifted and control groups. (G) No significant differences were found in Long-Term learning and memory between shifted and control groups. (H) No significant differences were found in Reversal Learning between shifted and control groups. (I) Schematic representation of Pattern Separation Task, showing the encoding and test phase with a 1-hour inter-trial interval. (J) Performance during encoding phase was not significantly different between the experimental groups. (K) Although not significantly different, there is a trend for altered performance in the shifted animals. (L) Although Discrimination Index is not statistically significant, there is a trend for altered performance in the shifted animals. All values are mean ± SEM.

**Supplementary Figure 17 – Chronic Manipulation of Circadian Rhythms does not impact hippocampal synaptic plasticity in Shaffer Collatoral-CA1 synapses**

(A) Schematic representation of the simplified circuitry of the hippocampus. A stimulation electrode was placed to stimulate the Schaffer collaterals and the recording electrode in the *stratum radiatum* of the CA1 area. (B, left) Changes in the field excitatory post-synaptic potential (fEPSP) slope following LTP induction by theta-burst stimulation (ten trains with four pulses each at 100 Hz, separated by 200 ms) recorded from control and CCD/shifted animal hippocampal slices (n = 5 and 7, respectively). (B, right) LTP magnitude after theta-burst stimulation. (C, left) Changes in the fEPSP slope following LTD induction by Low Frequency Stimulation (LFS) (three trains with 10-min interval of 2 Hz, 1,200 pulses) (C, right) LTD magnitude after Low Frequency Stimulation (change in fEPSP slope at 50–60 min)

| Region | Control Mean | | Control SEM | Shifted Mean | | Shifted SEM | p-value | Lost/Gained |
|---|---|---|---|---|---|---|---|---|
| *Cortex* | | | | | | | | |
| Amigdalo-Piriform Cortex (APir) | 0,46 | ± | 0,03 | 0,37 | ± | 0,05 | 52,66998 | |
| Cingulate (Cg1) | 0,45 | ± | 0,05 | **1,41*** | ± | **0,09** | 3,2E-06* | Gained |
| Dorsolateral Orbital (DLO) | 0,91 | ± | 0,03 | **1,35*** | ± | **0,06** | 0,0016* | Gained |
| Infralimbic (IL) | 0,72 | ± | 0,05 | 0,67 | ± | 0,10 | 298,4077 | |
| Lateral Entorhinal (LEC) | 0,90 | ± | 0,04 | 0,51 | ± | 0,06 | 0,021073 | |
| Lateral Orbital (LO) | 0,54 | ± | 0,04 | 0,36 | ± | 0,07 | 21,95772 | |
| Supplementary Motor Area (M2) | 0,73 | ± | 0,05 | 0,82 | ± | 0,06 | 124,8604 | |
| Medial Entorhinal (MEC) | 0,78 | ± | 0,06 | 0,67 | ± | 0,07 | 124,2155 | |
| Medial Orbital (MO) | 0,43 | ± | 0,04 | 0,71 | ± | 0,05 | 0,185812 | |
| Medial Prelimbic (mPrL) | 0,70 | ± | 0,05 | **1,40*** | ± | **0,06** | 9,2E-06* | Gained |
| Perirhinal (PRH) | 0,70 | ± | 0,03 | 0,52 | ± | 0,04 | 1,46572 | |
| Prelimbic (PrL) | **1,04** | ± | **0,05** | **1,15** | ± | **0,06** | 77,73604 | |
| Retrosplenial (RSC) | **1,52** | ± | **0,08** | **1,26** | ± | **0,07** | 9,88897 | |
| Ventral Orbital (VO) | 0,65 | ± | 0,09 | **1,18** | ± | **0,12** | 1,396442 | |
| *Hippocampus* | | | | | | | | |
| Dorsal CA1(dCA1) | 0,49 | ± | 0,05 | **1,56*** | ± | **0,06** | 3E-09* | Gained |
| Dorsal CA2 (dCA2) | 0,87 | ± | 0,05 | **1,39** | ± | **0,11** | 0,162294 | |
| Dorsal CA3 (dCA3) | 0,50 | ± | 0,03 | 0,73 | ± | 0,05 | 0,425081 | |
| Dorsal Dentate Gyrus - Rostral (rdDG) | **1,21** | ± | **0,09** | 0,76 | ± | 0,09 | 1,135093 | |
| Intermediate Dentate Gyrus (iDG) | 0,52 | ± | 0,04 | 0,89* | ± | 0,05 | 0,0050* | |
| Dorsal Hippocampus Molecular Layer (dML) | **1,17** | ± | **0,04** | **1,64*** | ± | **0,06** | 0,0007* | Increased |
| Dorsal Subiculum (dSub) | **1,77** | ± | **0,12** | 0,67* | ± | 0,04 | 1,6E-05* | Lost |
| Intermediate CA1 (iCA1) | **1,10** | ± | **0,06** | 0,38* | ± | 0,05 | 4E-06* | Lost |
| Ventral CA1 (vCA1) | 0,69 | ± | 0,03 | 0,36* | ± | 0,05 | 0,0110* | |
| Ventral CA2 (vCA2) | 0,55 | ± | 0,03 | 0,32 | ± | 0,05 | 0,221151 | |
| Ventral CA3 (vCA3) | 0,68 | ± | 0,04 | 0,87 | ± | 0,05 | 2,459915 | |
| Ventral Hippocampus Molecular Layer (vML) | 0,42 | ± | 0,04 | 0,32 | ± | 0,05 | 54,57965 | |
| *Circadian Rhythms* | | | | | | | | |
| Anterior Hypothalamus AHA | **1,96** | ± | **0,15** | 0,93* | ± | 0,08 | 0,0035* | Lost |
| Arcuate Nucleus (ARC) | 0,76 | ± | 0,06 | **1,09** | ± | **0,08** | 1,374363 | |
| Caudal SCN (cSCN) | **2,48** | ± | **0,11** | **1,52*** | ± | **0,10** | 0,0013* | Decreased |
| Dorsomedial Hypothalamus (DMH) | 0,51 | ± | 0,09 | 0,61 | ± | 0,13 | 248,1794 | |
| Intergeniculate Leaftlet (IGL) | 0,49 | ± | 0,06 | 0,34 | ± | 0,06 | 35,8419 | |
| Lateral Hypothalamus (LH) | **1,84** | ± | **0,04** | 0,81* | ± | 0,11 | 1,1E-05* | Lost |
| Medial Preoptic Nucleus - Caudal (cMPO) | **1,41** | ± | **0,09** | 0,63* | ± | 0,06 | 0,0003* | Lost |
| Medial Preoptic Nucleus - Rostral (rMPO) | **1,50** | ± | **0,06** | **1,98*** | ± | **0,03** | 0,0001* | Increased |
| Median Eminence (ME) | **1,39** | ± | **0,04** | 0,88* | ± | 0,05 | 3,6E-05* | Lost |
| Paraventricular Hypothalamic Anterior - Parvicellular (PVH) | 0,70 | ± | 0,04 | **1,60*** | ± | **0,08** | 2,2E-06* | Gained |
| Paraventricular Thalamic Nucleus (PVT) | 0,80 | ± | 0,08 | 0,53 | ± | 0,04 | 3,792214 | |
| Pineal Gland | 0,84 | ± | 0,06 | **1,72*** | ± | **0,03** | 4,3E-08* | Gained |
| Retrochiasmatic Area (ReCh) | 0,81 | ± | 0,05 | 0,69 | ± | 0,14 | 208,4929 | |
| Subparaventricular Zone (SPZ) | **1,89** | ± | **0,10** | **1,84** | ± | **0,03** | 304,9976 | |
| Supraoptic Nucleus (SO) | **1,20** | ± | **0,03** | 0,93 | ± | 0,10 | 8,027177 | |
| Ventrolateral Preoptic Nucleus (VLPO) | 0,71 | ± | 0,08 | **1,05** | ± | **0,04** | 0,638068 | |
| Ventromedial Hypothalamus (VMH) | 0,49 | ± | 0,06 | **1,06** | ± | **0,11** | 0,148051 | |
| Ventromedial Preoptic Nucleus (VMPO) | **1,03** | ± | **0,04** | 0,93 | ± | **0,06** | 87,10795 | |
| *Thalamus* | | | | | | | | |
| Mediodorsal (MD) | 0,62 | ± | 0,05 | 0,45 | ± | 0,08 | 46,51276 | |
| Reuniens (RE) | 0,82 | ± | 0,09 | 0,33 | ± | 0,11 | 1,430299 | |
| Ventral Posterolateral (VPL) | 0,80 | ± | 0,04 | **1,27*** | ± | **0,05** | 0,0002* | Gained |
| *Amygdala* | | | | | | | | |
| Basolateral (BLA) | **1,34** | ± | **0,08** | **1,02** | ± | **0,06** | 1,92554 | |
| Central (CA) | 0,84 | ± | 0,07 | 0,59 | ± | 0,04 | 3,53313 | |
| Medial (MA) | 0,86 | ± | 0,05 | 0,62 | ± | 0,03 | 0,546581 | |
| *Septum* | | | | | | | | |
| Lateral Septum (LS) | **1,16** | ± | **0,04** | 0,93 | ± | 0,11 | 32,08769 | |
| Medial Septum (MS) | **1,11** | ± | **0,05** | 0,33* | ± | 0,05 | 2,3E-07* | Lost |
| *Diagonal Band of Broca* | | | | | | | | |
| Horizontal Band (HDB) | 0,62 | ± | 0,07 | **1,04** | ± | **0,07** | 0,313851 | |
| Vertical Band (VDB) | 0,89 | ± | 0,04 | 0,74 | ± | 0,06 | 21,56559 | |
| *Basal Ganglia* | | | | | | | | |
| Dorsolateral Striatum (DLST) | 0,70 | ± | 0,03 | 0,44 | ± | 0,06 | 0,689628 | |
| Ventromedial Striatum (VMST) | 0,67 | ± | 0,03 | 0,49 | ± | 0,04 | 1,455906 | |
| Substantia Nigra pars compacta (SNC) | 0,48 | ± | 0,06 | 0,63 | ± | 0,04 | 21,42251 | |
| Substantia Nigra pars reticulata (SNR) | 0,77 | ± | 0,02 | **0,94** | ± | **0,06** | 7,717153 | |
| *Mesolimbic* | | | | | | | | |
| Bed Nucleus of the Stria Terminalis (BST) | 0,75 | ± | 0,06 | **1,50** | ± | **0,19** | 0,507345 | |
| Nucleus Accumbens core (AcbC) | 0,68 | ± | 0,07 | **1,65*** | ± | **0,09** | 1,9E-05* | Gained |
| Nucleus Accumbens shell (AcbS) | 0,61 | ± | 0,08 | **1,33*** | ± | **0,13** | 0,074083 | |
| *Multimodal* | | | | | | | | |
| Dorsal Raphé (DR) | 0,44 | ± | 0,08 | 0,31 | ± | 0,04 | 82,82838 | |
| Dorsal Tegmental Nucleus (DTg) | **1,09** | ± | **0,04** | 0,37* | ± | 0,03 | 3,4E-09* | Lost |
| Lateral Habenula (LHab) | **1,30** | ± | **0,05** | **1,18** | ± | **0,04** | 43,66221 | |
| Locus Coeruleus (LC) | 0,42 | ± | 0,04 | 0,36 | ± | 0,03 | 151,379 | |
| Median Raphé (MR) | 0,73 | ± | 0,11 | 0,39 | ± | 0,04 | 3,746624 | |
| Periaqueductal Gray Matter (PAG) | 0,51 | ± | 0,05 | 0,44 | ± | 0,03 | 127,3212 | |

**Supplementary Table 1 – Functional Connectivity of the Suprachiasmatic Nucleus (rostral)**

Bold denotes 95% confidence interval of the VIP > 1.0; * denotes p<0.05 difference (unpaired Student’s t-test with Bonferroni correction)

| Region | Control | | | Shifted | | | p-value | Lost/Gained |
|---|---|---|---|---|---|---|---|---|
| | Mean | | SEM | Mean | | SEM | | |
| *Cortex* | | | | | | | | |
| Amigdalo-Piriform Cortex (APir) | 0,77 | ± | 0,03 | 0,60 | ± | 0,06 | 11,89134 | |
| Cingulate (Cg1) | **1,24** | ± | **0,06** | **1,59** | ± | **0,10** | 3,808004 | |
| Dorsolateral Orbital (DLO) | **1,43** | ± | **0,05** | 0,69* | ± | 0,05 | 6,6E-07* | Lost |
| Infralimbic (IL) | 0,92 | ± | 0,05 | 0,64 | ± | 0,05 | 0,704843 | |
| Lateral Entorhinal (LEC) | 0,82 | ± | 0,07 | 0,71 | ± | 0,04 | 79,74861 | |
| Lateral Orbital (LO) | **1,13** | ± | **0,08** | **1,51** | ± | **0,07** | 0,727839 | |
| Supplementary Motor Area (M2) | **1,32** | ± | **0,04** | **1,25** | ± | **0,08** | 205,6756 | |
| Medial Entorhinal (MEC) | **1,58** | ± | **0,06** | **1,38** | ± | **0,05** | 7,268696 | |
| Medial Orbital (MO) | 0,99 | ± | 0,04 | **1,16** | ± | **0,06** | 8,944203 | |
| Medial Prelimbic (mPrL) | 0,71 | ± | 0,06 | 0,74 | ± | 0,03 | 360,351 | |
| Perirhinal (PRH) | 0,86 | ± | 0,04 | 0,75 | ± | 0,04 | 34,8756 | |
| Prelimbic (PrL) | 0,66 | ± | 0,11 | 0,67 | ± | 0,05 | 459,8219 | |
| Retrosplenial (RSC) | 0,86 | ± | 0,07 | 0,64 | ± | 0,08 | 26,74441 | |
| Ventral Orbital (VO) | **1,18** | ± | **0,06** | **1,18** | ± | **0,09** | 467,5114 | |
| *Hippocampus* | | | | | | | | |
| Dorsal CA1(dCA1) | **1,58** | ± | **0,07** | 0,78* | ± | 0,08 | 0,0002* | Lost |
| Dorsal CA2 (dCA2) | 0,61 | ± | 0,08 | 0,81 | ± | 0,07 | 38,21555 | |
| Dorsal CA3 (dCA3) | **1,14** | ± | **0,07** | **1,51** | ± | **0,10** | 2,693435 | |
| Intermediate Dentate Gyrus (iDG) | **1,05** | ± | **0,05** | **1,19** | ± | **0,07** | 57,96148 | |
| Dorsal Hippocampus Molecular Layer (dML) | 0,79 | ± | 0,04 | 0,67 | ± | 0,08 | 102,2743 | |
| Dorsal Subiculum (dSub) | **1,68** | ± | **0,05** | 0,99* | ± | 0,05 | 1,8E-06* | Lost |
| Intermediate CA1 (iCA1) | **1,03** | ± | **0,04** | **1,30*** | ± | **0,04** | 0,0393* | Increased |
| Ventral CA1 (vCA1) | 0,77 | ± | 0,04 | 0,65 | ± | 0,06 | 40,87393 | |
| Ventral CA2 (vCA2) | 0,80 | ± | 0,04 | 0,68 | ± | 0,06 | 43,91868 | |
| Ventral CA3 (vCA3) | **1,27** | ± | **0,06** | **1,11** | ± | **0,05** | 34,68662 | |
| Ventral Hippocampus Molecular Layer (vML) | 0,77 | ± | 0,04 | 0,79 | ± | 0,03 | 329,958 | |
| *Circadian Rhythms* | | | | | | | | |
| Anterior Hypothalamus AHA | **1,13** | ± | **0,07** | 0,56* | ± | 0,08 | 0,0164* | Lost |
| Arcuate Nucleus (ARC) | 0,62 | ± | 0,05 | 0,78 | ± | 0,13 | 136,9478 | |
| Suprachiasmatic Nucleus - Rostral (rSCN) | **1,11** | ± | **0,09** | 0,88 | ± | 0,08 | 31,32833 | |
| Suprachiasmatic Nucleus - Caudal (cSCN) | **1,50** | ± | **0,10** | 0,71* | ± | 0,07 | 0,0014* | Lost |
| Dorsomedial Hypothalamus (DMH) | 0,96 | ± | 0,05 | 0,52* | ± | 0,04 | 0,0006* | |
| Intergeniculate Leaftlet (IGL) | 0,66 | ± | 0,03 | 0,56 | ± | 0,07 | 99,7046 | |
| Lateral Hypothalamus (LH) | 0,72 | ± | 0,08 | 0,78 | ± | 0,06 | 257,5517 | |
| Medial Preoptic Nucleus - Caudal (cMPO) | **1,38** | ± | **0,07** | 0,79* | ± | 0,06 | 0,0019* | Lost |
| Medial Preoptic Nucleus - Rostral (rMPO) | 0,98 | ± | 0,10 | 0,53 | ± | 0,06 | 0,601298 | |
| Median Eminence (ME) | 0,88 | ± | 0,06 | 1,03 | ± | 0,11 | 126,9113 | |
| Paraventricular Hypothalamic Anterior - Parvicellular (PVH) | **1,10** | ± | **0,05** | 0,89 | ± | 0,04 | 0,987764 | |
| Paraventricular Thalamic Nucleus (PVT) | 0,61 | ± | 0,10 | 0,80 | ± | 0,09 | 84,63398 | |
| Pineal Gland | **1,06** | ± | **0,02** | 0,57* | ± | 0,06 | 6,1E-05* | Lost |
| Retrochiasmatic Area (ReCh) | **1,17** | ± | **0,09** | 0,75 | ± | 0,09 | 1,361238 | |
| Subparaventricular Zone (SPZ) | 0,67 | ± | 0,06 | 0,56 | ± | 0,06 | 100,9372 | |
| Supraoptic Nucleus (SO) | 0,99 | ± | 0,06 | **1,22** | ± | **0,06** | 8,4198 | |
| Ventrolateral Preoptic Nucleus (VLPO) | 0,83 | ± | 0,04 | 0,71 | ± | 0,06 | 60,93023 | |
| Ventromedial Hypothalamus (VMH) | 0,94 | ± | 0,04 | 0,68 | ± | 0,07 | 2,35402 | |
| Ventromedial Preoptic Nucleus (VMPO) | **1,17** | ± | **0,03** | 0,68* | ± | 0,06 | 0,0003* | Lost |
| *Thalamus* | | | | | | | | |
| Mediodorsal (MD) | 0,71 | ± | 0,04 | 0,77 | ± | 0,11 | 299,5734 | |
| Reuniens (RE) | **1,14** | ± | **0,07** | **1,05** | ± | **0,09** | 205,9862 | |
| Ventral Posterolateral (VPL) | 0,58 | ± | 0,07 | **1,89*** | ± | **0,05** | 5,7E-10* | Gained |
| *Amygdala* | | | | | | | | |
| Basolateral (BLA) | 0,56 | ± | 0,05 | 0,53 | ± | 0,06 | 353,3005 | |
| Central (CA) | 0,85 | ± | 0,03 | 0,93 | ± | 0,03 | 44,21471 | |
| Medial (MA) | 0,70 | ± | 0,04 | 0,50 | ± | 0,06 | 6,768114 | |
| *Septum* | | | | | | | | |
| Lateral Septum (LS) | **1,20** | ± | **0,04** | 1,02 | ± | 0,08 | 32,39167 | |
| Medial Septum (MS) | 0,61 | ± | 0,05 | 0,69 | ± | 0,08 | 195,7477 | |
| *Diagonal Band of Broca* | | | | | | | | |
| Horizontal Band (HDB) | 0,48 | ± | 0,04 | **1,04*** | ± | **0,09** | 0,0097* | Gained |
| Vertical Band (VDB) | 0,89 | ± | 0,06 | 0,68 | ± | 0,05 | 8,550372 | |
| *Basal Ganglia* | | | | | | | | |
| Dorsolateral Striatum (DLST) | 0,62 | ± | 0,05 | **1,85*** | ± | **0,08** | 3,4E-08* | Gained |
| Ventromedial Striatum (VMST) | 0,65 | ± | 0,05 | **1,65*** | ± | **0,09** | 2,5E-06 | Gained |
| Substantia Nigra pars compacta (SNC) | **1,16** | ± | **0,06** | **1,21** | ± | **0,06** | 247,9771 | |
| Substantia Nigra pars reticulata (SNR) | 0,73 | ± | 0,04 | 0,98 | ± | 0,06 | 1,474006 | |
| *Mesolimbic* | | | | | | | | |
| Bed Nucleus of the Stria Terminalis (BST) | 0,73 | ± | 0,07 | **1,25** | ± | **0,10** | 0,26159 | |
| Nucleus Accumbens core (AcbC) | 0,54 | ± | 0,04 | 0,63 | ± | 0,04 | 61,65113 | |
| Nucleus Accumbens shell (AcbS) | 0,64 | ± | 0,04 | **1,27*** | ± | **0,04** | 3,1E-07* | Gained |
| *Multimodal* | | | | | | | | |
| Dorsal Raphé (DR) | 0,68 | ± | 0,05 | **1,16*** | ± | **0,07** | 0,0039* | Gained |
| Dorsal Tegmental Nucleus (DTg) | 0,97 | ± | 0,06 | **1,11** | ± | **0,06** | 53,73059 | |
| Lateral Habenula (LHab) | **1,34** | ± | **0,05** | 0,80* | ± | 0,06 | 0,0007* | Lost |
| Locus Coeruleus (LC) | 0,97 | ± | 0,04 | **1,46*** | ± | **0,05** | 4,2E-05* | Gained |
| Median Raphé (MR) | **1,42** | ± | **0,05** | 0,74* | ± | 0,07 | 5,2E-05* | Lost |
| Periaqueductal Gray Matter (PAG) | 0,70 | ± | 0,04 | 0,77 | ± | 0,06 | 152,6102 | |

**Supplementary Table 2 – Functional Connectivity of the Rostrodorsal Dentate Gyrus**

Bold denotes 95% confidence interval of the VIP > 1.0; * denotes p<0.05 difference (unpaired Student's t-test with Bonferroni correction)

| Region | Control Mean | | Control SEM | Shifted Mean | | Shifted SEM | p-value | Lost/Gained |
|---|---|---|---|---|---|---|---|---|
| *Cortex* | | | | | | | | |
| Amigdalo-Piriform Cortex (APir) | **1,64** | ± | **0,01** | **1,24*** | ± | **0,02** | 2,2E-09* | Decreased |
| Cingulate (Cg1) | **1,06** | ± | **0,05** | 0,30* | ± | 0,04 | 1,7E-07* | Lost |
| Dorsolateral Orbital (DLO) | 0,53 | ± | 0,04 | 0,92* | ± | 0,04 | 3,6E-04* | |
| Infralimbic (IL) | **1,19** | ± | **0,02** | **1,21** | ± | **0,03** | 318,0462 | |
| Lateral Entorhinal (LEC) | **1,42** | ± | **0,02** | 0,92* | ± | 0,04 | 7,9E-08* | Lost |
| Lateral Orbital (LO) | 0,58 | ± | 0,04 | 0,84 | ± | 0,05 | 0,183161 | |
| Supplementary Motor Area (M2) | 0,77 | ± | 0,06 | **1,37*** | ± | **0,03** | 4,2E-06* | Gained |
| Medial Entorhinal (MEC) | **1,44** | ± | **0,01** | 0,89* | ± | 0,04 | 2,5E-09* | Lost |
| Medial Orbital (MO) | 0,28 | ± | 0,03 | **1,30*** | ± | **0,03** | 1,8E-13* | Gained |
| Medial Prelimbic (mPrL) | **1,12** | ± | **0,02** | 0,91* | ± | 0,03 | 8,8E-03* | Lost |
| Perirhinal (PRH) | 0,97 | ± | 0,05 | **1,08** | ± | **0,04** | 60,75625 | |
| Prelimbic (PrL) | 0,72 | ± | 0,04 | 0,99* | ± | 0,03 | 0,0223* | |
| Retrosplenial (RSC) | 0,80 | ± | 0,05 | 0,45* | ± | 0,04 | 0,0104* | |
| Ventral Orbital (VO) | 0,33 | ± | 0,01 | 1,02* | ± | 0,06 | 6,7E-07* | |
| *Hippocampus* | | | | | | | | |
| Dorsal CA1(dCA1) | **1,22** | ± | **0,03** | 0,42* | ± | 0,02 | 3,0E-13* | Lost |
| Dorsal CA2 (dCA2) | 0,73 | ± | 0,05 | 0,49 | ± | 0,04 | 0,304259 | |
| Dorsal CA3 (dCA3) | **1,03** | ± | 0,07 | 0,48* | ± | 0,07 | 0,0125* | Lost |
| Dorsal Dentate Gyrus - Rostral (rdDG) | 0,70 | ± | 0,05 | 0,99 | ± | 0,06 | 0,88208 | |
| Dorsal Hippocampus Molecular Layer (dML) | 0,85 | ± | 0,05 | 0,46* | ± | 0,03 | 0,0014* | |
| Dorsal Subiculum (dSub) | **1,34** | ± | **0,02** | **1,36** | ± | **0,03** | 353,4168 | |
| Intermediate CA1 (iCA1) | **1,49** | ± | **0,02** | **1,74*** | ± | **0,02** | 5,5E-06* | Increased |
| Ventral CA1 (vCA1) | **1,46** | ± | **0,02** | **1,56** | ± | **0,02** | 0,829384 | |
| Ventral CA2 (vCA2) | **1,67** | ± | **0,02** | **1,59** | ± | **0,02** | 4,537402 | |
| Ventral CA3 (vCA3) | **1,64** | ± | **0,02** | **1,79*** | ± | **0,02** | 0,0098* | Increased |
| Ventral Hippocampus Molecular Layer (vML) | **1,43** | ± | **0,02** | **1,76*** | ± | **0,02** | 2,1E-07* | Increased |
| *Circadian Rhythms* | | | | | | | | |
| Anterior Hypothalamus AHA | 0,22 | ± | 0,08 | **1,50*** | ± | **0,04** | 1,7E-09* | Gained |
| Arcuate Nucleus (ARC) | **1,18** | ± | **0,02** | 1,00 | ± | 0,06 | 6,935221 | |
| Suprachiasmatic Nucleus - Rostral (rSCN) | 0,34 | ± | 0,04 | 0,86* | ± | 0,04 | 7,7E-06* | |
| Suprachiasmatic Nucleus - Caudal (cSCN) | 0,33 | ± | 0,05 | **1,37*** | ± | **0,04** | 1,5E-10* | Gained |
| Dorsomedial Hypothalamus (DMH) | **1,45** | ± | **0,01** | 0,38* | ± | 0,06 | 1,1E-10* | Lost |
| Intergeniculate Leaflet (IGL) | **1,42** | ± | **0,02** | 0,97* | ± | 0,04 | 2,1E-07* | Lost |
| Lateral Hypothalamus (LH) | 0,58 | ± | 0,07 | 0,61 | ± | 0,04 | 369,1971 | |
| Medial Preoptic Nucleus - Caudal (cMPO) | 0,29 | ± | 0,03 | **1,20*** | ± | **0,06** | 1,6E-09* | Gained |
| Medial Preoptic Nucleus - Rostral (rMPO) | 0,96 | ± | 0,04 | 0,83 | ± | 0,07 | 58,55919 | |
| Median Eminence (ME) | 0,84 | ± | 0,03 | 0,68 | ± | 0,06 | 7,210105 | |
| Paraventricular Hypothalamic Anterior - Parvicellular (PVH) | 0,73 | ± | 0,06 | 0,98 | ± | 0,07 | 4,64645 | |
| Paraventricular Thalamic Nucleus (PVT) | 0,67 | ± | 0,04 | 0,68 | ± | 0,05 | 454,5827 | |
| Pineal Gland | **1,13** | ± | **0,04** | 0,79 | ± | 0,07 | 0,186143 | |
| Retrochiasmatic Area (ReCh) | 0,34 | ± | 0,05 | 0,45 | ± | 0,06 | 81,41836 | |
| Subparaventricular Zone (SPZ) | 0,56 | ± | 0,05 | **1,16*** | ± | **0,06** | 0,0002* | Gained |
| Supraoptic Nucleus (SO) | 0,43 | ± | 0,01 | **1,06*** | ± | **0,04** | 6,5E-10* | Gained |
| Ventrolateral Preoptic Nucleus (VLPO) | **1,24** | ± | **0,02** | 0,57* | ± | 0,04 | 8,7E-10* | Lost |
| Ventromedial Hypothalamus (VMH) | **1,28** | ± | **0,03** | 0,57* | ± | 0,07 | 1,4E-05* | Lost |
| Ventromedial Preoptic Nucleus (VMPO) | 0,60 | ± | 0,04 | 0,85* | ± | 0,03 | 0,0377* | |
| *Thalamus* | | | | | | | | |
| Mediodorsal (MD) | 0,33 | ± | 0,07 | 0,58 | ± | 0,05 | 4,691224 | |
| Reuniens (RE) | 0,86 | ± | 0,05 | **1,04** | ± | **0,05** | 6,973222 | |
| Ventral Posterolateral (VPL) | 0,43 | ± | 0,07 | 0,44 | ± | 0,04 | 444,0629 | |
| *Amygdala* | | | | | | | | |
| Basolateral (BLA) | 0,43 | ± | 0,07 | 0,84 | ± | 0,06 | 0,059377 | |
| Central (CA) | **1,02** | ± | **0,03** | 0,79 | ± | 0,04 | 0,083182 | |
| Medial (MA) | **1,31** | ± | **0,03** | **1,37** | ± | **0,03** | 79,32266 | |
| *Septum* | | | | | | | | |
| Lateral Septum (LS) | 0,47 | ± | 0,02 | 0,82* | ± | 0,05 | 0,0008* | |
| Medial Septum (MS) | 0,48 | ± | 0,03 | 0,62 | ± | 0,03 | 1,198276 | |
| *Diagonal Band of Broca* | | | | | | | | |
| Horizontal Band (HDB) | 0,35 | ± | 0,02 | 0,48 | ± | 0,06 | 26,2389 | |
| Vertical Band (VDB) | 0,91 | ± | 0,03 | 0,53* | ± | 0,03 | 8,2E-06* | |
| *Basal Ganglia* | | | | | | | | |
| Dorsolateral Striatum (DLST) | 0,75 | ± | 0,03 | **1,02** | ± | 0,06 | 0,229626 | |
| Ventromedial Striatum (VMST) | **1,08** | ± | **0,04** | 0,94 | ± | 0,07 | 33,43139 | |
| Substantia Nigra pars compacta (SNC) | **1,65** | ± | **0,01** | **1,51*** | ± | **0,02** | 0,0407* | Decreased |
| Substantia Nigra pars reticulata (SNR) | **1,44** | ± | **0,02** | 0,97* | ± | 0,03 | 2,8E-08* | Lost |
| *Mesolimbic* | | | | | | | | |
| Bed Nucleus of the Stria Terminalis (BST) | 0,72 | ± | 0,06 | 0,79 | ± | 0,06 | 204,369 | |
| Nucleus Accumbens core (AcbC) | 0,75 | ± | 0,03 | 0,70 | ± | 0,09 | 259,7042 | |
| Nucleus Accumbens shell (AcbS) | 0,62 | ± | 0,04 | 0,80 | ± | 0,06 | 7,956568 | |
| *Multimodal* | | | | | | | | |
| Dorsal Raphé (DR) | 0,69 | ± | 0,04 | **1,19*** | ± | **0,03** | 2,5E-06* | Gained |
| Dorsal Tegmental Nucleus (DTg) | 0,37 | ± | 0,02 | 0,69 | ± | 0,07 | 0,079947 | |
| Lateral Habenula (LHab) | 0,68 | ± | 0,03 | 0,80 | ± | 0,05 | 19,56333 | |
| Locus Coeruleus (LC) | **1,17** | ± | **0,02** | **1,03** | ± | **0,06** | 15,85993 | |
| Median Raphé (MR) | 0,74 | ± | 0,06 | 0,84 | ± | 0,06 | 134,1981 | |
| Periaqueductal Gray Matter (PAG) | **1,51** | ± | **0,02** | 0,82* | ± | 0,05 | 7,9E-09* | Lost |

**Supplementary Table 3 – Functional Connectivity of the Intermediate Dentate Gyrus**

Bold denotes 95% confidence interval of the VIP > 1.0; * denotes p<0.05 difference (unpaired Student's t-test with Bonferroni correction).

| Region | Control Mean | | Control SEM | Shifted Mean | | Shifted SEM | p-value | Lost/Gained |
|---|---|---|---|---|---|---|---|---|
| *Cortex* | | | | | | | | |
| Amigdalo-Piriform Cortex (APir) | **1,11** | ± | **0,02** | 0,96 | ± | 0,03 | 0,074038 | |
| Cingulate (Cg1) | 0,85 | ± | 0,05 | 0,39* | ± | 0,04 | 0,0007* | |
| Dorsolateral Orbital (DLO) | 0,63 | ± | 0,02 | **1,14*** | ± | **0,04** | 2,6E-07* | Gained |
| Infralimbic (IL) | **1,48** | ± | **0,02** | **1,32*** | ± | **0,02** | 0,0103* | Decreased |
| Lateral Entorhinal (LEC) | **1,13** | ± | **0,02** | 0,75* | ± | 0,03 | 3,1E-06* | Lost |
| Lateral Orbital (LO) | 0,53 | ± | 0,05 | 0,60 | ± | 0,03 | 108,3309 | |
| Supplementary Motor Area (M2) | 0,71 | ± | 0,03 | **0,97*** | ± | **0,04** | 0,0075* | Gained |
| Medial Orbital (MO) | 0,81 | ± | 0,02 | **1,17*** | ± | **0,02** | 9,8E-07* | Gained |
| Medial Prelimbic (mPrL) | **1,33** | ± | **0,02** | **1,12*** | ± | **0,03** | 0,0042* | Decreased |
| Perirhinal (PRH) | **1,15** | ± | **0,03** | 0,80* | ± | 0,03 | 0,0001* | Lost |
| Prelimbic (PrL) | 0,96 | ± | 0,05 | 0,83 | ± | 0,04 | 25,31849 | |
| Retrosplenial (RSC) | 1,00 | ± | 0,06 | 0,84 | ± | 0,05 | 30,83428 | |
| Ventral Orbital (VO) | 0,58 | ± | 0,02 | 0,98* | ± | 0,08 | 0,0375* | |
| *Hippocampus* | | | | | | | | |
| Dorsal CA1(dCA1) | **1,22** | ± | **0,04** | **1,13** | ± | **0,05** | 72,64147 | |
| Dorsal CA2 (dCA2) | **1,11** | ± | **0,05** | 1,00 | ± | 0,06 | 64,0889 | |
| Dorsal CA3 (dCA3) | **1,13** | ± | **0,06** | **1,21** | ± | **0,05** | 155,062 | |
| Dorsal Dentate Gyrus - Rostral (rdDG) | **1,37** | ± | **0,05** | **1,30** | ± | **0,05** | 175,8706 | |
| Intermediate Dentate Gyrus (iDG) | **1,47** | ± | **0,01** | **1,05*** | ± | **0,03** | 8,4E-09* | Decreased |
| Dorsal Hippocampus Molecular Layer (dML) | **1,21** | ± | **0,03** | 0,84* | ± | 0,06 | 0,0060* | Lost |
| Dorsal Subiculum (dSub) | **1,44** | ± | **0,03** | 0,94* | ± | 0,03 | 1,4E-07* | Lost |
| Intermediate CA1 (iCA1) | **1,01** | ± | **0,04** | **1,18** | ± | **0,01** | 0,18658 | |
| Ventral CA1 (vCA1) | **1,09** | ± | **0,02** | **1,16** | ± | **0,03** | 52,61082 | |
| Ventral CA2 (vCA2) | **1,21** | ± | **0,02** | **1,31*** | ± | **0,01** | 0,0299* | Increased |
| Ventral CA3 (vCA3) | **1,49** | ± | **0,02** | **1,16*** | ± | **0,03** | 2E-05* | Decreased |
| Ventral Hippocampus Molecular Layer (vML) | **1,17** | ± | **0,02** | 0,94* | ± | 0,02 | 0,0011* | Lost |
| *Circadian Rhythms* | | | | | | | | |
| Anterior Hypothalamus AHA | 0,93 | ± | 0,06 | **1,07** | ± | **0,04** | 31,77941 | |
| Arcuate Nucleus (ARC) | 0,96 | ± | 0,02 | 0,45* | ± | 0,07 | 0,0007* | |
| Suprachiasmatic Nucleus - Rostral (rSCN) | 0,66 | ± | 0,07 | 0,62 | ± | 0,07 | 349,3967 | |
| Suprachiasmatic Nucleus - Caudal (cSCN) | 0,69 | ± | 0,03 | **1,15*** | ± | **0,04** | 5,2E-06* | Gained |
| Dorsomedial Hypothalamus (DMH) | **1,33** | ± | **0,02** | 0,37* | ± | 0,03 | 8,4E-15* | Lost |
| Intergeniculate Leaftlet (IGL) | **1,09** | ± | **0,01** | 0,75* | ± | 0,04 | 2,2E-05* | Lost |
| Lateral Hypothalamus (LH) | 0,39 | ± | 0,09 | 0,74 | ± | 0,05 | 1,429953 | |
| Medial Preoptic Nucleus - Caudal (cMPO) | 0,99 | ± | 0,04 | **1,72*** | ± | **0,04** | 2E-09* | Gained |
| Medial Preoptic Nucleus - Rostral (rMPO) | 0,80 | ± | 0,05 | 0,85 | ± | 0,05 | 247,0587 | |
| Median Eminence (ME) | 0,82 | ± | 0,02 | 0,41* | ± | 0,05 | 0,0005* | |
| Paraventricular Hypothalamic Anterior - Parvicellular (PVH) | 0,73 | ± | 0,02 | **1,03*** | ± | **0,05** | 0,0037* | Gained |
| Paraventricular Thalamic Nucleus (PVT) | **1,03** | ± | **0,05** | 1,02 | ± | 0,07 | 434,4526 | |
| Pineal Gland | 1,00 | ± | 0,05 | 0,97 | ± | 0,07 | 360,6744 | |
| Retrochiasmatic Area (ReCh) | 0,86 | ± | 0,04 | 0,46* | ± | 0,06 | 0,0168* | |
| Subparaventricular Zone (SPZ) | 0,57 | ± | 0,03 | 0,91* | ± | 0,04 | 0,0008* | |
| Supraoptic Nucleus (SO) | 0,47 | ± | 0,03 | **1,08*** | ± | **0,04** | 1,6E-08* | Gained |
| Ventrolateral Preoptic Nucleus (VLPO) | **1,18** | ± | **0,02** | 0,45* | ± | 0,03 | 5,9E-13* | Lost |
| Ventromedial Hypothalamus (VMH) | **1,09** | ± | **0,03** | 0,40* | ± | 0,08 | 8,1E-05* | Lost |
| Ventromedial Preoptic Nucleus (VMPO) | 0,71 | ± | 0,03 | **1,03*** | ± | **0,03** | 4,8E-05* | Gained |
| *Thalamus* | | | | | | | | |
| Mediodorsal (MD) | 0,71 | ± | 0,04 | **1,08** | ± | **0,09** | 0,491915 | |
| Reuniens (RE) | **1,11** | ± | **0,04** | 0,64* | ± | 0,04 | 3,7E-05* | Lost |
| Ventral Posterolateral (VPL) | 0,51 | ± | 0,04 | **1,19*** | ± | **0,05** | 2,2E-06* | Gained |
| *Amygdala* | | | | | | | | |
| Basolateral (BLA) | 0,36 | ± | 0,04 | 0,55 | ± | 0,05 | 2,182046 | |
| Central (CA) | **1,11** | ± | **0,03** | **1,02** | ± | **0,03** | 32,77601 | |
| Medial (MA) | **1,12** | ± | **0,03** | **1,08** | ± | **0,02** | 94,38153 | |
| *Septum* | | | | | | | | |
| Lateral Septum (LS) | 0,48 | ± | 0,03 | 0,72 | ± | 0,05 | 0,348771 | |
| Medial Septum (MS) | 0,52 | ± | 0,04 | 0,36 | ± | 0,02 | 0,818654 | |
| *Diagonal Band of Broca* | | | | | | | | |
| Horizontal Band (HDB) | 0,97 | ± | 0,03 | 0,53* | ± | 0,07 | 0,0041* | |
| Vertical Band (VDB) | 0,55 | ± | 0,03 | 0,44 | ± | 0,06 | 50,23243 | |
| *Basal Ganglia* | | | | | | | | |
| Dorsolateral Striatum (DLST) | 0,37 | ± | 0,03 | 0,62 | ± | 0,06 | 0,504237 | |
| Ventromedial Striatum (VMST) | 0,68 | ± | 0,06 | 0,66 | ± | 0,05 | 424,9658 | |
| Substantia Nigra pars compacta (SNC) | **1,54** | ± | **0,01** | **1,12*** | ± | **0,02** | 2,6E-10* | Decreased |
| Substantia Nigra pars reticulata (SNR) | **1,34** | ± | **0,04** | 0,79* | ± | 0,05 | 8,5E-06* | Lost |
| *Mesolimbic* | | | | | | | | |
| Bed Nucleus of the Stria Terminalis (BST) | 0,72 | ± | 0,04 | 0,41 | ± | 0,10 | 4,82201 | |
| Nucleus Accumbens core (AcbC) | **1,03** | ± | **0,02** | 0,63* | ± | 0,03 | 9,8E-07* | Lost |
| Nucleus Accumbens shell (AcbS) | **1,05** | ± | **0,03** | 0,75* | ± | 0,04 | 0,0021* | Lost |
| *Multimodal* | | | | | | | | |
| Dorsal Raphé (DR) | 0,93 | ± | 0,02 | 0,99 | ± | 0,04 | 109,5642 | |
| Dorsal Tegmental Nucleus (DTg) | 0,38 | ± | 0,04 | **1,33*** | ± | **0,05** | 1,5E-09* | Gained |
| Lateral Habenula (LHab) | 0,55 | ± | 0,02 | 0,91* | ± | 0,07 | 0,0356* | |
| Locus Coeruleus (LC) | 0,80 | ± | 0,03 | **1,26*** | ± | **0,08** | 0,0084* | Gained |
| Median Raphé (MR) | **1,34** | ± | **0,04** | **1,78*** | ± | **0,04** | 0,0001* | Increased |
| Periaqueductal Gray Matter (PAG) | **1,07** | ± | **0,02** | **1,65*** | ± | **0,05** | 6,7E-07* | Increased |

**Supplementary Table 4 – Functional Connectivity of the Medial Entorhinal Cortex**

Bold denotes 95% confidence interval of the VIP > 1.0; * denotes p<0.05 difference (unpaired Student's t-test with Bonferroni correction).

| Region | Control Mean | ± | Control SEM | Shifted Mean | ± | Shifted SEM | p-value | Lost/Gained |
|---|---|---|---|---|---|---|---|---|
| *Cortex* | | | | | | | | |
| Amigdalo-Piriform Cortex (APir) | 0,88 | ± | 0,05 | **1,46*** | ± | **0,02** | 3,7E-07* | Gained |
| Cingulate (Cg1) | 1,00 | ± | 0,06 | 0,69 | ± | 0,04 | 0,118093 | |
| Dorsolateral Orbital (DLO) | 0,85 | ± | 0,05 | 0,94 | ± | 0,03 | 66,37011 | |
| Infralimbic (IL) | 0,99 | ± | 0,02 | **1,05** | ± | **0,01** | 27,62106 | |
| Lateral Entorhinal (LEC) | 0,96 | ± | 0,04 | **1,56*** | ± | **0,03** | 5,8E-08* | Gained |
| Lateral Orbital (LO) | 0,76 | ± | 0,06 | 0,56 | ± | 0,06 | 18,63667 | |
| Supplementary Motor Area (M2) | 0,73 | ± | 0,09 | **1,07** | ± | **0,03** | 0,704968 | |
| Medial Entorhinal (MEC) | **1,30** | ± | **0,02** | 0,73* | ± | 0,03 | 1,1E-09* | Lost |
| Medial Orbital (MO) | **1,53** | ± | **0,07** | **1,03*** | ± | **0,02** | 0,0009* | Decreased |
| Medial Prelimbic (mPrL) | **1,01** | ± | **0,02** | 0,88* | ± | 0,02 | 0,0344* | Lost |
| Prelimbic (PrL) | 0,92 | ± | 0,03 | 1,00 | ± | 0,02 | 29,10254 | |
| Retrosplenial (RSC) | 0,76 | ± | 0,05 | **1,29*** | ± | **0,03** | 2,3E-06* | Gained |
| Ventral Orbital (VO) | 1,00 | ± | 0,08 | **1,09** | ± | **0,03** | 132,515 | |
| *Hippocampus* | | | | | | | | |
| Dorsal CA1(dCA1) | **1,58** | ± | **0,03** | 0,31* | ± | 0,04 | 2,4E-14* | Lost |
| Dorsal CA2 (dCA2) | 0,97 | ± | 0,05 | 0,68 | ± | 0,07 | 1,606866 | |
| Dorsal CA3 (dCA3) | **1,34** | ± | **0,09** | 0,70* | ± | 0,03 | 0,0004* | Lost |
| Dorsal Dentate Gyrus - Rostral (rdDG) | 0,84 | ± | 0,04 | 0,64 | ± | 0,03 | 0,739192 | |
| Intermediate Dentate Gyrus (iDG) | **1,17** | ± | **0,04** | **1,13** | ± | **0,03** | 197,7391 | |
| Dorsal Hippocampus Molecular Layer (dML) | **1,70** | ± | **0,03** | 0,61* | ± | 0,06 | 2,6E-10* | Lost |
| Dorsal Subiculum (dSub) | 0,80 | ± | 0,04 | **1,10*** | ± | **0,01** | 0,0004* | Gained |
| Intermediate CA1 (iCA1) | 0,83 | ± | 0,06 | **1,11** | ± | **0,03** | 0,435136 | |
| Ventral CA1 (vCA1) | 0,94 | ± | 0,03 | **1,63*** | ± | **0,02** | 3,3E-12* | Gained |
| Ventral CA2 (vCA2) | 0,99 | ± | 0,05 | **1,49*** | ± | **0,02** | 4,6E-06* | Gained |
| Ventral CA3 (vCA3) | 1,01 | ± | 0,05 | **1,17** | ± | **0,03** | 5,257298 | |
| Ventral Hippocampus Molecular Layer (vML) | **1,07** | ± | **0,01** | **1,38*** | ± | **0,02** | 2,5E-07* | Increased |
| *Circadian Rhythms* | | | | | | | | |
| Anterior Hypothalamus AHA | 0,51 | ± | 0,04 | 0,90* | ± | 0,04 | 0,0009* | |
| Arcuate Nucleus (ARC) | 0,91 | ± | 0,03 | **1,24*** | ± | **0,04** | 0,0033* | Gained |
| Suprachiasmatic Nucleus - Rostral (rSCN) | 0,71 | ± | 0,04 | 0,50 | ± | 0,04 | 0,338461 | |
| Suprachiasmatic Nucleus - Caudal (cSCN) | 0,97 | ± | 0,06 | 0,90 | ± | 0,07 | 214,8404 | |
| Dorsomedial Hypothalamus (DMH) | **1,19** | ± | **0,01** | **1,66*** | ± | **0,06** | 0,0001* | Increased |
| Intergeniculate Leaftlet (IGL) | 0,99 | ± | 0,03 | **1,19*** | ± | **0,03** | 0,0106* | Gained |
| Lateral Hypothalamus (LH) | 0,45 | ± | 0,08 | 0,73 | ± | 0,04 | 1,784527 | |
| Medial Preoptic Nucleus - Caudal (cMPO) | **1,15** | ± | **0,06** | 1,01 | ± | 0,03 | 29,91208 | |
| Medial Preoptic Nucleus - Rostral (rMPO) | 0,97 | ± | 0,04 | 0,72 | ± | 0,08 | 4,81487 | |
| Median Eminence (ME) | 0,90 | ± | 0,03 | 0,66 | ± | 0,06 | 0,630929 | |
| Paraventricular Hypothalamic Anterior - Parvicellular (PVH) | 0,75 | ± | 0,08 | 0,74 | ± | 0,04 | 454,6048 | |
| Paraventricular Thalamic Nucleus (PVT) | 0,79 | ± | 0,04 | **1,07** | ± | **0,08** | 2,119928 | |
| Pineal Gland | **1,47** | ± | **0,04** | 0,94* | ± | 0,06 | 0,0002* | Lost |
| Retrochiasmatic Area (ReCh) | **1,35** | ± | **0,07** | 0,62* | ± | 0,04 | 1,9E-05* | Lost |
| Subparaventricular Zone (SPZ) | 0,78 | ± | 0,05 | 0,86 | ± | 0,06 | 134,4916 | |
| Supraoptic Nucleus (SO) | 0,85 | ± | 0,07 | **1,20** | ± | **0,06** | 0,357318 | |
| Ventrolateral Preoptic Nucleus (VLPO) | 0,99 | ± | 0,02 | **1,18** | ± | **0,06** | 3,925039 | |
| Ventromedial Hypothalamus (VMH) | 0,97 | ± | 0,04 | **1,18** | ± | **0,05** | 2,319945 | |
| Ventromedial Preoptic Nucleus (VMPO) | 0,76 | ± | 0,03 | **1,22*** | ± | **0,04** | 8,6E-06* | Gained |
| *Thalamus* | | | | | | | | |
| Mediodorsal (MD) | **1,35** | ± | **0,06** | **1,22** | ± | **0,06** | 70,70184 | |
| Reuniens (RE) | 0,77 | ± | 0,03 | 0,76 | ± | 0,03 | 387,5601 | |
| Ventral Posterolateral (VPL) | **1,19** | ± | **0,04** | 0,53* | ± | 0,03 | 6,5E-08* | Lost |
| *Amygdala* | | | | | | | | |
| Basolateral (BLA) | 0,64 | ± | 0,05 | **1,10** | ± | **0,09** | 0,134321 | |
| Central (CA) | 0,97 | ± | 0,03 | **1,26*** | ± | **0,03** | 0,0009* | Gained |
| Medial (MA) | 0,94 | ± | 0,02 | **1,52*** | ± | **0,02** | 1,2E-11* | Gained |
| *Septum* | | | | | | | | |
| Lateral Septum (LS) | 0,69 | ± | 0,08 | 0,85 | ± | 0,05 | 45,50253 | |
| Medial Septum (MS) | 0,92 | ± | 0,10 | 0,81 | ± | 0,03 | 148,9929 | |
| *Diagonal Band of Broca* | | | | | | | | |
| Horizontal Band (HDB) | 0,95 | ± | 0,07 | 0,70 | ± | 0,04 | 2,719208 | |
| Vertical Band (VDB) | **1,07** | ± | **0,07** | 0,78 | ± | 0,02 | 0,272261 | |
| *Basal Ganglia* | | | | | | | | |
| Dorsolateral Striatum (DLST) | 0,48 | ± | 0,07 | 0,50 | ± | 0,02 | 372,8079 | |
| Ventromedial Striatum (VMST) | 0,63 | ± | 0,06 | 0,40 | ± | 0,04 | 1,267786 | |
| Substantia Nigra pars compacta (SNC) | **1,19** | ± | **0,04** | **1,20** | ± | **0,04** | 432,697 | |
| Substantia Nigra pars reticulata (SNR) | **1,02** | ± | **0,04** | **1,51*** | ± | **0,07** | 0,0023* | Increased |
| *Mesolimbic* | | | | | | | | |
| Bed Nucleus of the Stria Terminalis (BST) | 0,81 | ± | 0,08 | 0,46 | ± | 0,08 | 2,501214 | |
| Nucleus Accumbens core (AcbC) | 0,87 | ± | 0,03 | 0,61* | ± | 0,02 | 0,0014* | |
| Nucleus Accumbens shell (AcbS) | 0,88 | ± | 0,06 | 0,54 | ± | 0,03 | 0,05385 | |
| *Multimodal* | | | | | | | | |
| Dorsal Raphé (DR) | 0,69 | ± | 0,04 | 0,91 | ± | 0,04 | 0,795011 | |
| Dorsal Tegmental Nucleus (DTg) | 0,85 | ± | 0,07 | 0,92 | ± | 0,07 | 229,0388 | |
| Lateral Habenula (LHab) | 0,57 | ± | 0,06 | 0,69 | ± | 0,05 | 51,97868 | |
| Locus Coeruleus (LC) | 0,81 | ± | 0,05 | 0,87 | ± | 0,04 | 223,9133 | |
| Median Raphé (MR) | **1,16** | ± | **0,06** | 0,71 | ± | 0,07 | 0,056796 | |
| Periaqueductal Gray Matter (PAG) | 0,94 | ± | 0,05 | 0,74 | ± | 0,04 | 2,033008 | |

**Supplementary Table 5 – Functional Connectivity of the Perirhinal Cortex**

Bold denotes 95% confidence interval of the VIP > 1.0; * denotes p<0.05 difference (unpaired Student's t-test with Bonferroni correction).

| Region | Control Mean | | SEM | Shifted Mean | | SEM | p-value | Lost/Gained |
|---|---|---|---|---|---|---|---|---|
| *Cortex* | | | | | | | | |
| Amigdalo-Piriform Cortex (APir) | 0,79 | ± | 0,04 | 0,46 | ± | 0,05 | 0,06099 | |
| Cingulate (Cg1) | 0,86 | ± | 0,07 | 0,76 | ± | 0,07 | 158,1914 | |
| Dorsolateral Orbital (DLO) | 0,83 | ± | 0,05 | 0,45 | ± | 0,07 | 0,14756 | |
| Infralimbic (IL) | **1,05** | ± | **0,03** | 0,49* | ± | 0,06 | 4,2E-05* | Lost |
| Lateral Entorhinal (LEC) | 0,77 | ± | 0,09 | 0,52 | ± | 0,05 | 11,14558 | |
| Lateral Orbital (LO) | 0,50 | ± | 0,05 | 0,66 | ± | 0,07 | 40,16005 | |
| Supplementary Motor Area (M2) | 0,68 | ± | 0,08 | 0,50 | ± | 0,07 | 54,66502 | |
| Medial Entorhinal (MEC) | 0,71 | ± | 0,05 | **1,35*** | ± | **0,05** | 7,8E-06* | Gained |
| Medial Orbital (MO) | **1,36** | ± | **0,07** | 0,41* | ± | 0,06 | 1,5E-06* | Lost |
| Medial Prelimbic (mPrL) | 0,74 | ± | 0,04 | 0,43 | ± | 0,08 | 1,666807 | |
| Perirhinal (PRH) | **1,21** | ± | **0,06** | 0,59* | ± | 0,04 | 2,7E-05* | Lost |
| Prelimbic (PrL) | 0,97 | ± | 0,02 | 0,50* | ± | 0,07 | 0,0008* | |
| Retrosplenial (RSC) | **1,20** | ± | **0,05** | 0,98 | ± | 0,07 | 11,55956 | |
| Ventral Orbital (VO) | 0,42 | ± | 0,06 | **1,05*** | ± | **0,09** | 0,0072* | Gained |
| *Hippocampus* | | | | | | | | |
| Dorsal CA1(dCA1) | 0,57 | ± | 0,05 | **1,30*** | ± | **0,08** | 1E-04* | Gained |
| Dorsal CA2 (dCA2) | **1,49** | ± | **0,04** | **2,00*** | ± | **0,04** | 5,2E-06* | Increased |
| Dorsal CA3 (dCA3) | 0,85 | ± | 0,07 | **1,79*** | ± | **0,06** | 8,8E-07* | Gained |
| Dorsal Dentate Gyrus - Rostral (rdDG) | 0,57 | ± | 0,10 | **1,85*** | ± | **0,04** | 5E-08* | Gained |
| Intermediate Dentate Gyrus (iDG) | 0,68 | ± | 0,05 | 0,52 | ± | 0,06 | 19,94158 | |
| Dorsal Hippocampus Molecular Layer (dML) | **1,51** | ± | **0,06** | **1,52** | ± | **0,07** | 415,2585 | |
| Dorsal Subiculum (dSub) | 0,65 | ± | 0,05 | 0,47 | ± | 0,06 | 14,36856 | |
| Intermediate CA1 (iCA1) | 0,85 | ± | 0,07 | 0,41 | ± | 0,06 | 0,07533 | |
| Ventral CA1 (vCA1) | 0,67 | ± | 0,07 | 0,58 | ± | 0,07 | 182,3806 | |
| Ventral CA2 (vCA2) | 0,67 | ± | 0,05 | 0,49 | ± | 0,07 | 21,44194 | |
| Ventral CA3 (vCA3) | 0,65 | ± | 0,05 | 0,52 | ± | 0,06 | 55,61236 | |
| Ventral Hippocampus Molecular Layer (vML) | 0,79 | ± | 0,04 | 0,31* | ± | 0,05 | 0,0002* | |
| *Circadian Rhythms* | | | | | | | | |
| Anterior Hypothalamus (AHA) | 0,61 | ± | 0,05 | **1,30*** | ± | **0,04** | 1,6E-06* | Gained |
| Arcuate Nucleus (ARC) | 0,84 | ± | 0,04 | **1,11** | ± | **0,10** | 8,82679 | |
| Suprachiasmatic Nucleus - Rostral (rSCN) | 0,65 | ± | 0,06 | **1,36*** | ± | **0,05** | 2,6E-06* | Gained |
| Suprachiasmatic Nucleus - Caudal (cSCN) | 0,84 | ± | 0,06 | **1,56*** | ± | **0,09** | 0,0006* | Gained |
| Dorsomedial Hypothalamus (DMH) | 0,81 | ± | 0,04 | 1,00 | ± | 0,09 | 32,55157 | |
| Intergeniculate Leaftlet (IGL) | 0,72 | ± | 0,05 | 0,50 | ± | 0,06 | 6,990504 | |
| Lateral Hypothalamus (LH) | 0,70 | ± | 0,05 | **1,65*** | ± | **0,03** | 2,3E-10* | Gained |
| Medial Preoptic Nucleus - Caudal (cMPO) | 0,97 | ± | 0,07 | 1,01 | ± | 0,05 | 332,1872 | |
| Medial Preoptic Nucleus - Rostral (rMPO) | 0,60 | ± | 0,04 | 0,92 | ± | 0,13 | 13,21387 | |
| Median Eminence (ME) | 0,99 | ± | 0,04 | **1,87*** | ± | **0,07** | 2,4E-07* | Gained |
| Paraventricular Hypothalamic Anterior - Parvicellular (PVH) | 0,51 | ± | 0,04 | 0,42 | ± | 0,06 | 95,02289 | |
| Paraventricular Thalamic Nucleus (PVT) | **1,37** | ± | **0,04** | **1,65** | ± | **0,08** | 1,847099 | |
| Pineal Gland | 0,57 | ± | 0,05 | 0,61 | ± | 0,08 | 331,7611 | |
| Retrochiasmatic Area (ReCh) | 0,53 | ± | 0,04 | 0,81 | ± | 0,08 | 1,909965 | |
| Subparaventricular Zone (SPZ) | 0,38 | ± | 0,03 | **1,24*** | ± | **0,08** | 9,6E-07* | Gained |
| Supraoptic Nucleus (SO) | **1,02** | ± | **0,05** | 0,86 | ± | 0,08 | 56,92811 | |
| Ventrolateral Preoptic Nucleus (VLPO) | 0,93 | ± | 0,04 | 0,62 | ± | 0,14 | 22,7751 | |
| Ventromedial Hypothalamus (VMH) | 0,62 | ± | 0,04 | **1,07** | ± | **0,09** | 0,134445 | |
| Ventromedial Preoptic Nucleus (VMPO) | **1,33** | ± | **0,04** | 0,48* | ± | 0,12 | 0,0010* | Lost |
| *Thalamus* | | | | | | | | |
| Mediodorsal (MD) | **1,71** | ± | **0,13** | **1,90** | ± | **0,08** | 106,6233 | |
| Reuniens (RE) | 0,71 | ± | 0,06 | 0,58 | ± | 0,05 | 38,49561 | |
| *Amygdala* | | | | | | | | |
| Basolateral (BLA) | **1,45** | ± | **0,10** | 0,77* | ± | 0,07 | 0,0075* | Lost |
| Central (CA) | 0,94 | ± | 0,04 | 0,42* | ± | 0,06 | 0,0002* | |
| Medial (MA) | 0,84 | ± | 0,06 | 0,42* | ± | 0,05 | 0,0061* | |
| *Septum* | | | | | | | | |
| Lateral Septum (LS) | **1,55** | ± | **0,04** | 0,97* | ± | 0,10 | 0,0174* | Lost |
| Medial Septum (MS) | **1,52** | ± | **0,09** | 0,75* | ± | 0,04 | 5,3E-05* | Lost |
| *Diagonal Band of Broca* | | | | | | | | |
| Horizontal Band (HDB) | **1,33** | ± | **0,07** | **1,34** | ± | **0,14** | 464,6444 | |
| Vertical Band (VDB) | **1,81** | ± | **0,05** | **1,45** | ± | **0,08** | 0,541108 | |
| *Basal Ganglia* | | | | | | | | |
| Dorsolateral Striatum (DLST) | **1,31** | ± | **0,11** | 0,60* | ± | 0,06 | 0,0081* | Lost |
| Ventromedial Striatum (VMST) | **1,14** | ± | **0,09** | 0,55* | ± | 0,07 | 0,0263* | Lost |
| Substantia Nigra pars compacta (SNC) | 0,71 | ± | 0,04 | 0,42 | ± | 0,05 | 0,14086 | |
| Substantia Nigra pars reticulata (SNR) | 0,71 | ± | 0,05 | 0,68 | ± | 0,13 | 411,4401 | |
| *Mesolimbic* | | | | | | | | |
| Bed Nucleus of the Stria Terminalis (BST) | 0,76 | ± | 0,05 | **1,37*** | ± | **0,09** | 0,0026* | Gained |
| Nucleus Accumbens core (AcbC) | **1,36** | ± | **0,03** | 0,65* | ± | 0,06 | 2,1E-06* | Lost |
| Nucleus Accumbens shell (AcbS) | **1,26** | ± | **0,05** | 0,89 | ± | 0,06 | 0,05477 | |
| *Multimodal* | | | | | | | | |
| Dorsal Raphé (DR) | 0,76 | ± | 0,02 | 0,52 | ± | 0,09 | 7,534984 | |
| Dorsal Tegmental Nucleus (DTg) | 0,86 | ± | 0,06 | 0,52 | ± | 0,09 | 2,628154 | |
| Lateral Habenula (LHab) | **1,49** | ± | **0,04** | 0,55* | ± | 0,15 | 0,0034* | Lost |
| Locus Coeruleus (LC) | 0,70 | ± | 0,04 | 0,54 | ± | 0,07 | 25,31947 | |
| Median Raphé (MR) | **1,22** | ± | **0,03** | 0,54* | ± | 0,07 | 1E-05* | Lost |
| Periaqueductal Gray Matter (PAG) | 0,68 | ± | 0,05 | 0,45 | ± | 0,05 | 1,906182 | |

**Supplementary Table 6– Functional Connectivity of the Ventral Posterolateral Thalamus.** Bold denotes 95% confidence interval of the VIP > 1.0; * denotes p<0.05 difference (unpaired Student's t-test with Bonferroni correction).

| Region | Control | | | Shifted | | | p-value | Lost/Gained |
|---|---|---|---|---|---|---|---|---|
| | Mean | | SEM | Mean | | SEM | | |
| *Cortex* | | | | | | | | |
| Amigdalo-Piriform Cortex (APir) | 0,96 | ± | 0,02 | **1,22*** | ± | **0,03** | 3E-05* | Gained |
| Cingulate (Cg1) | **1,69** | ± | **0,08** | **1,11*** | ± | **0,03** | 0,0006* | Decreased |
| Dorsolateral Orbital (DLO) | 0,73 | ± | 0,06 | 0,95 | ± | 0,02 | 1,200836 | |
| Infralimbic (IL) | **1,14** | ± | **0,05** | **1,16** | ± | **0,01** | 345,2196 | |
| Lateral Entorhinal (LEC) | **1,05** | ± | **0,01** | **1,08** | ± | **0,04** | 219,1273 | |
| Lateral Orbital (LO) | 0,46 | ± | 0,05 | **1,09*** | ± | **0,05** | 5,2E-06* | Gained |
| Supplementary Motor Area (M2) | 0,72 | ± | 0,08 | **1,49*** | ± | **0,03** | 4,6E-06* | Gained |
| Medial Entorhinal (MEC) | **1,33** | ± | **0,04** | 0,95* | ± | 0,04 | 0,0008* | Lost |
| Medial Orbital (MO) | 0,89 | ± | 0,07 | **1,61*** | ± | **0,02** | 3,2E-06* | Gained |
| Medial Prelimbic (mPrL) | **1,16** | ± | **0,06** | 1,00 | ± | 0,02 | 14,0793 | |
| Perirhinal (PRH) | 0,74 | ± | 0,05 | 0,96 | ± | 0,03 | 1,182481 | |
| Prelimbic (PrL) | **1,13** | ± | **0,06** | **1,38** | ± | **0,02** | 0,463944 | |
| Retrosplenial (RSC) | 0,96 | ± | 0,05 | 0,64 | ± | 0,07 | 0,417808 | |
| Ventral Orbital (VO) | **1,71** | ± | **0,11** | **1,04*** | ± | **0,04** | 0,0046* | Decreased |
| *Hippocampus* | | | | | | | | |
| Dorsal CA1(dCA1) | 0,79 | ± | 0,06 | 0,46 | ± | 0,04 | 0,066675 | |
| Dorsal CA2 (dCA2) | **1,06** | ± | **0,07** | 0,65 | ± | 0,07 | 0,277738 | |
| Dorsal CA3 (dCA3) | 0,81 | ± | 0,05 | 0,80 | ± | 0,08 | 433,2633 | |
| Dorsal Dentate Gyrus - Rostral (rdDG) | 0,86 | ± | 0,05 | 0,98 | ± | 0,07 | 89,7224 | |
| Intermediate Dentate Gyrus (iDG) | **1,10** | ± | **0,01** | **1,22** | ± | **0,03** | 0,134317 | |
| Dorsal Hippocampus Molecular Layer (dML) | 0,85 | ± | 0,03 | 0,76 | ± | 0,06 | 78,0664 | |
| Dorsal Subiculum (dSub) | 0,85 | ± | 0,03 | **1,28*** | ± | **0,04** | 1,1E-05* | Gained |
| Intermediate CA1 (iCA1) | 0,85 | ± | 0,04 | **1,47*** | ± | **0,03** | 4,8E-08* | Gained |
| Ventral CA1 (vCA1) | **1,18** | ± | **0,03** | **1,38*** | ± | **0,03** | 0,0448* | Increased |
| Ventral CA2 (vCA2) | **1,01** | ± | **0,02** | **1,28*** | ± | **0,02** | 5,1E-06* | Increased |
| Ventral CA3 (vCA3) | **1,12** | ± | **0,03** | **1,36*** | ± | **0,02** | 0,0012* | Increased |
| Ventral Hippocampus Molecular Layer (vML) | 1,01 | ± | 0,03 | **1,16** | ± | **0,02** | 0,141254 | |
| *Circadian Rhythms* | | | | | | | | |
| Anterior Hypothalamus AHA | 0,30 | ± | 0,04 | **1,41*** | ± | **0,04** | 3,5E-11* | Gained |
| Arcuate Nucleus (ARC) | 0,96 | ± | 0,03 | 0,38* | ± | 0,04 | 1,5E-07* | |
| Suprachiasmatic Nucleus - Rostral (rSCN) | 0,48 | ± | 0,09 | 0,30 | ± | 0,04 | 42,14014 | |
| Suprachiasmatic Nucleus - Caudal (cSCN) | 0,52 | ± | 0,07 | **1,36*** | ± | **0,06** | 6E-06* | Gained |
| Dorsomedial Hypothalamus (DMH) | **1,06** | ± | **0,04** | 0,60* | ± | 0,05 | 0,0002* | Lost |
| Intergeniculate Leaftlet (IGL) | **1,33** | ± | **0,03** | **1,07*** | ± | **0,02** | 0,0002* | Decreased |
| Lateral Hypothalamus (LH) | 0,77 | ± | 0,09 | 0,95 | ± | 0,07 | 64,38911 | |
| Medial Preoptic Nucleus - Caudal (cMPO) | 0,57 | ± | 0,05 | **1,07*** | ± | **0,02** | 1,4E-05* | Gained |
| Medial Preoptic Nucleus - Rostral (rMPO) | 0,78 | ± | 0,06 | 0,49 | ± | 0,04 | 0,280025 | |
| Median Eminence (ME) | 0,90 | ± | 0,03 | 0,47* | ± | 0,05 | 0,0001* | |
| Paraventricular Hypothalamic Anterior - Parvicellular (PVH) | 0,71 | ± | 0,08 | 0,78 | ± | 0,08 | 264,1665 | |
| Paraventricular Thalamic Nucleus (PVT) | **1,17** | ± | **0,07** | 0,63* | ± | 0,03 | 0,0005* | Lost |
| Pineal Gland | 0,85 | ± | 0,02 | 0,90 | ± | 0,07 | 268,2402 | |
| Retrochiasmatic Area (ReCh) | **1,54** | ± | **0,07** | 0,79* | ± | 0,08 | 0,0003* | Lost |
| Subparaventricular Zone (SPZ) | 0,62 | ± | 0,04 | **1,04*** | ± | **0,04** | 0,0003* | Gained |
| Supraoptic Nucleus (SO) | 0,53 | ± | 0,06 | 0,92* | ± | 0,03 | 0,0025* | |
| Ventrolateral Preoptic Nucleus (VLPO) | **1,12** | ± | **0,03** | 0,63* | ± | 0,04 | 4E-06* | Lost |
| Ventromedial Hypothalamus (VMH) | 0,93 | ± | 0,07 | 0,67 | ± | 0,06 | 5,780179 | |
| Ventromedial Preoptic Nucleus (VMPO) | 1,00 | ± | 0,06 | 0,82 | ± | 0,04 | 12,0436 | |
| *Thalamus* | | | | | | | | |
| Mediodorsal (MD) | 0,95 | ± | 0,11 | 0,86 | ± | 0,04 | 202,8197 | |
| Reuniens (RE) | 0,84 | ± | 0,05 | 0,95 | ± | 0,03 | 46,90857 | |
| Ventral Posterolateral (VPL) | 0,73 | ± | 0,04 | 0,49 | ± | 0,08 | 6,032183 | |
| *Amygdala* | | | | | | | | |
| Basolateral (BLA) | 0,69 | ± | 0,05 | **1,32*** | ± | **0,07** | 0,0003* | Gained |
| Central (CA) | 0,96 | ± | 0,03 | **1,22** | ± | **0,06** | 0,390442 | |
| Medial (MA) | **1,04** | ± | **0,04** | 1,00 | ± | 0,02 | 184,0655 | |
| *Septum* | | | | | | | | |
| Lateral Septum (LS) | **1,03** | ± | **0,06** | 0,88 | ± | 0,05 | 32,37286 | |
| Medial Septum (MS) | **1,05** | ± | **0,08** | 0,63 | ± | 0,04 | 0,061947 | |
| *Diagonal Band of Broca* | | | | | | | | |
| Horizontal Band (HDB) | 0,67 | ± | 0,09 | 0,62 | ± | 0,07 | 309,7224 | |
| Vertical Band (VDB) | 0,78 | ± | 0,06 | 0,59 | ± | 0,06 | 12,42687 | |
| *Basal Ganglia* | | | | | | | | |
| Dorsolateral Striatum (DLST) | 0,47 | ± | 0,06 | 0,94* | ± | 0,07 | 0,0141* | |
| Ventromedial Striatum (VMST) | **1,21** | ± | **0,06** | 0,85 | ± | 0,05 | 0,12686 | |
| Substantia Nigra pars compacta (SNC) | **1,14** | ± | **0,01** | **1,60*** | ± | **0,03** | 7E-09* | Increased |
| Substantia Nigra pars reticulata (SNR) | **1,11** | ± | **0,01** | **1,04** | ± | **0,06** | 134,6818 | |
| *Mesolimbic* | | | | | | | | |
| Bed Nucleus of the Stria Terminalis (BST) | **1,54** | ± | **0,08** | 0,64* | ± | 0,03 | 3,7E-07* | Lost |
| Nucleus Accumbens core (AcbC) | **1,19** | ± | **0,05** | 0,65* | ± | 0,03 | 1,1E-05* | Lost |
| Nucleus Accumbens shell (AcbS) | **1,01** | ± | **0,03** | 0,57* | ± | 0,03 | 4,3E-06* | Lost |
| *Multimodal* | | | | | | | | |
| Dorsal Tegmental Nucleus (DTg) | 0,51 | ± | 0,04 | 0,95 | ± | 0,10 | 0,325293 | |
| Lateral Habenula (LHab) | 0,55 | ± | 0,07 | 0,61 | ± | 0,06 | 249,2176 | |
| Locus Coeruleus (LC) | **1,28** | ± | **0,04** | **1,18** | ± | **0,06** | 90,24178 | |
| Median Raphé (MR) | 0,74 | ± | 0,03 | 1,02 | ± | 0,08 | 1,741856 | |
| Periaqueductal Gray Matter (PAG) | **1,20** | ± | **0,02** | 0,91* | ± | 0,06 | 0,0380* | Lost |

**Supplementary Table 7– Functional Connectivity of the Dorsal Raphé**

Bold denotes 95% confidence interval of the VIP > 1.0; * denotes p<0.05 difference (unpaired Student's t-test with Bonferroni correction).

| Region | Control Mean | | Control SEM | Shifted Mean | | Shifted SEM | p-value | Lost/Gained |
|---|---|---|---|---|---|---|---|---|
| *Cortex* | | | | | | | | |
| Amigdalo-Piriform Cortex (APir) | 0,78 | ± | 0,05 | **1,42*** | ± | **0,03** | 3,2E-07* | **Gained** |
| Cingulate (Cg1) | 0,45 | ± | 0,03 | **1,62*** | ± | **0,08** | 4,2E-09* | **Gained** |
| Dorsolateral Orbital (DLO) | **1,13** | ± | **0,06** | 0,64 | ± | 0,10 | 0,278039 | |
| Infralimbic (IL) | 0,74 | ± | 0,06 | 0,61 | ± | 0,06 | 61,12196 | |
| Lateral Entorhinal (LEC) | 0,54 | ± | 0,07 | **1,51*** | ± | **0,03** | 2,4E-08* | **Gained** |
| Lateral Orbital (LO) | **1,37** | ± | **0,06** | **1,57** | ± | **0,05** | 7,624523 | |
| Supplementary Motor Area (M2) | 0,45 | ± | 0,10 | 0,76 | ± | 0,02 | 3,667758 | |
| Medial Entorhinal (MEC) | 0,62 | ± | 0,05 | 0,45 | ± | 0,02 | 2,471894 | |
| Medial Orbital (MO) | 0,77 | ± | 0,08 | 0,83 | ± | 0,02 | 242,7231 | |
| Medial Prelimbic (mPrL) | 0,55 | ± | 0,05 | 0,64 | ± | 0,04 | 80,8741 | |
| Perirhinal (PRH) | 0,96 | ± | 0,10 | **1,01** | ± | **0,03** | 318,7428 | |
| Prelimbic (PrL) | **1,15** | ± | **0,07** | 0,54* | ± | 0,07 | 0,0039* | **Lost** |
| Retrosplenial (RSC) | 0,65 | ± | 0,05 | **1,13*** | ± | **0,05** | 0,0004* | **Gained** |
| Ventral Orbital (VO) | 0,92 | ± | 0,07 | 0,50* | ± | 0,04 | 0,0098* | |
| *Hippocampus* | | | | | | | | |
| Dorsal CA1(dCA1) | 0,59 | ± | 0,06 | 0,70 | ± | 0,08 | 126,2646 | |
| Dorsal CA2 (dCA2) | **1,22** | ± | **0,06** | 0,83* | ± | 0,05 | 0,0487* | **Lost** |
| Dorsal CA3 (dCA3) | 0,53 | ± | 0,04 | 0,65 | ± | 0,03 | 11,09526 | |
| Dorsal Dentate Gyrus - Rostral (rdDG) | 0,54 | ± | 0,05 | 0,64 | ± | 0,06 | 100,6566 | |
| Intermediate Dentate Gyrus (iDG) | 0,63 | ± | 0,03 | 0,67 | ± | 0,02 | 165,0692 | |
| Dorsal Hippocampus Molecular Layer (dML) | **1,28** | ± | **0,06** | 0,46* | ± | 0,08 | 3,8E-05* | **Lost** |
| Dorsal Subiculum (dSub) | 0,86 | ± | 0,04 | 0,67 | ± | 0,07 | 12,60266 | |
| Intermediate CA1 (iCA1) | **1,17** | ± | **0,06** | 0,53* | ± | 0,03 | 2E-06* | **Lost** |
| Ventral CA1 (vCA1) | 0,58 | ± | 0,06 | 0,61 | ± | 0,06 | 373,0141 | |
| Ventral CA2 (vCA2) | 0,67 | ± | 0,04 | 0,62 | ± | 0,06 | 252,8421 | |
| Ventral CA3 (vCA3) | 0,74 | ± | 0,03 | 0,64 | ± | 0,03 | 23,33498 | |
| Ventral Hippocampus Molecular Layer (vML) | 0,86 | ± | 0,06 | 0,57 | ± | 0,06 | 1,197215 | |
| *Circadian Rhythms* | | | | | | | | |
| Anterior Hypothalamus AHA | 0,57 | ± | 0,05 | **1,14*** | ± | **0,05** | 5,6E-05* | **Gained** |
| Arcuate Nucleus (ARC) | 0,79 | ± | 0,05 | **1,11** | ± | **0,06** | 0,149277 | |
| Suprachiasmatic Nucleus - Rostral (rSCN) | 0,94 | ± | 0,07 | 0,35* | ± | 0,06 | 0,0012* | |
| Suprachiasmatic Nucleus - Caudal (cSCN) | **1,18** | ± | **0,07** | 0,50* | ± | 0,03 | 2,6E-05* | **Lost** |
| Dorsomedial Hypothalamus (DMH) | 0,57 | ± | 0,04 | 0,84 | ± | 0,06 | 0,384328 | |
| Intergeniculate Leaftlet (IGL) | 0,63 | ± | 0,06 | **1,67*** | ± | **0,03** | 4,1E-10* | **Gained** |
| Lateral Hypothalamus (LH) | **1,38** | ± | **0,04** | **1,87*** | ± | **0,07** | 0,0043* | **Increased** |
| Medial Preoptic Nucleus - Caudal (cMPO) | 0,48 | ± | 0,08 | 0,97* | ± | 0,03 | 0,0139* | |
| Medial Preoptic Nucleus - Rostral (rMPO) | 0,33 | ± | 0,03 | 0,70* | ± | 0,04 | 0,0005* | |
| Median Eminence (ME) | 0,98 | ± | 0,06 | 0,69 | ± | 0,05 | 0,931756 | |
| Paraventricular Hypothalamic Anterior - Parvicellular (PVH) | 0,65 | ± | 0,07 | **1,03** | ± | **0,07** | 0,37176 | |
| Paraventricular Thalamic Nucleus (PVT) | **1,38** | ± | **0,07** | **1,04** | ± | **0,05** | 0,251598 | |
| Pineal Gland | 1,02 | ± | 0,08 | **1,08** | ± | **0,06** | 284,144 | |
| Retrochiasmatic Area (ReCh) | 0,60 | ± | 0,04 | **1,74*** | ± | **0,07** | 5,1E-09* | **Gained** |
| Subparaventricular Zone (SPZ) | 0,37 | ± | 0,04 | 0,45 | ± | 0,04 | 81,10023 | |
| Supraoptic Nucleus (SO) | **1,52** | ± | **0,06** | **1,11** | ± | **0,06** | 0,067885 | |
| Ventrolateral Preoptic Nucleus (VLPO) | 0,73 | ± | 0,05 | **1,62*** | ± | **0,06** | 4,7E-08* | **Gained** |
| Ventromedial Hypothalamus (VMH) | 0,38 | ± | 0,05 | 0,78* | ± | 0,05 | 0,0055* | |
| Ventromedial Preoptic Nucleus (VMPO) | **1,03** | ± | **0,08** | 1,00 | ± | 0,05 | 353,5119 | |
| *Thalamus* | | | | | | | | |
| Mediodorsal (MD) | **1,58** | ± | **0,06** | **1,10*** | ± | **0,05** | 0,0038* | **Decreased** |
| Reuniens (RE) | 0,54 | ± | 0,03 | **1,04*** | ± | **0,03** | 3,2E-08* | **Gained** |
| Ventral Posterolateral (VPL) | **1,55** | ± | **0,07** | 0,72* | ± | 0,06 | 1,3E-05* | **Lost** |
| *Amygdala* | | | | | | | | |
| Basolateral (BLA) | 0,91 | ± | 0,07 | 0,62 | ± | 0,04 | 0,727886 | |
| Central (CA) | 0,77 | ± | 0,05 | **1,89*** | ± | **0,05** | 1,2E-09* | **Gained** |
| Medial (MA) | 0,50 | ± | 0,07 | **1,35*** | ± | **0,05** | 3,6E-06* | **Gained** |
| *Septum* | | | | | | | | |
| Lateral Septum (LS) | **1,22** | ± | **0,05** | 0,60* | ± | 0,04 | 1,9E-06* | **Lost** |
| *Diagonal Band of Broca* | | | | | | | | |
| Horizontal Band (HDB) | **1,60** | ± | **0,04** | 0,45* | ± | 0,04 | 3,9E-12* | **Lost** |
| Vertical Band (VDB) | **1,63** | ± | **0,06** | **2,42*** | ± | **0,06** | 5,3E-06* | **Increased** |
| *Basal Ganglia* | | | | | | | | |
| Dorsolateral Striatum (DLST) | **1,71** | ± | **0,04** | **1,63** | ± | **0,04** | 66,87423 | |
| Ventromedial Striatum (VMST) | **1,24** | ± | **0,07** | 0,75* | ± | 0,04 | 0,0043* | **Lost** |
| Substantia Nigra pars compacta (SNC) | 0,49 | ± | 0,03 | 0,59 | ± | 0,06 | 69,32014 | |
| Substantia Nigra pars reticulata (SNR) | 0,57 | ± | 0,03 | 0,92* | ± | 0,05 | 0,0094* | |
| *Mesolimbic* | | | | | | | | |
| Bed Nucleus of the Stria Terminalis (BST) | 0,93 | ± | 0,06 | 0,74 | ± | 0,11 | 77,56037 | |
| Nucleus Accumbens core (AcbC) | **1,18** | ± | **0,04** | 0,74* | ± | 0,07 | 0,0068* | **Lost** |
| Nucleus Accumbens shell (AcbS) | **1,24** | ± | **0,04** | 0,48* | ± | 0,12 | 0,0052* | **Lost** |
| *Multimodal* | | | | | | | | |
| Dorsal Raphé (DR) | 0,89 | ± | 0,07 | 0,56 | ± | 0,03 | 0,124244 | |
| Dorsal Tegmental Nucleus (DTg) | **1,15** | ± | **0,06** | 0,49* | ± | 0,03 | 5,9E-07* | **Lost** |
| Lateral Habenula (LHab) | **1,43** | ± | **0,07** | 0,57* | ± | 0,05 | 5,2E-07* | **Lost** |
| Locus Coeruleus (LC) | **1,23** | ± | **0,05** | 0,54* | ± | 0,03 | 1,8E-08* | **Lost** |
| Median Raphé (MR) | **1,38** | ± | **0,05** | 0,61* | ± | 0,03 | 1,1E-08* | **Lost** |
| Periaqueductal Gray Matter (PAG) | 0,67 | ± | 0,04 | 0,52 | ± | 0,04 | 9,782086 | |

**Supplementary Table 8 – Functional Connectivity of the Medial Septum** Bold denotes 95% confidence interval of the VIP > 1.0; * denotes p<0.05 difference (unpaired Student's t-test with Bonferroni correction).

| | Control | | | Shifted | | | | |
|---|---|---|---|---|---|---|---|---|
| **Region** | **Mean** | | **SEM** | **Mean** | | **SEM** | **p-value** | **Lost/Gained** |
| *Cortex* | | | | | | | | |
| Amigdalo-Piriform Cortex (APir) | **1,20** | ± | **0,04** | 0,78* | ± | 0,05 | 0,0013* | Lost |
| Cingulate (Cg1) | **1,07** | ± | **0,05** | **1,44** | ± | **0,07** | 0,1620 | |
| Dorsolateral Orbital (DLO) | **1,39** | ± | **0,05** | **1,21** | ± | **0,05** | 9,8058 | |
| Infralimbic (IL) | 0,77 | ± | 0,06 | **1,01** | ± | **0,02** | 0,3790 | |
| Lateral Entorhinal (LEC) | **1,13** | ± | **0,03** | 0,80 | ± | 0,06 | 0,0732 | |
| Lateral Orbital (LO) | 0,53 | ± | 0,06 | 0,82 | ± | 0,08 | 4,7172 | |
| Supplementary Motor Area (M2) | **1,84** | ± | **0,05** | 0,59* | ± | 0,06 | 3,1E-10* | Lost |
| Medial Entorhinal (MEC) | 0,81 | ± | 0,02 | **1,10*** | ± | **0,05** | 0,0212* | Gained |
| Medial Orbital (MO) | **1,65** | ± | **0,06** | 0,74* | ± | 0,04 | 1,3E-08* | Lost |
| Medial Prelimbic (mPrL) | 0,74 | ± | 0,05 | **1,58*** | ± | **0,05** | 3,8E-08* | Gained |
| Perirhinal (PRH) | 0,62 | ± | 0,08 | 0,88 | ± | 0,05 | 3,9599 | |
| Prelimbic (PrL) | 0,70 | ± | 0,03 | 0,94 | ± | 0,06 | 1,0075 | |
| Retrosplenial (RSC) | 0,71 | ± | 0,06 | 0,40 | ± | 0,06 | 1,1465 | |
| Ventral Orbital (VO) | **1,36** | ± | **0,03** | 0,63* | ± | 0,05 | 4,5E-08* | Lost |
| *Hippocampus* | | | | | | | | |
| Dorsal CA1(dCA1) | 0,61 | ± | 0,03 | 0,76 | ± | 0,06 | 22,4124 | |
| Dorsal CA2 (dCA2) | 0,69 | ± | 0,05 | 0,74 | ± | 0,07 | 277,9045 | |
| Dorsal CA3 (dCA3) | 0,63 | ± | 0,11 | 0,38 | ± | 0,06 | 27,7908 | |
| Dorsal Dentate Gyrus - Rostral (rdDG) | 1,01 | ± | 0,06 | 0,89 | ± | 0,05 | 69,4914 | |
| Intermediate Dentate Gyrus (iDG) | 1,01 | ± | 0,05 | **1,09** | ± | **0,07** | 155,3840 | |
| Dorsal Hippocampus Molecular Layer (dML) | 0,62 | ± | 0,04 | 0,95 | ± | 0,09 | 1,3778 | |
| Dorsal Subiculum (dSub) | 0,56 | ± | 0,06 | 0,66 | ± | 0,08 | 165,6038 | |
| Intermediate CA1 (iCA1) | 1,03 | ± | 0,06 | 0,75 | ± | 0,05 | 0,9152 | |
| Ventral CA1 (vCA1) | **1,27** | ± | **0,03** | 0,68* | ± | 0,10 | 0,0082* | Lost |
| Ventral CA2 (vCA2) | **1,22** | ± | **0,04** | 0,67* | ± | 0,08 | 0,0011* | Lost |
| Ventral CA3 (vCA3) | 0,88 | ± | 0,06 | 0,85 | ± | 0,06 | 365,6435 | |
| Ventral Hippocampus Molecular Layer (vML) | **1,17** | ± | **0,05** | 0,85 | ± | 0,09 | 2,2723 | |
| *Circadian Rhythms* | | | | | | | | |
| Anterior Hypothalamus AHA | **1,43** | ± | **0,06** | 0,92* | ± | 0,08 | 0,0209* | Lost |
| Arcuate Nucleus (ARC) | 0,84 | ± | 0,03 | 0,65 | ± | 0,08 | 15,21218 | |
| Suprachiasmatic Nucleus - Rostral (rSCN) | 0,58 | ± | 0,05 | **1,68*** | ± | **0,09** | 2,8E-07* | Gained |
| Suprachiasmatic Nucleus - Caudal (cSCN) | 0,75 | ± | 0,06 | **1,27*** | ± | **0,05** | 0,0006* | Gained |
| Dorsomedial Hypothalamus (DMH) | 0,80 | ± | 0,05 | 0,94 | ± | 0,04 | 18,21334 | |
| Intergeniculate Leaftlet (IGL) | **1,40** | ± | **0,02** | 0,52* | ± | 0,06 | 1,6E-09* | Lost |
| Lateral Hypothalamus (LH) | 0,69 | ± | 0,08 | 0,49 | ± | 0,06 | 25,68397 | |
| Medial Preoptic Nucleus - Caudal (cMPO) | **1,41** | ± | **0,08** | 0,66* | ± | 0,08 | 0,00063* | Lost |
| Medial Preoptic Nucleus - Rostral (rMPO) | 0,70 | ± | 0,04 | **1,41*** | ± | **0,06** | 4,5E-06* | Gained |
| Median Eminence (ME) | **1,27** | ± | **0,03** | 0,64* | ± | 0,07 | 8,4E-05* | Lost |
| Paraventricular Thalamic Nucleus (PVT) | 0,57 | ± | 0,03 | 0,69 | ± | 0,06 | 54,94755 | |
| Pineal Gland | 0,47 | ± | 0,04 | **1,49*** | ± | **0,08** | 9,1E-08* | Gained |
| Retrochiasmatic Area (ReCh) | 0,84 | ± | 0,09 | **1,26** | ± | **0,07** | 0,914779 | |
| Subparaventricular Zone (SPZ) | **1,33** | ± | **0,04** | **1,80*** | ± | **0,04** | 0,00013* | Increased |
| Supraoptic Nucleus (SO) | 0,70 | ± | 0,07 | 0,71 | ± | 0,05 | 412,2315 | |
| Ventrolateral Preoptic Nucleus (VLPO) | 0,90 | ± | 0,05 | **1,23** | ± | **0,06** | **0,104061** | |
| Ventromedial Hypothalamus (VMH) | 0,81 | ± | 0,04 | 0,85 | ± | 0,09 | 340,8404 | |
| Ventromedial Preoptic Nucleus (VMPO) | **1,34** | ± | **0,04** | 0,87* | ± | 0,02 | 1E-06* | Lost |
| *Thalamus* | | | | | | | | |
| Mediodorsal (MD) | 0,59 | ± | 0,06 | 1,01 | ± | 0,07 | 0,068733 | |
| Reuniens (RE) | **1,33** | ± | **0,05** | 0,61* | ± | 0,09 | 0,00023* | Lost |
| Ventral Posterolateral (VPL) | 0,49 | ± | 0,06 | 0,43 | ± | 0,04 | 192,5895 | |
| *Amygdala* | | | | | | | | |
| Basolateral (BLA) | 0,78 | ± | 0,09 | 0,90 | ± | 0,06 | 124,8995 | |
| Central (CA) | 0,77 | ± | 0,03 | 0,98 | ± | 0,04 | 0,365791 | |
| Medial (MA) | 0,89 | ± | 0,04 | **1,29*** | ± | **0,05** | 0,0011* | Gained |
| *Septum* | | | | | | | | |
| Lateral Septum (LS) | 0,62 | ± | 0,06 | 0,56 | ± | 0,05 | 208,3836 | |
| Medial Septum (MS) | 0,59 | ± | 0,05 | **1,04*** | ± | **0,08** | 0,0464* | Gained |
| *Diagonal Band of Broca* | | | | | | | | |
| Horizontal Band (HDB) | **1,14** | ± | **0,05** | 0,44* | ± | 0,07 | 4,1E-05* | Lost |
| Vertical Band (VDB) | 0,93 | ± | 0,04 | 1,02 | ± | 0,08 | 150,2365 | |
| *Basal Ganglia* | | | | | | | | |
| Dorsolateral Striatum (DLST) | 0,73 | ± | 0,07 | 0,42 | ± | 0,10 | 10,17919 | |
| Ventromedial Striatum (VMST) | **1,12** | ± | **0,05** | 0,72* | ± | 0,04 | 0,0052* | Lost |
| Substantia Nigra pars compacta (SNC) | 0,94 | ± | 0,06 | 0,91 | ± | 0,04 | 297,8425 | |
| Substantia Nigra pars reticulata (SNR) | 1,01 | ± | 0,05 | 0,50* | ± | 0,07 | 0,0020* | Lost |
| *Mesolimbic* | | | | | | | | |
| Bed Nucleus of the Stria Terminalis (BST) | **1,52** | ± | **0,04** | **1,28** | ± | **0,05** | 0,543684 | |
| Nucleus Accumbens core (AcbC) | 0,80 | ± | 0,04 | **1,93*** | ± | **0,07** | 3,7E-09* | Gained |
| Nucleus Accumbens shell (AcbS) | 0,98 | ± | 0,05 | **1,62*** | ± | **0,09** | 0,00232* | Gained |
| *Multimodal* | | | | | | | | |
| Dorsal Raphé (DR) | 0,68 | ± | 0,07 | 0,84 | ± | 0,08 | 59,4325 | |
| Dorsal Tegmental Nucleus (DTg) | 0,58 | ± | 0,07 | 0,99 | ± | 0,08 | 0,360425 | |
| Lateral Habenula (LHab) | 0,56 | ± | 0,05 | **1,41*** | ± | **0,06** | 6,6E-07* | Gained |
| Locus Coeruleus (LC) | **1,16** | ± | **0,05** | 0,56* | ± | 0,07 | 0,00068* | Lost |
| Median Raphé (MR) | **1,04** | ± | **0,04** | **1,16** | ± | **0,07** | 79,84287 | |
| Periaqueductal Gray Matter (PAG) | **1,55** | ± | **0,02** | 0,69* | ± | 0,05 | 1,3E-09* | Lost |

**Supplementary Table 9 – Functional Connectivity of the Periventricular Hypothalamic Nucleus**

Bold denotes 95% confidence interval of the VIP > 1.0; * denotes p<0.05 difference (unpaired Student's t-test with Bonferroni correction)
